# Conjugation related costs have reduced impact on *in silico* plasmid persistence

**DOI:** 10.1101/2024.01.18.576315

**Authors:** Arthur Newbury

## Abstract

Due to the important role they play in the antimicrobial resistance (AMR) crisis and in microbial evolution in general, a great deal of empirical and theoretical work is currently underway, trying to understand plasmid ecology. One of the key questions is how these often costly genetic elements persist in host populations. Here I show that when modelling plasmid population dynamics, it is not sufficient to treat them as always costly (or beneficial). I argue that conjugation related costs may be more important to plasmids in nature than they are in benign laboratory settings. Furthermore, I show that these conjugation related costs can be very severe and still not lead to extinction of a plasmid from a host population.

## Introduction

Plasmids are mobile genetic elements (MGEs) that are major vectors for antimicrobial resistance (AMR), spreading AMR genes between diverse bacterial species within hospitals, allowing pathogenic species to evade treatment (Sheppard et al. 2016; Weingarten et al. 2018; Peter et al. 2020). Furthermore, they often encode virulence factors, enabling pathogens to establish infections and evade host immune responses (Bruto et al. 2017; Pilla and Tang 2018). While AMR plasmids may be beneficial to host bacteria in the presence of antibiotics, plasmids can also be costly (Millan and MacLean 2017), leading to the so called *paradox* (Harrison and Brockhurst 2012) of plasmid persistence in situations where they may be predicted not to survive (Stewart and Levin 1977). In the present work I investigate how predictions about plasmid persistence can change when costs arise in different ways. Specifically, I focus on costs related to conjugation, the mechanism by which plasmids are horizontally transferred between cells.

In laboratory settings, the costs associated with plasmid carriage are often ameliorated or nullified due to compensatory evolution. Compensatory evolution often involves mutation of just one or a small number of genes (Harrison et al. 2015; San Millan et al. 2015; Loftie-Eaton et al. 2017; Hall et al. 2020, 2021), suggesting that plasmid costs typically stem from specific genetic interactions between plasmid and host (Hall et al. 2021). However, in addition to the usual difficulties associated with extrapolating results from experiments growing bacteria in monoculture (or small synthetic communities) in nutrient media to attempt to understand complex natural microbial communities (e.g. costs associated with conjugation become more apparent at lower nutrient levels (Bethke et al. 2023) and plasmid costs are associated with the presence of phage (Jalasvuori et al. 2011)) there are specific methodological challenges to incorporating conjugation related costs into fitness assays. Measuring growth rates of fully infected populations explicitly negates all costs associated with conjugation. Furthermore, in coculture competition assays if all or most conjugation takes place between rather than within strains then the costs associated with conjugation may impact both populations approximately equally.

While costs caused by specific genetic interactions may be nullified by compensatory evolution, metabolically costly behaviour such as conjugation (Turner, Cooper, and Lenski 1998; Ilangovan, Connery, and Waksman 2015) can at best be alleviated e.g. by improved efficiency. Furthermore, formation of the pilus structure (necessary for conjugation) confers a fitness cost to individual bacterial cells outside of laboratory settings since phage target the pilus (Jalasvuori et al. 2011). Since we can expect the loss of genetic interaction based costs in any longstanding associations between plasmids and bacteria in nature, but not necessarily a loss of conjugation related costs, the question asked here is how should the cost of conjugation affect the fate of plasmids within a host population. Pioneering mathematical modelling work by Stewart and Levin applying ecological population modelling techniques to plasmid and host populations (Stewart and Levin 1977) and more recent applications of similar modelling approaches multiple hosts (Alonso-del Valle et al. 2021; Newbury et al. 2022) and multiple plasmids (Risely et al. 2024) assume a cost (or benefit in some cases) applied to the growth function of the sub-population of bacteria which are infected with a plasmid. Within this framework, the ability for a plasmid population to grow from rare and maintain a stable frequency within the host population is determined by it’s impact on host growth rate and the rate of conjugation between hosts, with stronger negative impacts to growth rate requiring higher conjugation rates in order to maintain a plasmid population, that is higher rates of horizontal transmission are needed to balance low rates of vertical transmission of plasmids. However, this simplifying assumption - that the plasmid host incurs a uniform cost across it’s lifespan - may differ sufficiently from reality to yield incorrect predictions about the fate of plasmid populations.

Viewing the plasmid as a fitness maximising agent, we may ask “how much of a cost to future vertical transmission would it be willing to pay in exchange for a single instance of horizontal transmission?”. The exact answer to this will depend on biological details (life history and mortality rates of hosts etc.) but intuitively it makes sense that a plasmid should be willing to pay a higher price on the basis that the price is only paid in *exchange* for reproduction in the form conjugation. Here, I formalise this intuitive idea, first contrasting a simple model of plasmid population dynamics with an alternative model in which the costs incurred by plasmid hosts only occur after conjugation. Next I analyse a more sophisticated model in which the proportion of costs of plasmid carriage which stem from conjugation can be varied.

## Results

We first consider a simple logistic model of bacterial growth with horizontal transmission of plasmids

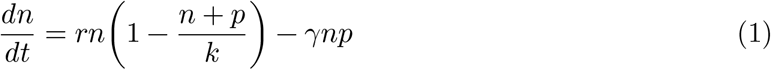

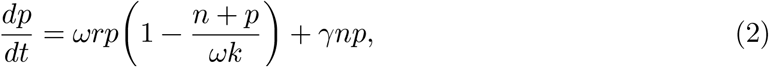

where *n* and *p* represent the sizes of the bacterial sub-populations with and without plasmids respectively, *r* is the hosts intrinsic growth rate, *k* it’s carrying capacity, *γ* is the rate of conjugation and *ω* is a fitness modifier - a number between 0 and 1 by which we multiply both *r* and *k* for infected hosts. This is a textbook model of population growth, with transmission dynamics from a susseptible infiected (SI) model (OTTO and DAY 2007). Here, plasmid can spread from an initially negligible frequency, so long as

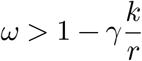

That is - the plasmid is more likely to spread if *ω* is large (low fitness cost or even benefit) and/or *γ* is large (high conjugation rate).

Adapting the predvious model to include a third class *c* for costly plasmid, we now assume the costs *ω* apply only to the growth of hosts when the plasmid is in it’s costly state which it transitions to upon conjugation, and returns from at rate *δ*. This gives the following dynamical equations

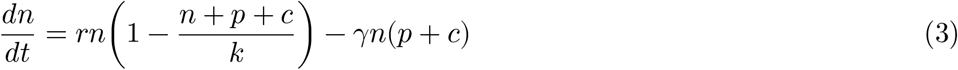

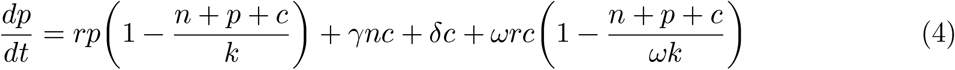

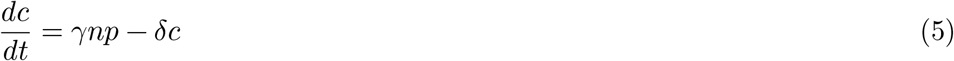

Now the plasmid can always invade the host population as long as *γ* > 0, i.e., whenever there is some conjugation (see methods). This suggests that costly plasmids should be able to persist in bacterial populations, so long as the costs manifest during conjugation.

The previous result comes from a simplified model, which allowed for an instructive analytical solution. However, since it was lacking in biological detail, I investigated the impact of conjugation related costs in a plasmid population dynamics model adapted from Alonso-del Valle et al(2021). In the model, bacterial populations grow according to a saturating function of the concentration of a resource *R*

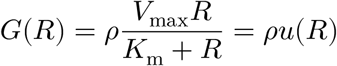

where *V*_max_ is the maximum consumption rate of the bacteria and *K*_m_ is a half-saturation constant. This is equivalent to Equation 3 in Alonso-del Valle et al(2021).

We consider 3 host sub-populations: uninfected *B*_0_, infected *B*_*p*_ and infected having recently conjugated *B*_*c*_. When either *B*_*p*_ or *B*_*c*_ reproduce, they produce mainly *B*_*p*_ offspring, though a proportion *λ* of the offsping will be *B*_0_ due to loss of plasmids during segregation. Bacteria in each sub-population die at a rate *d* multiplied by the total density of all bacteria, i.e., density dependent mortality. Both *B*_*p*_ and *B*_*c*_ sub-populations conjugate with available *B*_0_ at rate *γ*, to produce new cells of *B*_*p*_. Since *B*_*p*_ cells become *B*_*c*_ upon conjugation, this leaves *B*_*p*_ sub-population density unchanged. Additionally, *B*_*c*_ cells recover at rate *δ*. This gives the following dynamical equations:

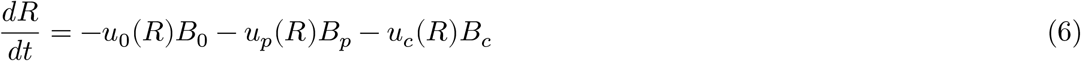

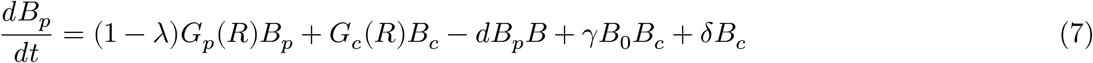

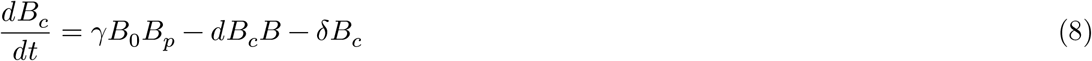

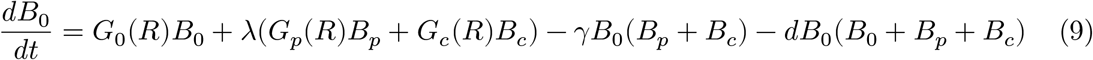

Where the subscripts 0, *p* and *c* denote the growth *G* and consumption *u* functions for *B*_0_, *B*_*p*_ and *B*_*c*_ respectively.

To analyse the effects of plasmid costs on plasmid persistence I numerically solved this system of equations for 100,000 time steps (enough to be confident the system is near equilibrium) for a range of conjugation rates and plasmid costs, and recorded the final density of plasmid bearing cells *B*_*p*_ + *B*_*c*_. The cost of carrying a plasmid was modelled by multiplying both *V*_max_ and *ρ* by *ω*_*p*_ for the growth and consumption of *B*_*p*_ and by *ω*_*p*_*ω*_*c*_ for *B*_*c*_. That is, *ω*_*p*_ is the relative fitness of *B*_*p*_ compared with *B*_0_ and *ω*_*c*_ is the relative fitness of *B*_*c*_ compared with *B*_*p*_.

While the original intention was to investigate the impact of varying *ω*_*c*_, it became apparent that even in the most extreme case of setting *ω*_*c*_ = 0 and *δ* = 0 (so that after conjugation the donor will never produce any offspring cells again) plasmids will persist and even go to fixation in the population, so long as there are no other costs, i.e., *ω*_*p*_ = 1 (Figure 1 b). That is not to say that conjugation costs did not affect the persistence of plasmids in general. We can see that towards the upper end of Figure 1 b we are reaching the lower limit of conjugation rate that can sustain a plasmid population with such extreme conjugation costs. Also in Figure 1 b we can see that the addition of these conjugation costs reduces the range of parameters *ω*_*p*_ and *γ* that allow plasmids to persist. Qualitatively similar patterns can be seen for a range of parameter values in the supplementary materials.

**Figure 1:**
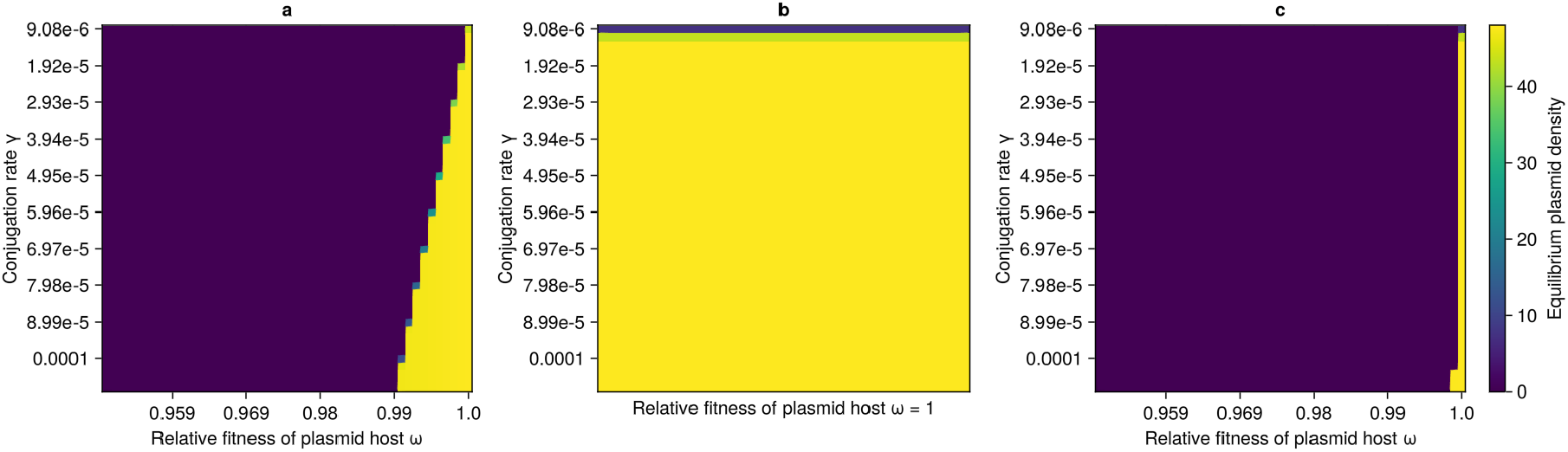
Density of plasmid bearing cells at equilibrium when a) plasmid costs do not increase with conjugation; b) plasmids do not impose any cost until conjugation, at which point the donor loses all reproductive fitness; c) plasmids are costly and additionally plasmid donors lose all reproductive fitness after conjugation.

## Discussion

Here, I have demonstrated the importance incorporating the timing of plasmid costs into plasmid population models. When costs are only paid as a result of conjugation, they will likely not be sufficient to prevent a plasmid from spreading throughout a host population. Furthermore, even when costs are mixed (some owing to conjugation, some not) the conjugation related costs are unlikely to determine the fate of the plasmid population.

The actual source of costs of plasmids in nature is a question that requires investigation. Laboratory based experiments can miss key variables that are important to bacteria and plasmid ecology in natural communities. Indeed, the idea that plasmids are costly and may be lost from host populations due to these costs comes from the laboratory setting, i.e., we do not have good direct evidence of plasmid costs in natural communities. However, we can infer that such costs will exist (at least during and after conjugation) due to both the metabolic cost of conjugations and the large number and variety of phages that target the pilus (Jalasvuori et al. 2011). The former may or may not translate into a significant decrease in host fitness in nature, but it is difficult to conceive that the latter would not.

Given the results reported here it is unsurprising that plasmids are common in nature. Experiments have shown that the genetic incompatibilities experienced when unfamiliar plasmid/host combinations are first brought together are readily overcome by compensatory evolution within relatively few generations (Harrison et al. 2015; San Millan et al. 2015; Loftie-Eaton et al. 2017; Hall et al. 2020, 2021). This leaves the question of the more difficult to avoid costs, which will not be apparent in most experimental systems (nutrient rich media reduces the burden of metabolic costs and absence of phage negates target attacks on pili). Our demonstration that such conjugation related costs should not be expected stop a plasmid population from spread, combined with the afore mentioned experimental data on compensatory evolution suggests that the prevalence of plasmids in microbial communities is not paradoxical after all.

## Methods

### Analytical solutions

For the dynamical systems represented by Equations 1-2 and 3-6 I assessed the plasmids ability to invade the population by considering whether or not the fixed point where the density of plasmid bearing cells is 0 and the plasmid free cells = *k* was stable, using linear stability analysis (OTTO and DAY 2007). For Equations 1-2 the eigen values of the jacobian are

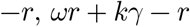

−*r* will always be negative, so this point will only be unstable (the plasmid will invade) if

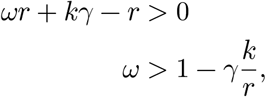

For Equations 3-6 the eigen values of the jacobian are

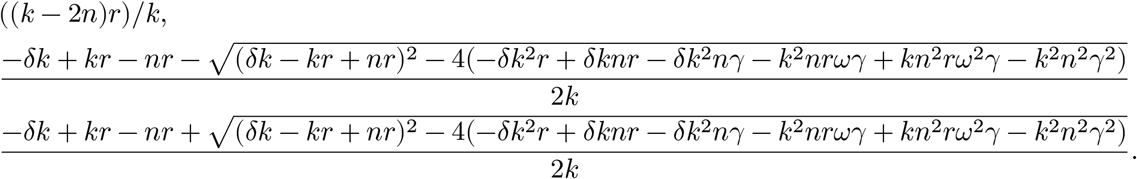

Given that *n* = *k* at this fixed point, the third eigenvalue will be positive so long as the term within second brackets the square root is negative. This will always be the case unless *γ* = 0.

### Numerical methods

Differential equations were solved using the Runge–Kutta order 5 (Tsitouras 2011) solver implemented in `DifferentialEquations.jl` (Rackauckas and Nie 2017) in Julia (Bezanson et al. 2017), with results plotted in (Danisch and Krumbiegel 2021).

## Supporting information

Supplementary figures showing results for a range of parameter values

## Code availability

All code to reproduce this work is available at https://github.com/EvoArt/plasmid-costs

