## Supplementary figures showing results for a range of parameter values for "Conjugation related costs have reduced impact on *in silico* plasmid persistence"

ω2 = 0.9, λ = 7.5e-6, d = 0.025

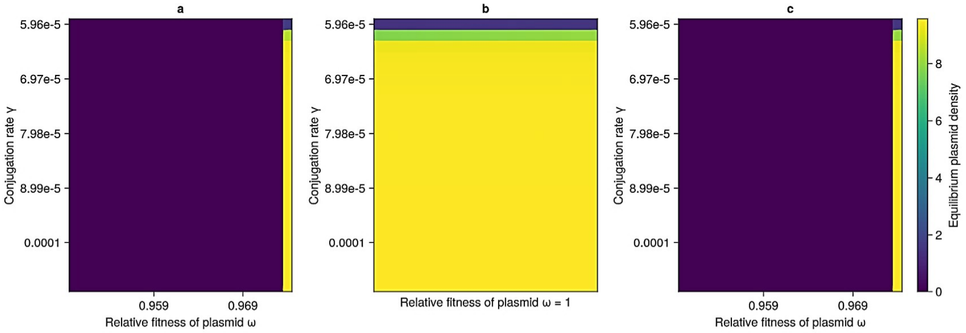


ω2 = 0.9, λ = 5.0e-6, d = 0.05

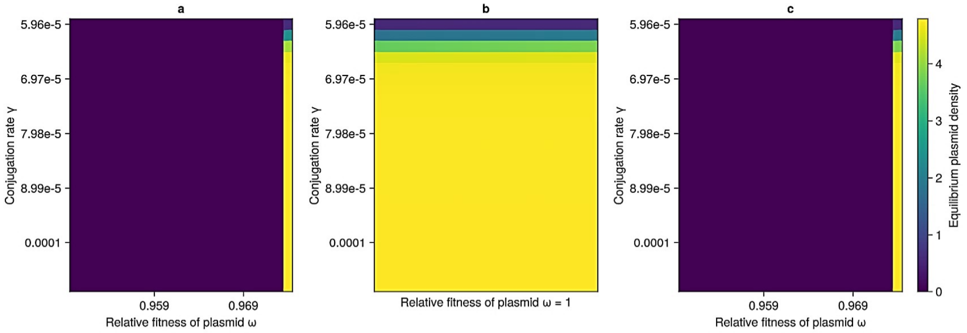


ω2 = 0.9, λ = 7.5e-6, d = 0.05

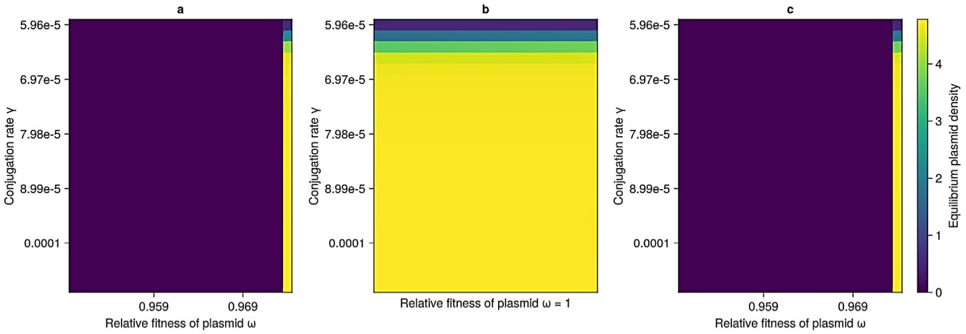


ω2 = 0.9, λ = 7.5e-6, d = 5.0e-5

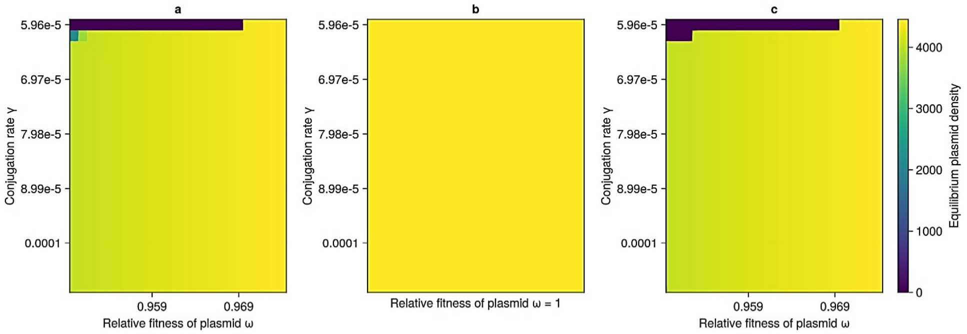

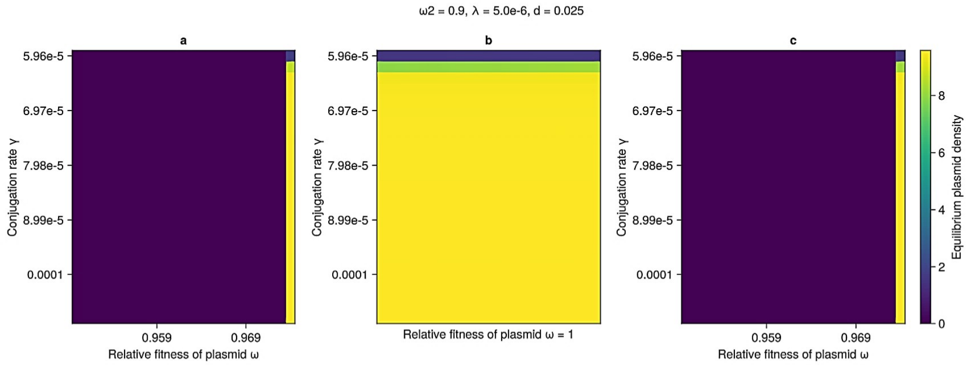
ω2 = 0.9, λ = 5.0e-6, d = 0.025
a
b
c
Conjugation rate γ
5.96e-5
6.97e-5
7.98e-5
8.99e-5
0.0001
0.959
0.969
Relative fitness of plasmid ω
Relative fitness of plasmid ω = 1
Equilibrium plasmid density
0
2
4
6
8

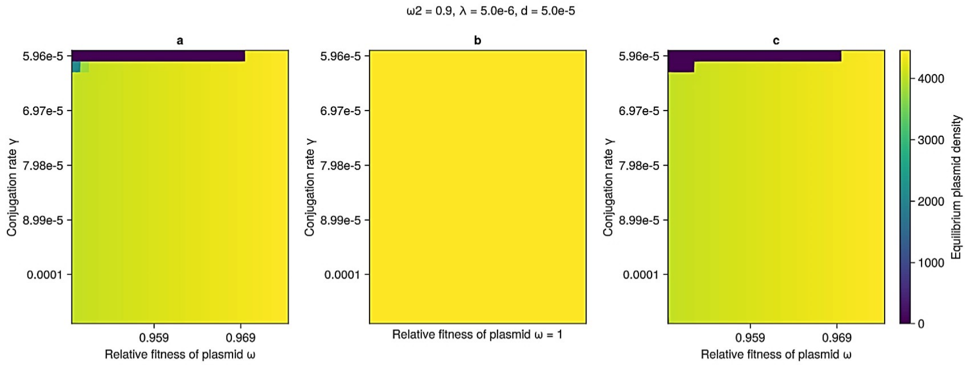
ω2 = 0.9, λ = 5.0e-6, d = 5.0e-5
a
b
c
Conjugation rate γ
5.96e-5
6.97e-5
7.98e-5
8.99e-5
0.0001
0.959
0.969
Relative fitness of plasmid ω
Relative fitness of plasmid ω = 1
Equilibrium plasmid density
0
1000
2000
3000
4000

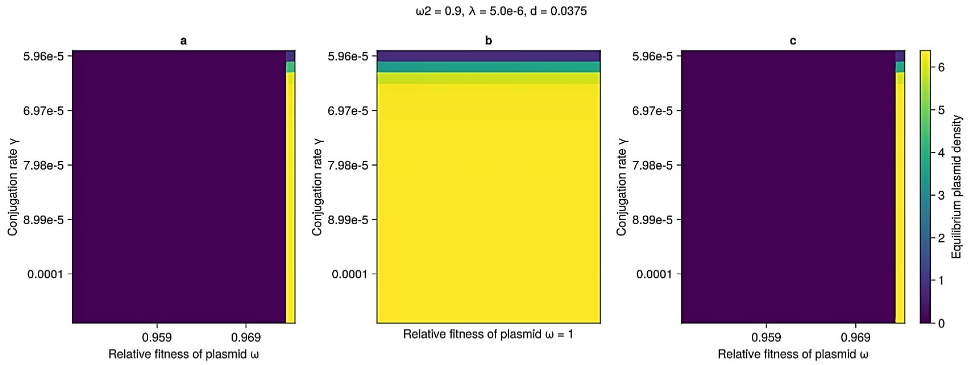
ω2 = 0.9, λ = 5.0e-6, d = 0.0375
a
b
c
Conjugation rate γ
5.96e-5
6.97e-5
7.98e-5
8.99e-5
0.0001
0.959
0.969
Relative fitness of plasmid ω
Relative fitness of plasmid ω = 1
Equilibrium plasmid density
0
1
2
3
4
5
6

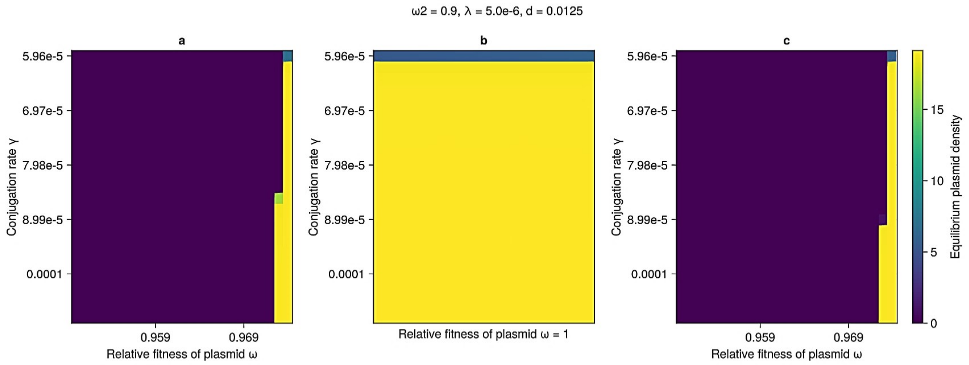
ω2 = 0.9, λ = 5.0e-6, d = 0.0125
a
b
c
Conjugation rate γ
5.96e-5
6.97e-5
7.98e-5
8.99e-5
0.0001
0.959
0.969
Relative fitness of plasmid ω
Relative fitness of plasmid ω = 1
Equilibrium plasmid density
0
5
10
15

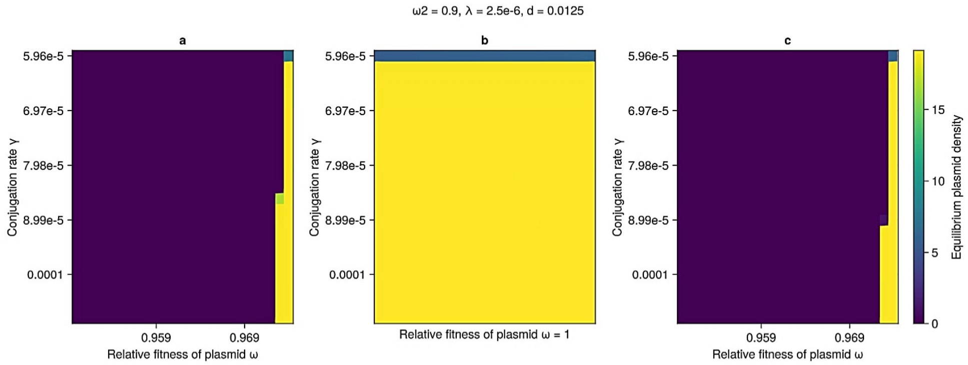
ω2 = 0.9, λ = 2.5e-6, d = 0.0125
a
b
c
Conjugation rate γ
5.96e-5
6.97e-5
7.98e-5
8.99e-5
0.0001
0.959
0.969
Relative fitness of plasmid ω
Relative fitness of plasmid ω = 1
Equilibrium plasmid density
0
5
10
15

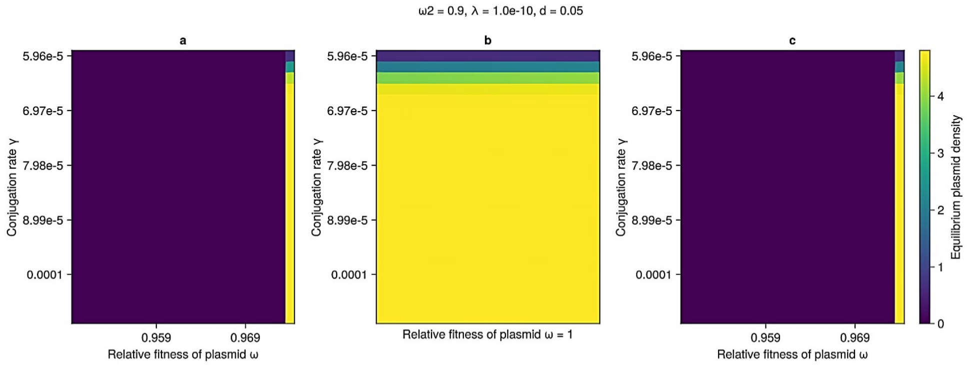
ω2 = 0.9, λ = 1.0e-10, d = 0.05
a
b
c
Conjugation rate γ
5.96e-5
6.97e-5
7.98e-5
8.99e-5
0.0001
0.959
0.969
Relative fitness of plasmid ω
Relative fitness of plasmid ω = 1
Equilibrium plasmid density
0
1
2
3
4

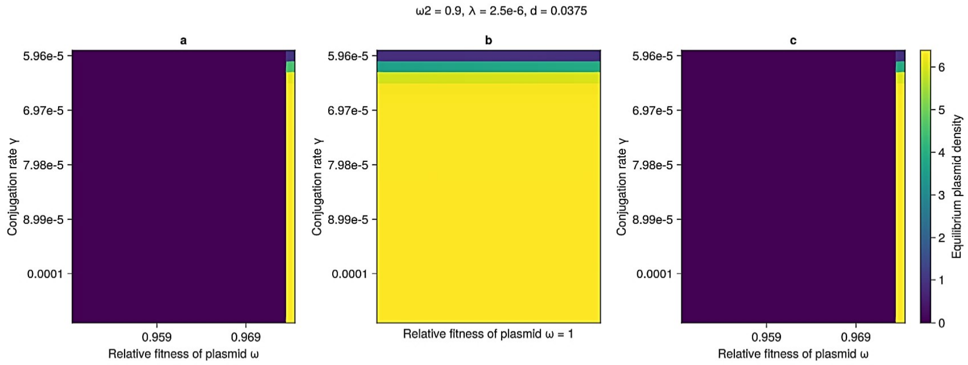
ω2 = 0.9, λ = 2.5e-6, d = 0.0375
a
b
c
Conjugation rate γ
5.96e-5
6.97e-5
7.98e-5
8.99e-5
0.0001
0.959
0.969
Relative fitness of plasmid ω
Relative fitness of plasmid ω = 1
Equilibrium plasmid density
0
1
2
3
4
5
6

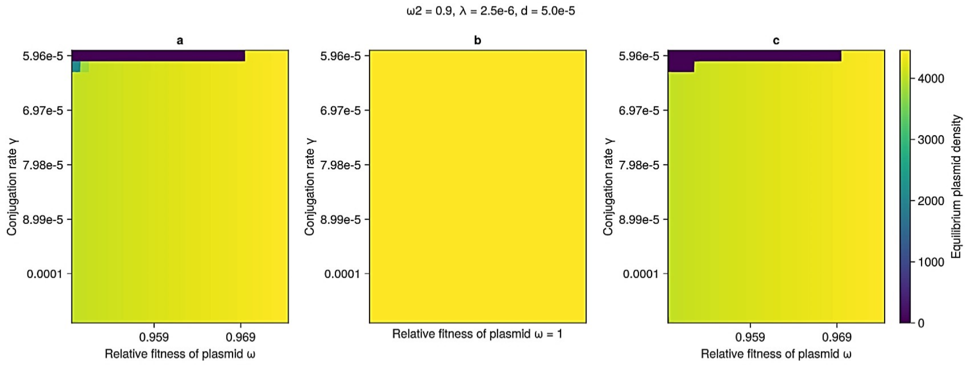
ω2 = 0.9, λ = 2.5e-6, d = 5.0e-5
a
b
c
Conjugation rate γ
5.96e-5
6.97e-5
7.98e-5
8.99e-5
0.0001
0.959
0.969
Relative fitness of plasmid ω
Relative fitness of plasmid ω = 1
Equilibrium plasmid density
0
1000
2000
3000
4000

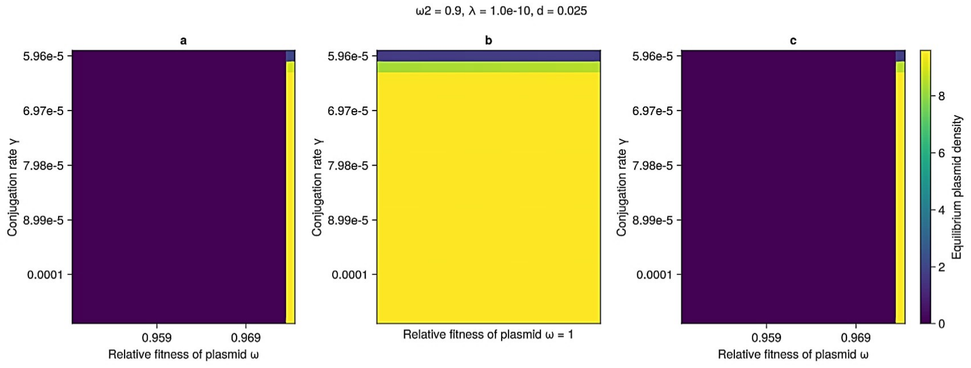
ω2 = 0.9, λ = 1.0e-10, d = 0.025
a
b
c
Conjugation rate γ
5.96e-5
6.97e-5
7.98e-5
8.99e-5
0.0001
0.959
0.969
Relative fitness of plasmid ω
Relative fitness of plasmid ω = 1
Equilibrium plasmid density
0
2
4
6
8

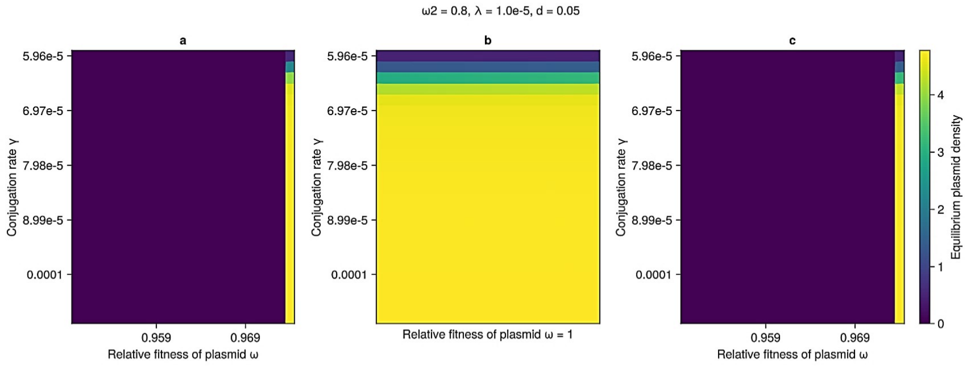
ω2 = 0.8, λ = 1.0e-5, d = 0.05
a
b
c
Conjugation rate γ
5.96e-5
6.97e-5
7.98e-5
8.99e-5
0.0001
0.959
0.969
Relative fitness of plasmid ω
Relative fitness of plasmid ω = 1
Equilibrium plasmid density
0
1
2
3
4

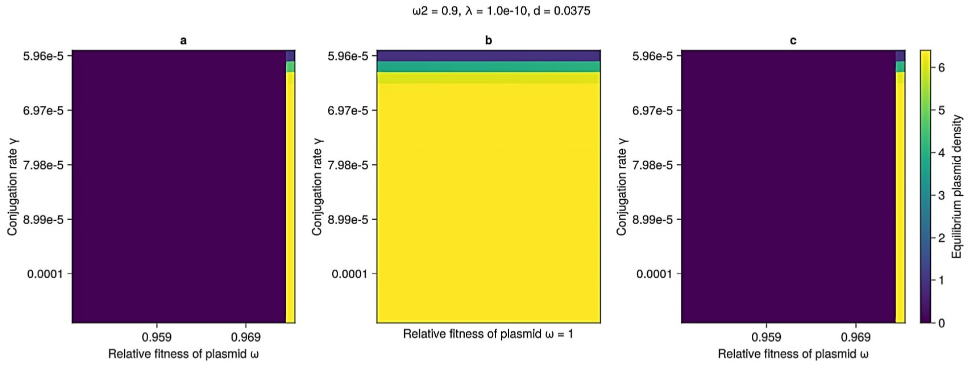
ω2 = 0.9, λ = 1.0e-10, d = 0.0375
a
b
c
Conjugation rate γ
5.96e-5
6.97e-5
7.98e-5
8.99e-5
0.0001
0.959
0.969
Relative fitness of plasmid ω
Relative fitness of plasmid ω = 1
Equilibrium plasmid density
0
1
2
3
4
5
6

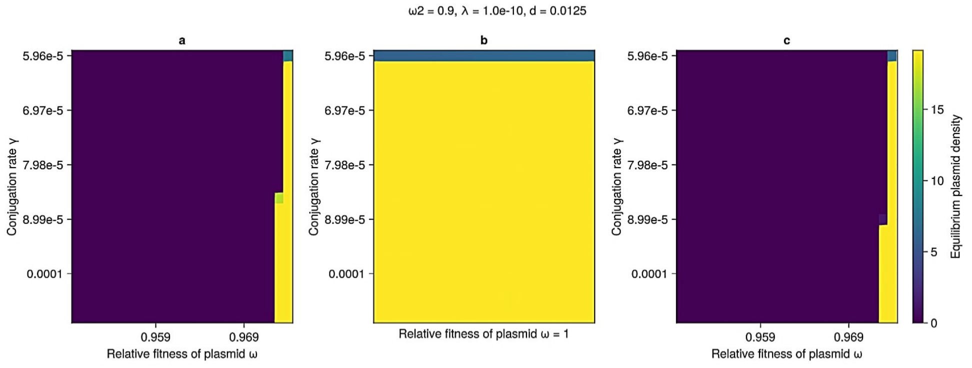
ω2 = 0.9, λ = 1.0e-10, d = 0.0125
a
b
c
Conjugation rate γ
5.96e-5
6.97e-5
7.98e-5
8.99e-5
0.0001
0.959
0.969
Relative fitness of plasmid ω
Relative fitness of plasmid ω = 1
Equilibrium plasmid density
0
5
10
15

ω2 = 0.8, λ = 1.0e-5, d = 0.0125

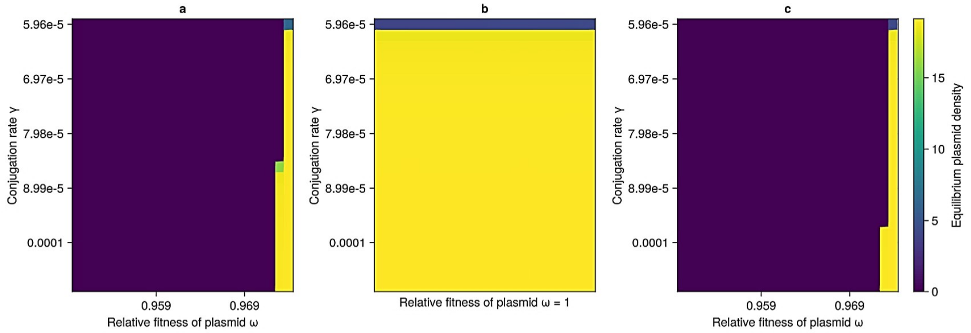


ω2 = 0.8, λ = 7.5e-6, d = 0.05

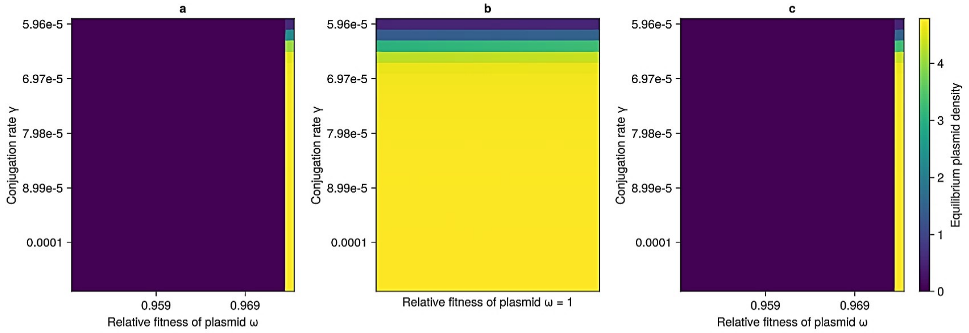


ω2 = 0.8, λ = 1.0e-5, d = 0.025

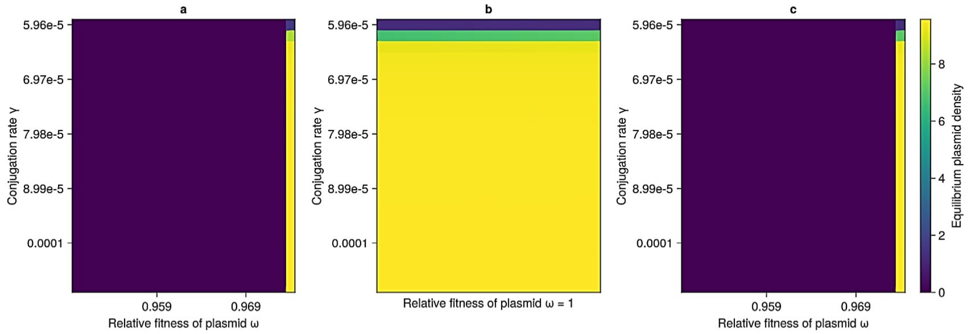


ω2 = 0.8, λ = 1.0e-5, d = 5.0e-5

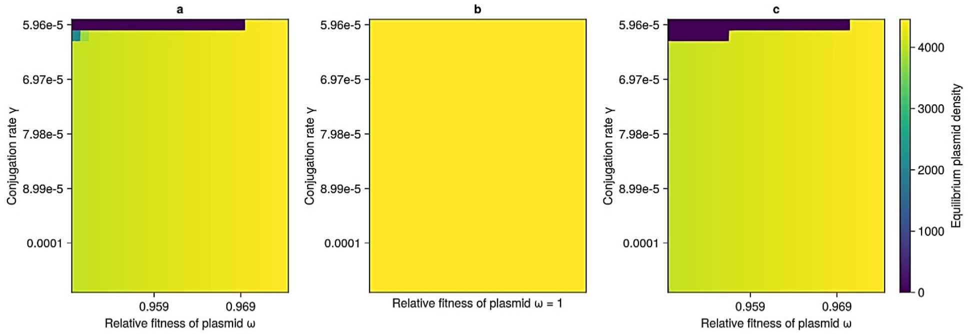

ω2 = 0.8, λ = 7.5e-6, d = 0.025

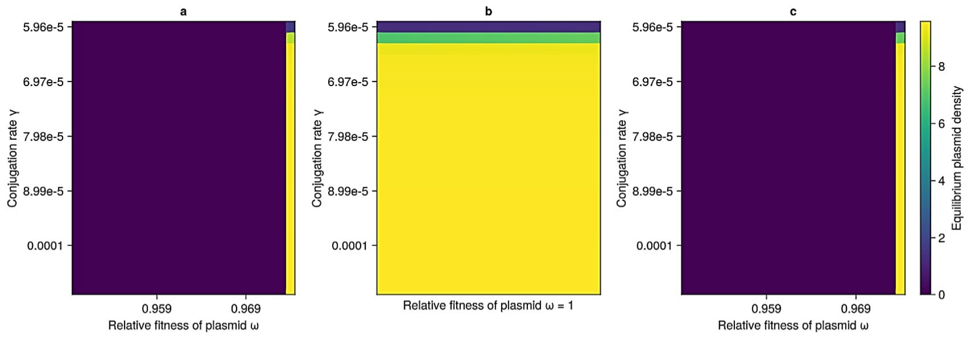


ω2 = 0.8, λ = 5.0e-6, d = 0.0375

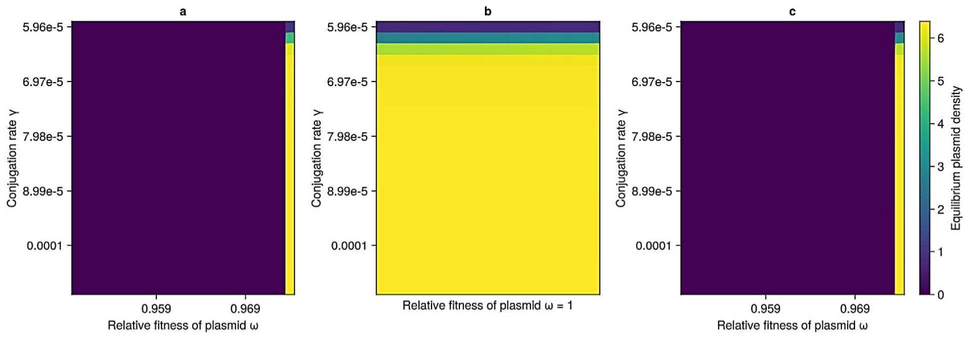


ω2 = 0.8, λ = 7.5e-6, d = 0.0375

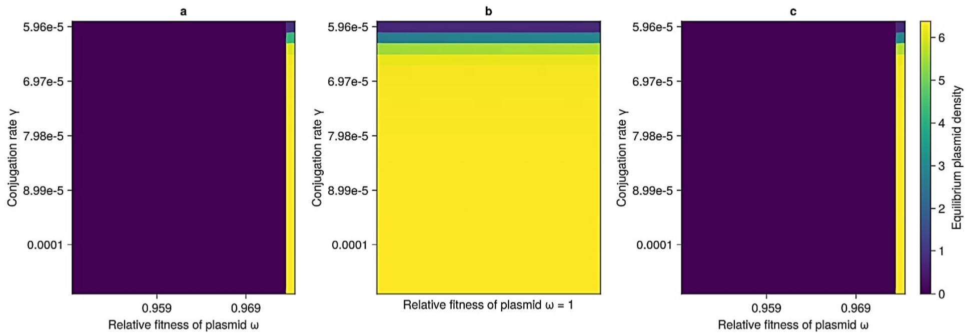


ω2 = 0.8, λ = 7.5e-6, d = 5.0e-5

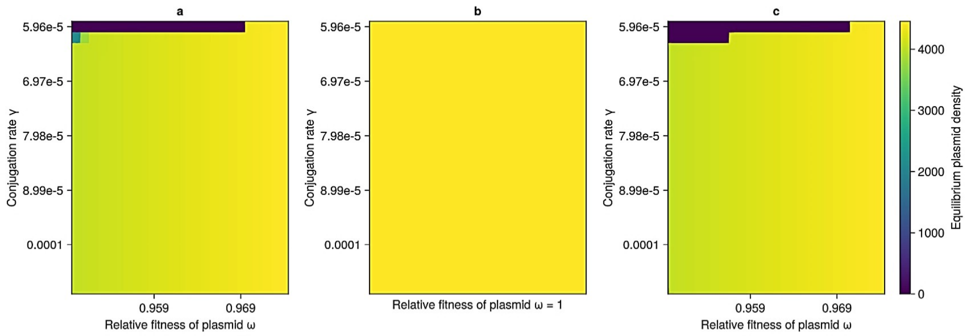

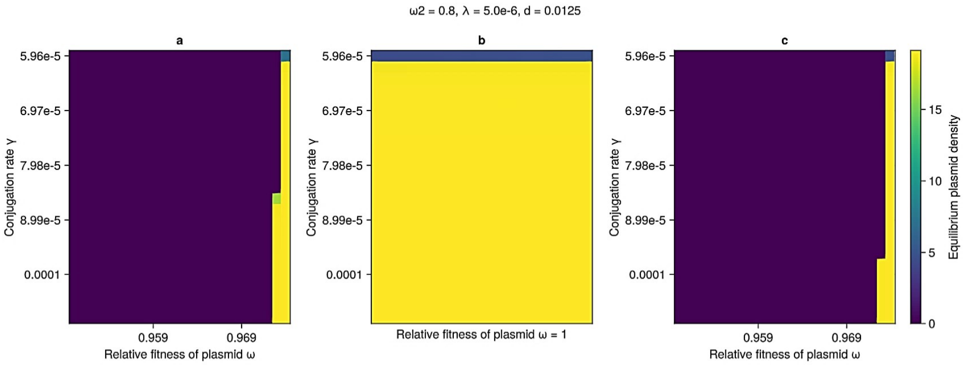
ω2 = 0.8, λ = 5.0e-6, d = 0.0125
a
b
c
5.96e-5
6.97e-5
7.98e-5
8.99e-5
0.0001
Conjugation rate γ
0.959
0.969
Relative fitness of plasmid ω
Relative fitness of plasmid ω = 1
0
5
10
15
Equilibrium plasmid density

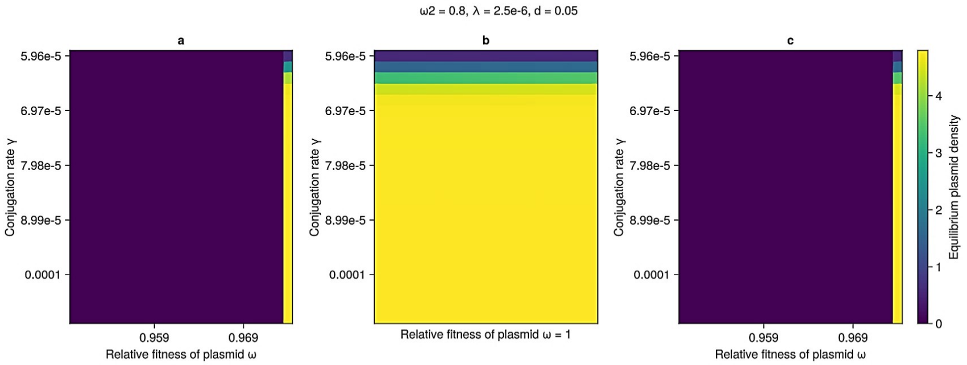
ω2 = 0.8, λ = 2.5e-6, d = 0.05
a
b
c
5.96e-5
6.97e-5
7.98e-5
8.99e-5
0.0001
Conjugation rate γ
0.959
0.969
Relative fitness of plasmid ω
Relative fitness of plasmid ω = 1
0
1
2
3
4
Equilibrium plasmid density

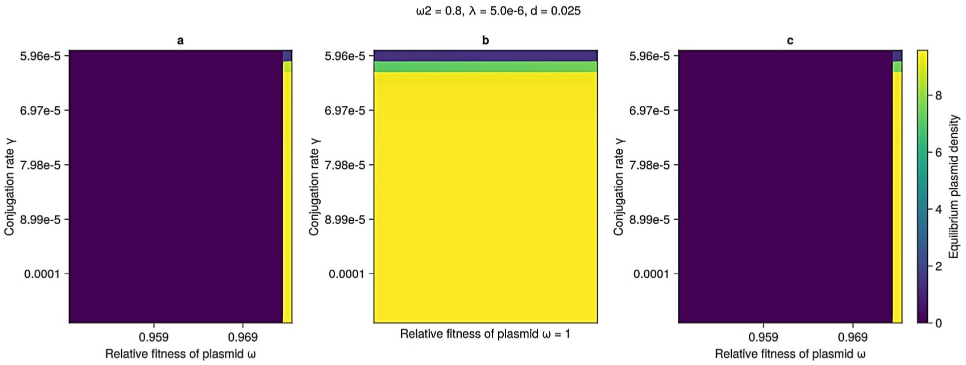
ω2 = 0.8, λ = 5.0e-6, d = 0.025
a
b
c
5.96e-5
6.97e-5
7.98e-5
8.99e-5
0.0001
Conjugation rate γ
0.959
0.969
Relative fitness of plasmid ω
Relative fitness of plasmid ω = 1
0
2
4
6
8
Equilibrium plasmid density

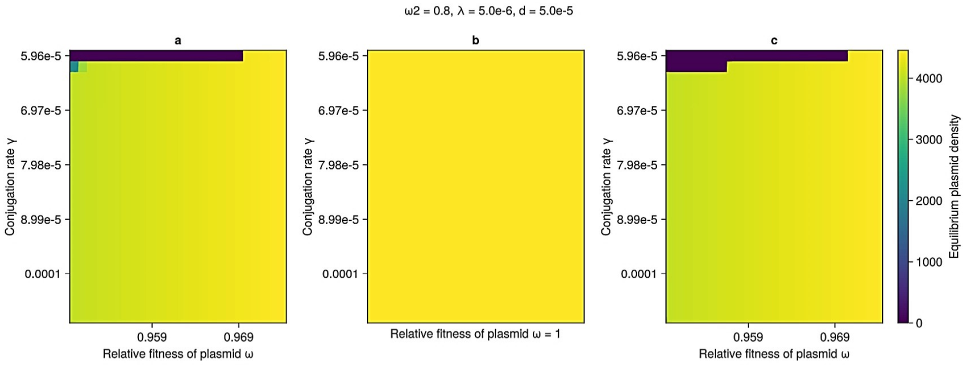
ω2 = 0.8, λ = 5.0e-6, d = 5.0e-5
a
b
c
5.96e-5
6.97e-5
7.98e-5
8.99e-5
0.0001
Conjugation rate γ
0.959
0.969
Relative fitness of plasmid ω
Relative fitness of plasmid ω = 1
0
1000
2000
3000
4000
Equilibrium plasmid density

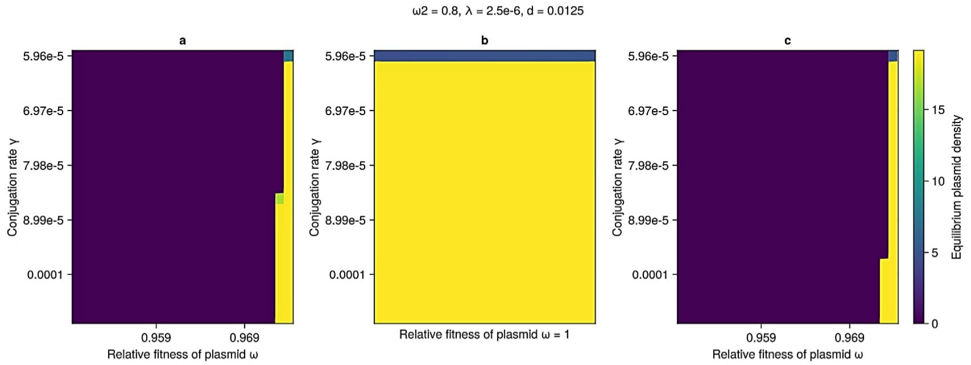

ω2 = 0.8, λ = 2.5e-6, d = 0.0125
a
b
c
Conjugation rate γ
5.96e-5
6.97e-5
7.98e-5
8.99e-5
0.0001
0.959
0.969
Relative fitness of plasmid ω
Relative fitness of plasmid ω = 1
Equilibrium plasmid density
0
5
10
15


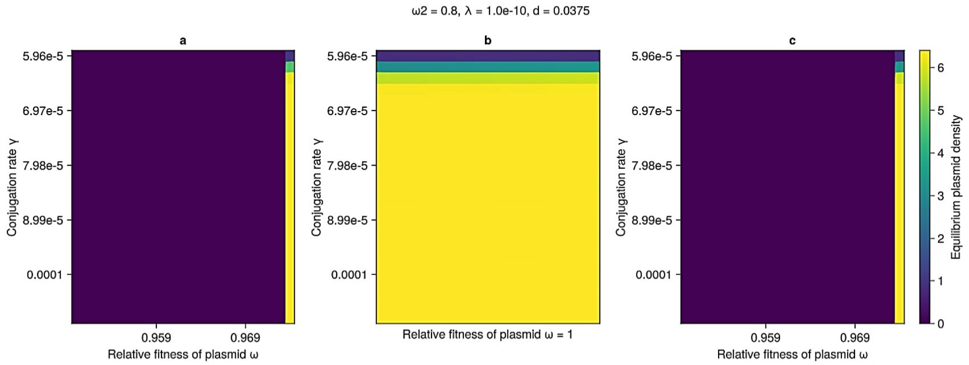

ω2 = 0.8, λ = 1.0e-10, d = 0.0375
a
b
c
Conjugation rate γ
5.96e-5
6.97e-5
7.98e-5
8.99e-5
0.0001
0.959
0.969
Relative fitness of plasmid ω
Relative fitness of plasmid ω = 1
Equilibrium plasmid density
0
1
2
3
4
5
6


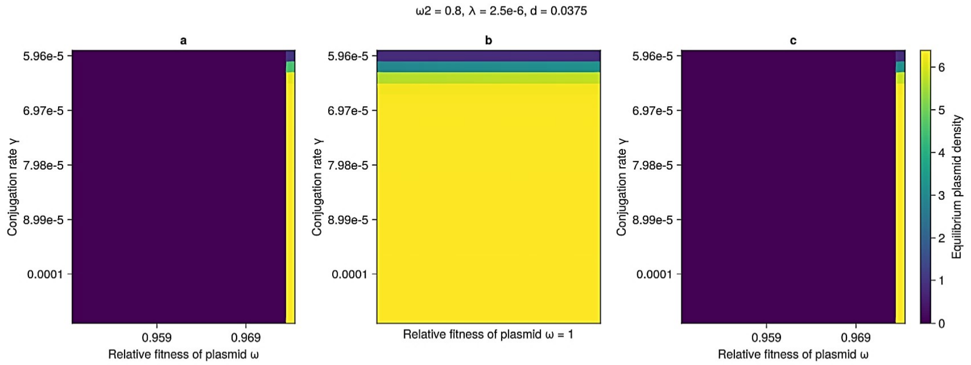

ω2 = 0.8, λ = 2.5e-6, d = 0.0375
a
b
c
Conjugation rate γ
5.96e-5
6.97e-5
7.98e-5
8.99e-5
0.0001
0.959
0.969
Relative fitness of plasmid ω
Relative fitness of plasmid ω = 1
Equilibrium plasmid density
0
1
2
3
4
5
6


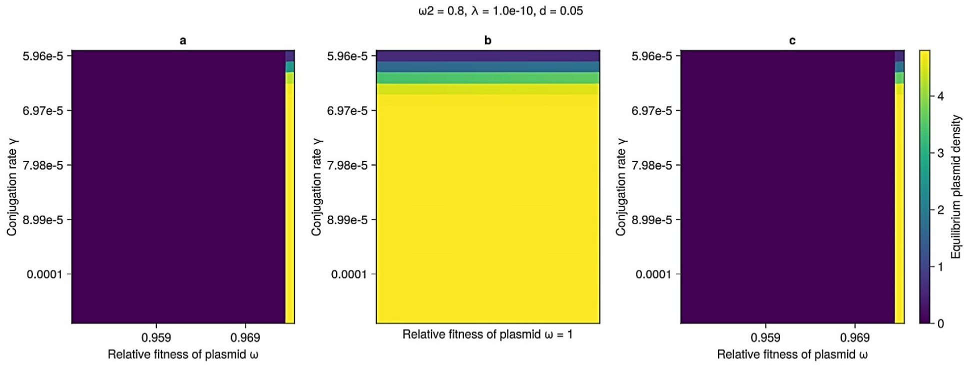

ω2 = 0.8, λ = 1.0e-10, d = 0.05
a
b
c
Conjugation rate γ
5.96e-5
6.97e-5
7.98e-5
8.99e-5
0.0001
0.959
0.969
Relative fitness of plasmid ω
Relative fitness of plasmid ω = 1
Equilibrium plasmid density
0
1
2
3
4

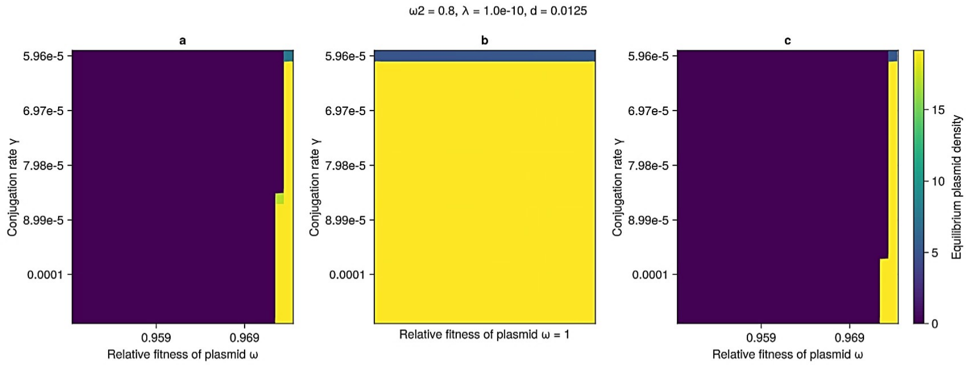
ω2 = 0.8, λ = 1.0e-10, d = 0.0125
a
b
c
Conjugation rate γ
5.96e-5
6.97e-5
7.98e-5
8.99e-5
0.0001
0.959
0.969
Relative fitness of plasmid ω
Relative fitness of plasmid ω = 1
Equilibrium plasmid density
0
5
10
15

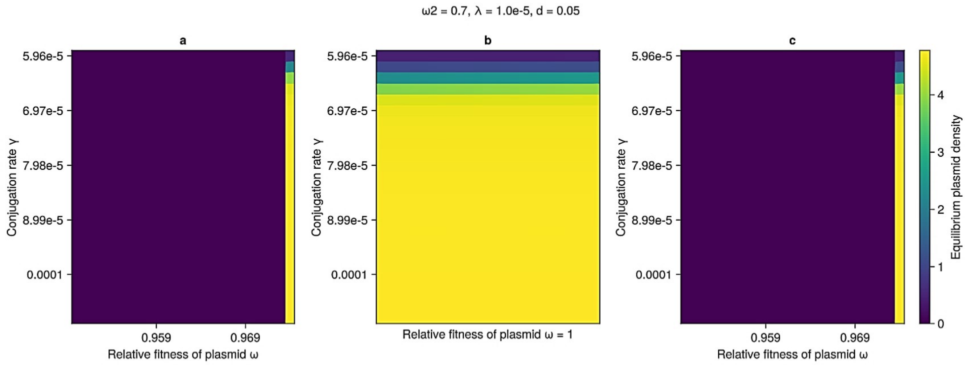
ω2 = 0.7, λ = 1.0e-5, d = 0.05
a
b
c
Conjugation rate γ
5.96e-5
6.97e-5
7.98e-5
8.99e-5
0.0001
0.959
0.969
Relative fitness of plasmid ω
Relative fitness of plasmid ω = 1
Equilibrium plasmid density
0
1
2
3
4

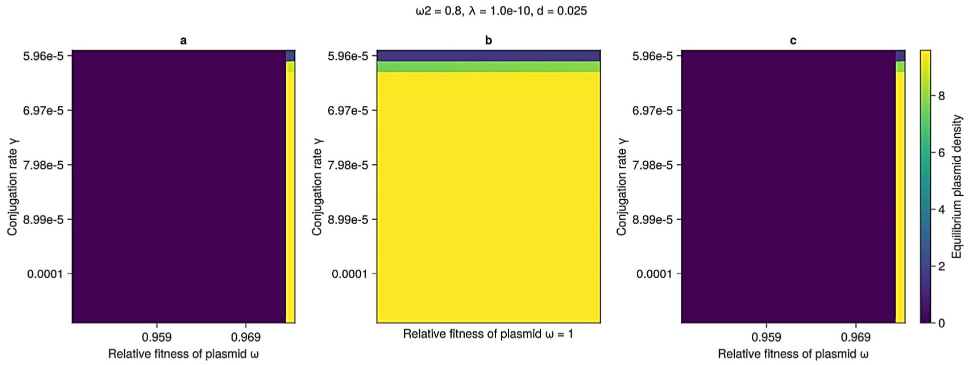
ω2 = 0.8, λ = 1.0e-10, d = 0.025
a
b
c
Conjugation rate γ
5.96e-5
6.97e-5
7.98e-5
8.99e-5
0.0001
0.959
0.969
Relative fitness of plasmid ω
Relative fitness of plasmid ω = 1
Equilibrium plasmid density
0
2
4
6
8

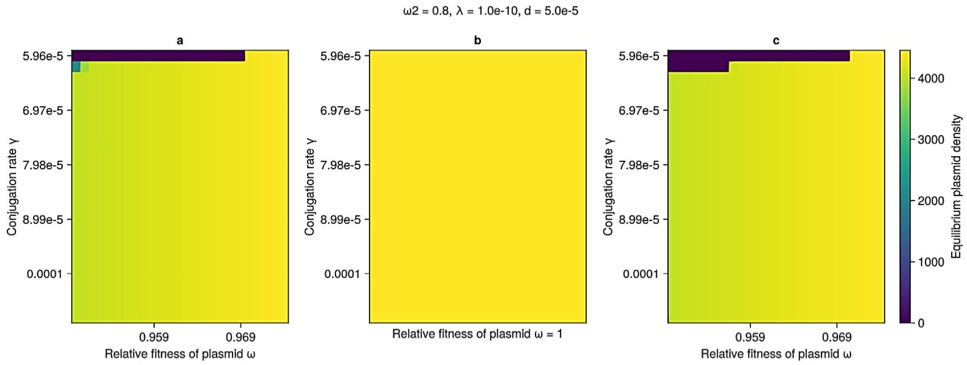
ω2 = 0.8, λ = 1.0e-10, d = 5.0e-5
a
b
c
Conjugation rate γ
5.96e-5
6.97e-5
7.98e-5
8.99e-5
0.0001
0.959
0.969
Relative fitness of plasmid ω
Relative fitness of plasmid ω = 1
Equilibrium plasmid density
0
1000
2000
3000
4000

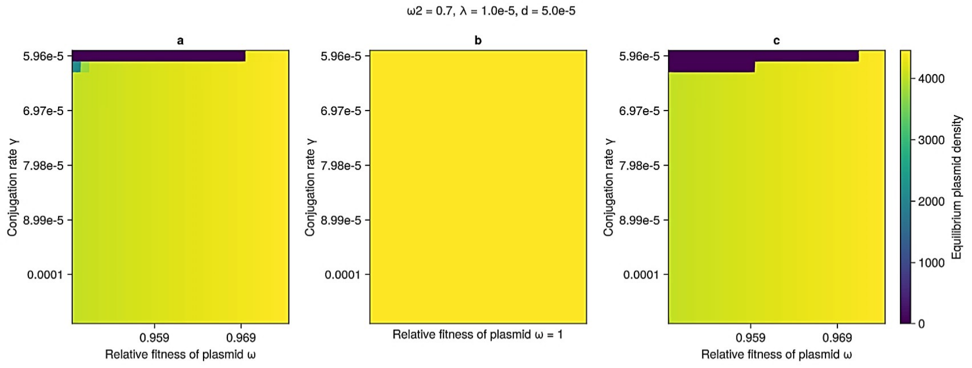
ω2 = 0.7, λ = 1.0e-5, d = 5.0e-5
a
b
c
Conjugation rate γ
5.96e-5
6.97e-5
7.98e-5
8.99e-5
0.0001
0.959
0.969
Relative fitness of plasmid ω
Relative fitness of plasmid ω = 1
Equilibrium plasmid density
0
1000
2000
3000
4000

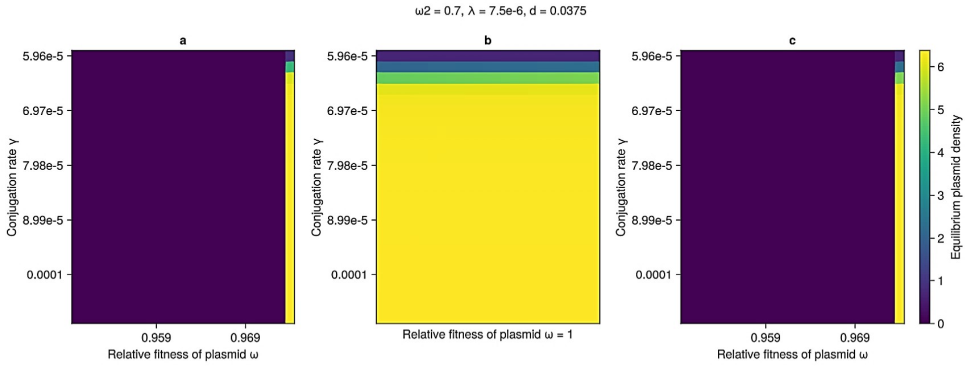
ω2 = 0.7, λ = 7.5e-6, d = 0.0375
a
b
c
Conjugation rate γ
5.96e-5
6.97e-5
7.98e-5
8.99e-5
0.0001
0.959
0.969
Relative fitness of plasmid ω
Relative fitness of plasmid ω = 1
Equilibrium plasmid density
0
1
2
3
4
5
6

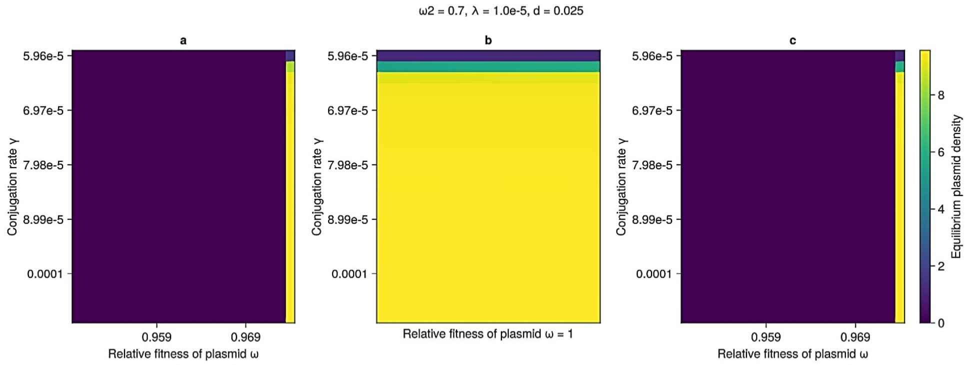
ω2 = 0.7, λ = 1.0e-5, d = 0.025
a
b
c
Conjugation rate γ
5.96e-5
6.97e-5
7.98e-5
8.99e-5
0.0001
0.959
0.969
Relative fitness of plasmid ω
Relative fitness of plasmid ω = 1
Equilibrium plasmid density
0
2
4
6
8

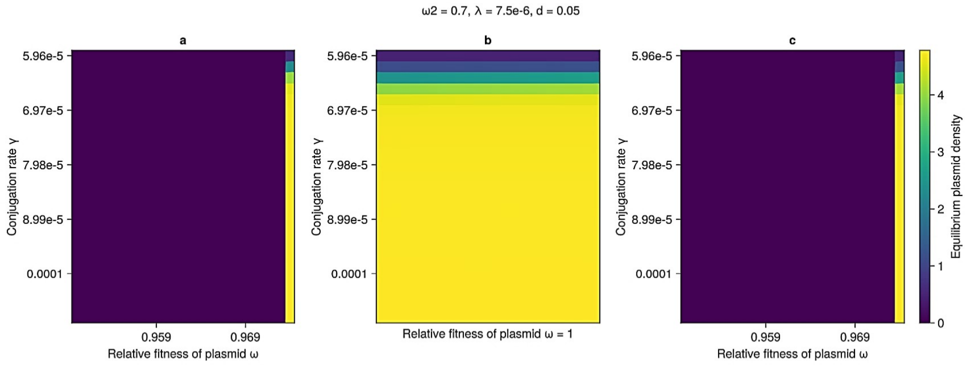
ω2 = 0.7, λ = 7.5e-6, d = 0.05
a
b
c
Conjugation rate γ
5.96e-5
6.97e-5
7.98e-5
8.99e-5
0.0001
0.959
0.969
Relative fitness of plasmid ω
Relative fitness of plasmid ω = 1
Equilibrium plasmid density
0
1
2
3
4

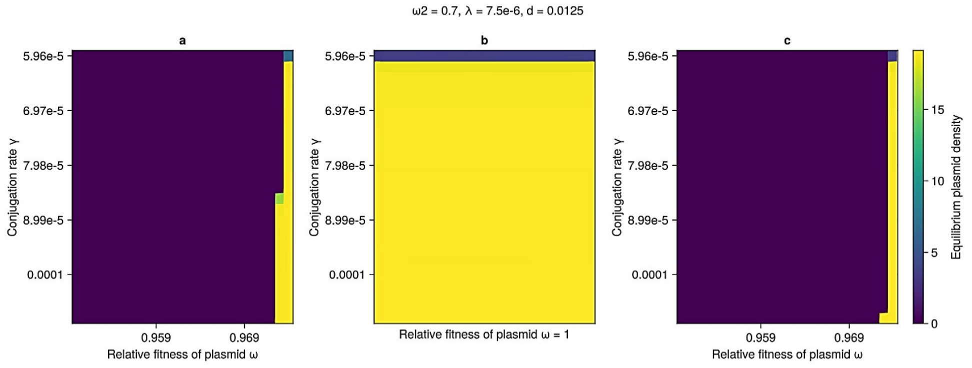
ω2 = 0.7, λ = 7.5e-6, d = 0.0125
a
b
c
Conjugation rate γ
5.96e-5
6.97e-5
7.98e-5
8.99e-5
0.0001
0.959
0.969
Relative fitness of plasmid ω
Relative fitness of plasmid ω = 1
Equilibrium plasmid density
0
5
10
15

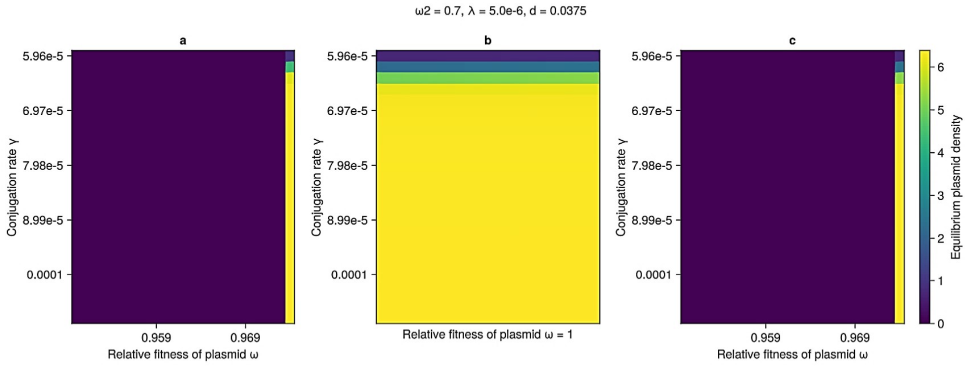
ω2 = 0.7, λ = 5.0e-6, d = 0.0375
a
b
c
Conjugation rate γ
5.96e-5
6.97e-5
7.98e-5
8.99e-5
0.0001
0.959
0.969
Relative fitness of plasmid ω
Relative fitness of plasmid ω = 1
Equilibrium plasmid density
0
1
2
3
4
5
6

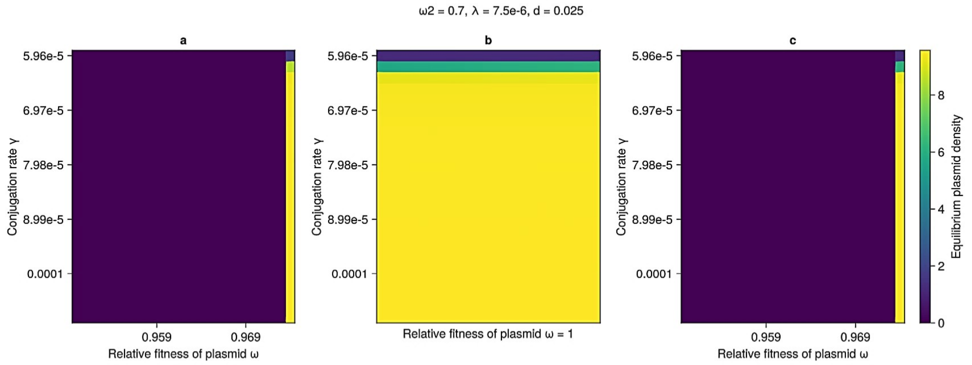
ω2 = 0.7, λ = 7.5e-6, d = 0.025
a
b
c
Conjugation rate γ
5.96e-5
6.97e-5
7.98e-5
8.99e-5
0.0001
0.959
0.969
Relative fitness of plasmid ω
Relative fitness of plasmid ω = 1
Equilibrium plasmid density
0
2
4
6
8

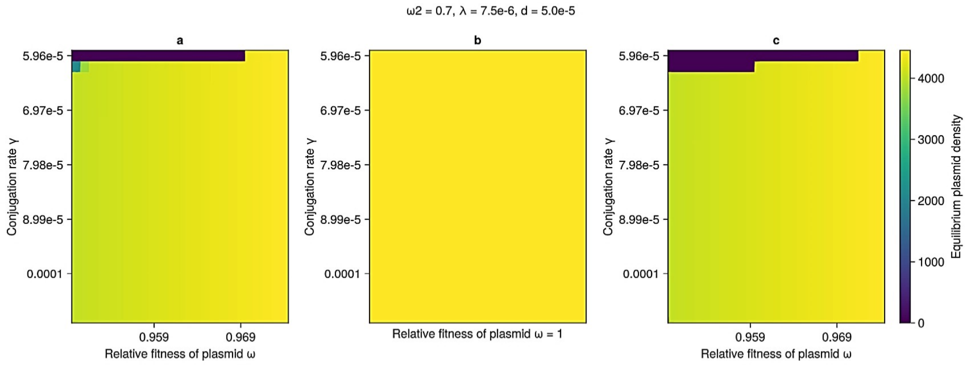
ω2 = 0.7, λ = 7.5e-6, d = 5.0e-5
a
b
c
Conjugation rate γ
5.96e-5
6.97e-5
7.98e-5
8.99e-5
0.0001
0.959
0.969
Relative fitness of plasmid ω
Relative fitness of plasmid ω = 1
Equilibrium plasmid density
0
1000
2000
3000
4000

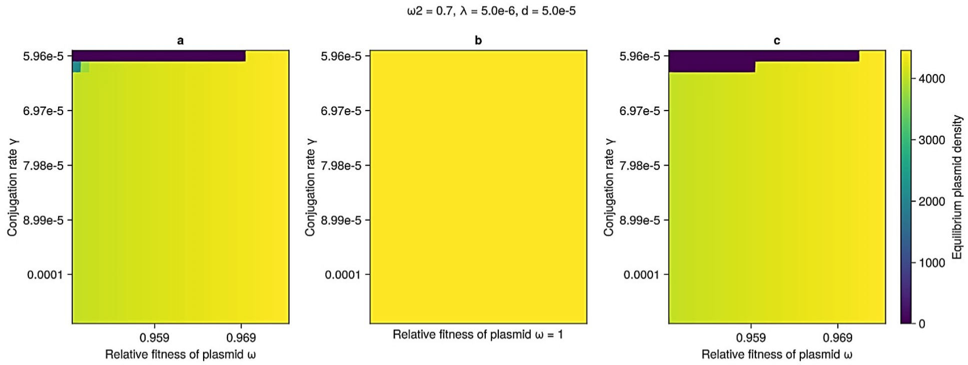
ω2 = 0.7, λ = 5.0e-6, d = 5.0e-5
a
b
c
Conjugation rate γ
5.96e-5
6.97e-5
7.98e-5
8.99e-5
0.0001
0.959
0.969
Relative fitness of plasmid ω
Relative fitness of plasmid ω = 1
Equilibrium plasmid density
0
1000
2000
3000
4000

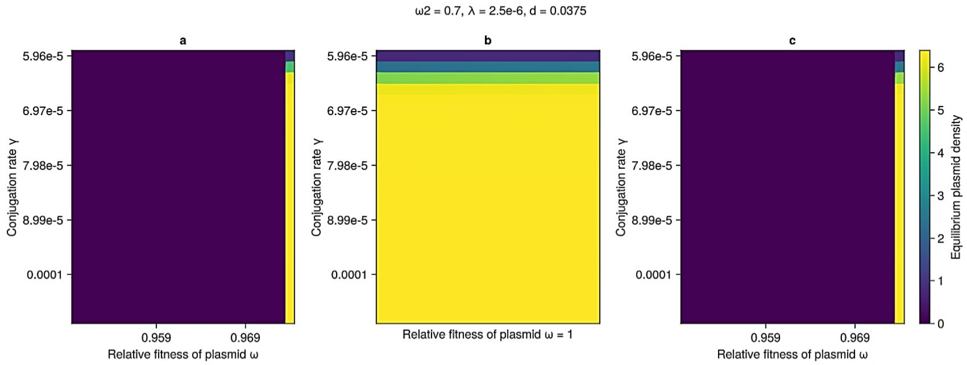
ω2 = 0.7, λ = 2.5e-6, d = 0.0375
a
b
c
Conjugation rate γ
5.96e-5
6.97e-5
7.98e-5
8.99e-5
0.0001
0.959
0.969
Relative fitness of plasmid ω
Relative fitness of plasmid ω = 1
Equilibrium plasmid density
0
1
2
3
4
5
6

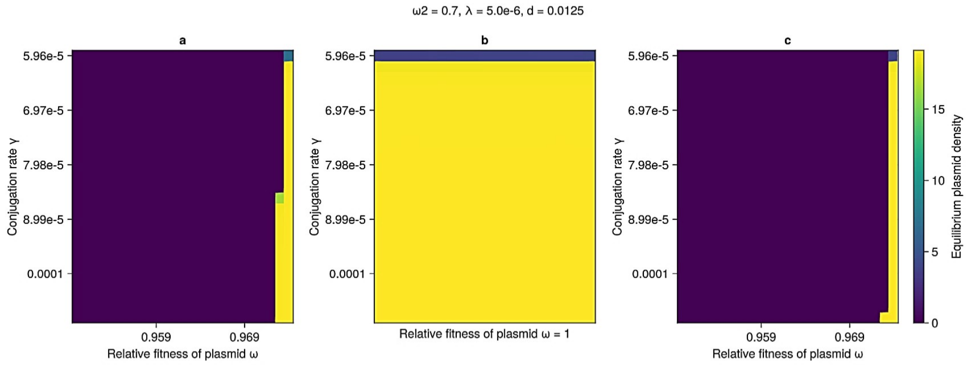
ω2 = 0.7, λ = 5.0e-6, d = 0.0125
a
b
c
Conjugation rate γ
5.96e-5
6.97e-5
7.98e-5
8.99e-5
0.0001
0.959
0.969
Relative fitness of plasmid ω
Relative fitness of plasmid ω = 1
Equilibrium plasmid density
0
5
10
15

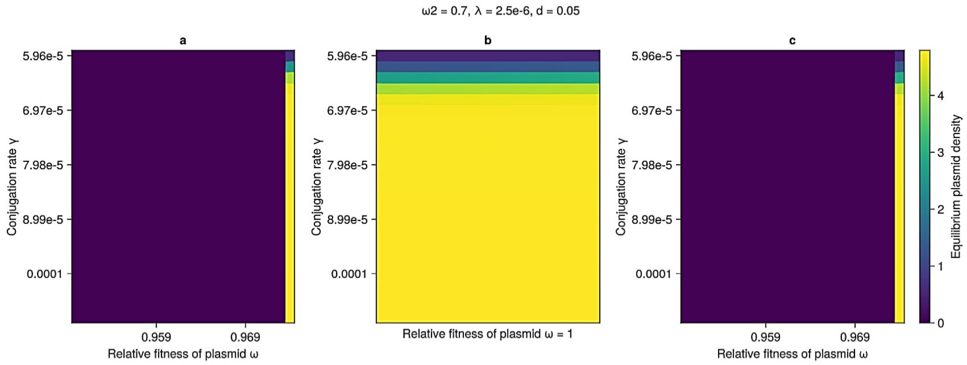
ω2 = 0.7, λ = 2.5e-6, d = 0.05
a
b
c
Conjugation rate γ
5.96e-5
6.97e-5
7.98e-5
8.99e-5
0.0001
0.959
0.969
Relative fitness of plasmid ω
Relative fitness of plasmid ω = 1
Equilibrium plasmid density
0
1
2
3
4

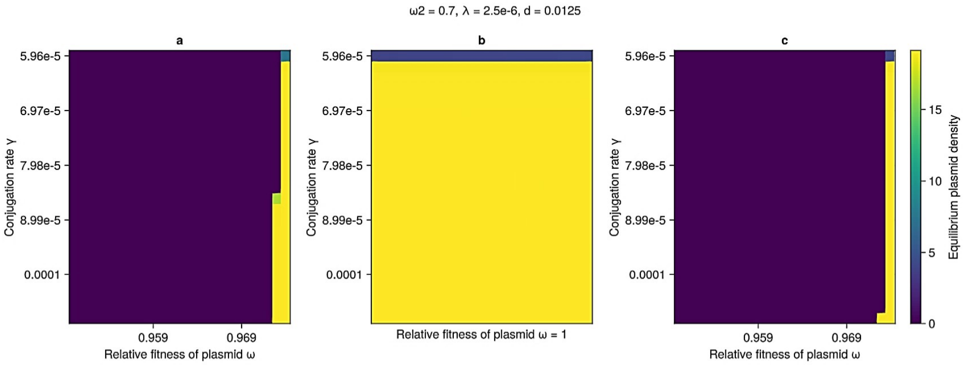

ω2 = 0.7, λ = 2.5e-6, d = 0.0125
a
b
c
Conjugation rate γ
5.96e-5
6.97e-5
7.98e-5
8.99e-5
0.0001
0.959
0.969
Relative fitness of plasmid ω
Relative fitness of plasmid ω = 1
Equilibrium plasmid density
0
5
10
15


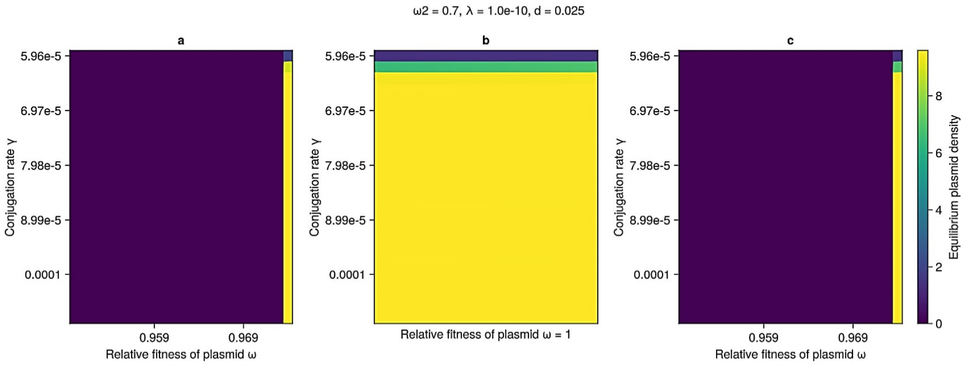

ω2 = 0.7, λ = 1.0e-10, d = 0.025
a
b
c
Conjugation rate γ
5.96e-5
6.97e-5
7.98e-5
8.99e-5
0.0001
0.959
0.969
Relative fitness of plasmid ω
Relative fitness of plasmid ω = 1
Equilibrium plasmid density
0
2
4
6
8


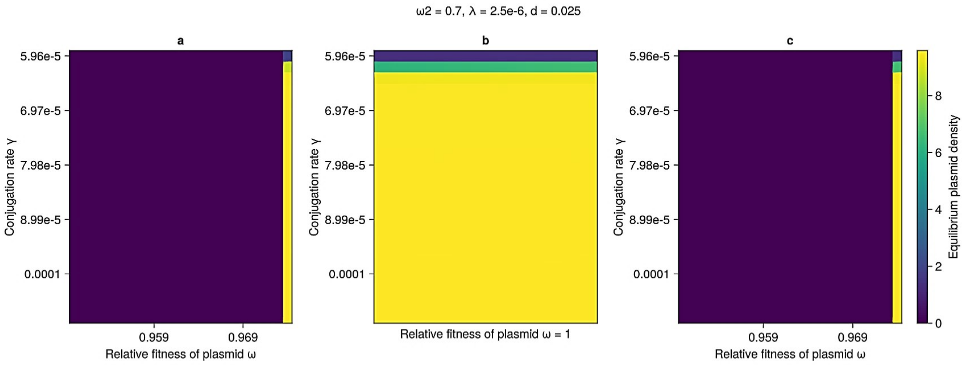

ω2 = 0.7, λ = 2.5e-6, d = 0.025
a
b
c
Conjugation rate γ
5.96e-5
6.97e-5
7.98e-5
8.99e-5
0.0001
0.959
0.969
Relative fitness of plasmid ω
Relative fitness of plasmid ω = 1
Equilibrium plasmid density
0
2
4
6
8


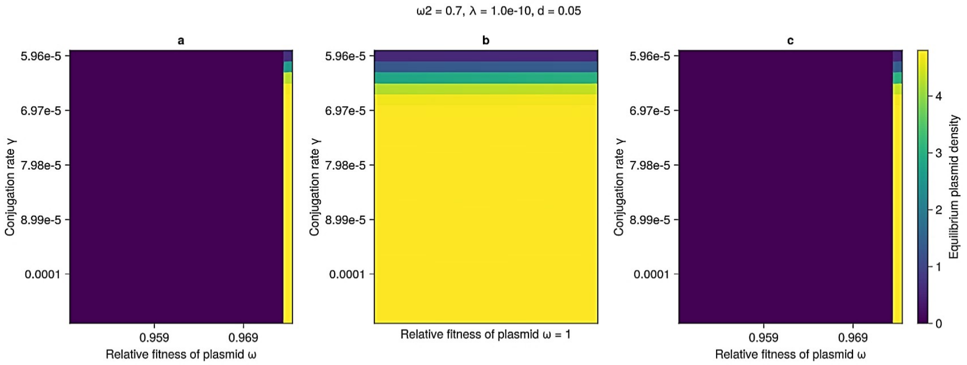

ω2 = 0.7, λ = 1.0e-10, d = 0.05
a
b
c
Conjugation rate γ
5.96e-5
6.97e-5
7.98e-5
8.99e-5
0.0001
0.959
0.969
Relative fitness of plasmid ω
Relative fitness of plasmid ω = 1
Equilibrium plasmid density
0
1
2
3
4

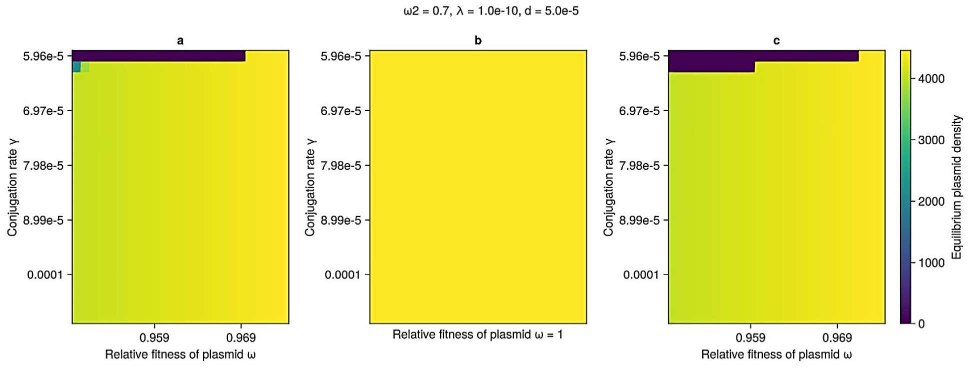
ω2 = 0.7, λ = 1.0e-10, d = 5.0e-5
a
b
c
Conjugation rate γ
5.96e-5
6.97e-5
7.98e-5
8.99e-5
0.0001
0.959
0.969
Relative fitness of plasmid ω
Relative fitness of plasmid ω = 1
Equilibrium plasmid density
0
1000
2000
3000
4000

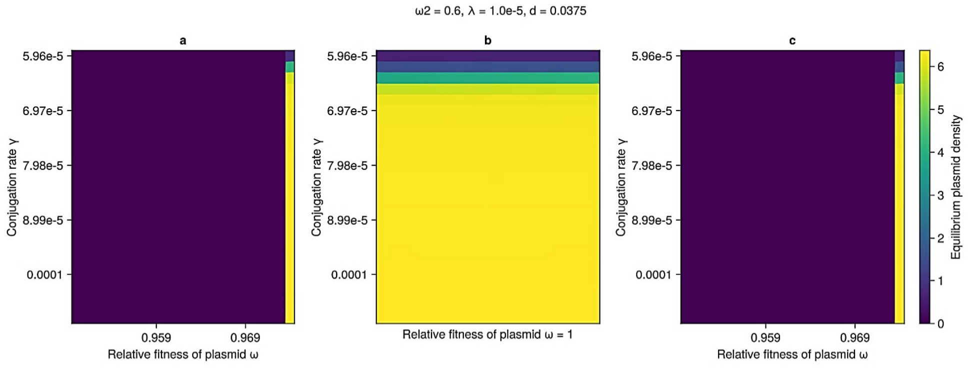
ω2 = 0.6, λ = 1.0e-5, d = 0.0375
a
b
c
Conjugation rate γ
5.96e-5
6.97e-5
7.98e-5
8.99e-5
0.0001
0.959
0.969
Relative fitness of plasmid ω
Relative fitness of plasmid ω = 1
Equilibrium plasmid density
0
1
2
3
4
5
6

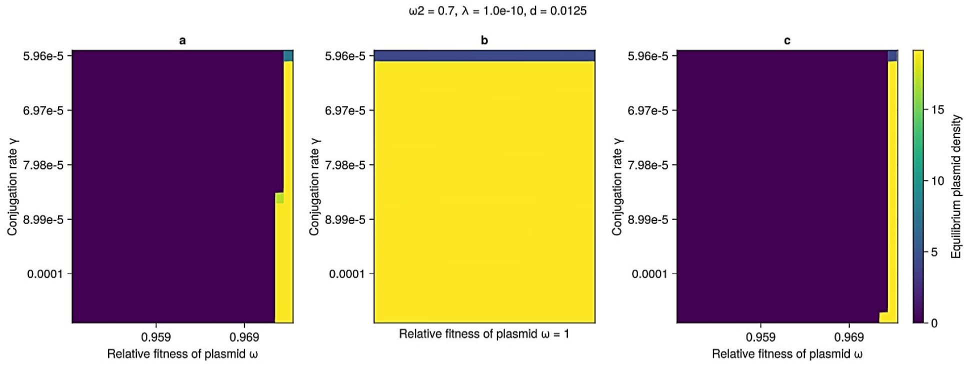
ω2 = 0.7, λ = 1.0e-10, d = 0.0125
a
b
c
Conjugation rate γ
5.96e-5
6.97e-5
7.98e-5
8.99e-5
0.0001
0.959
0.969
Relative fitness of plasmid ω
Relative fitness of plasmid ω = 1
Equilibrium plasmid density
0
5
10
15

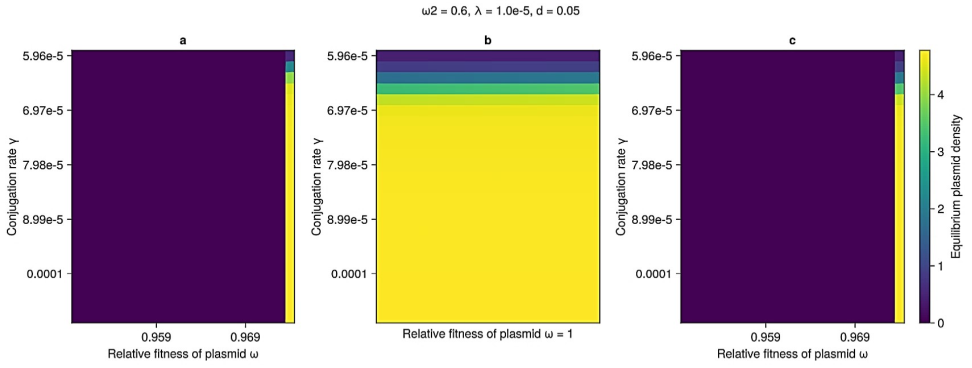
ω2 = 0.6, λ = 1.0e-5, d = 0.05
a
b
c
Conjugation rate γ
5.96e-5
6.97e-5
7.98e-5
8.99e-5
0.0001
0.959
0.969
Relative fitness of plasmid ω
Relative fitness of plasmid ω = 1
Equilibrium plasmid density
0
1
2
3
4

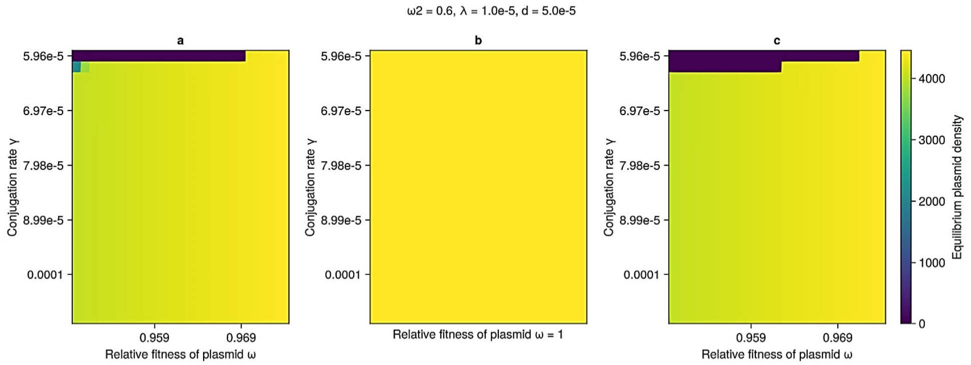
ω2 = 0.6, λ = 1.0e-5, d = 5.0e-5
a
b
c
Conjugation rate γ
5.96e-5
6.97e-5
7.98e-5
8.99e-5
0.0001
0.959
0.969
Relative fitness of plasmid ω
Relative fitness of plasmid ω = 1
Equilibrium plasmid density
0
1000
2000
3000
4000

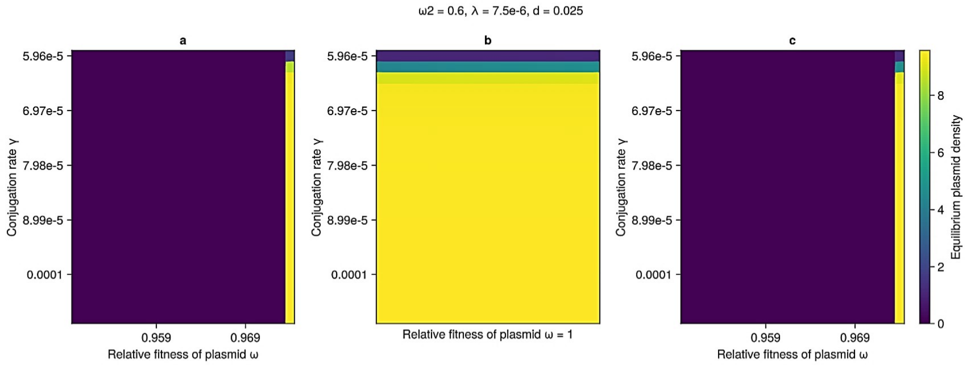
ω2 = 0.6, λ = 7.5e-6, d = 0.025
a
b
c
Conjugation rate γ
5.96e-5
6.97e-5
7.98e-5
8.99e-5
0.0001
0.959
0.969
Relative fitness of plasmid ω
Relative fitness of plasmid ω = 1
Equilibrium plasmid density
0
2
4
6
8

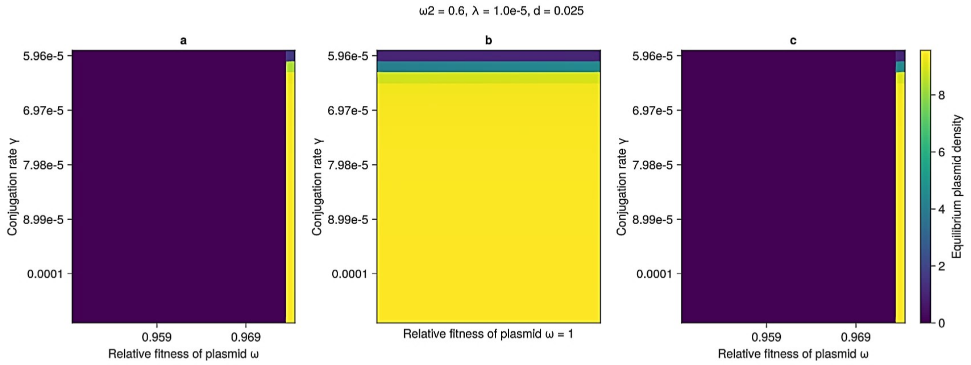
ω2 = 0.6, λ = 1.0e-5, d = 0.025
a
b
c
Conjugation rate γ
5.96e-5
6.97e-5
7.98e-5
8.99e-5
0.0001
0.959
0.969
Relative fitness of plasmid ω
Relative fitness of plasmid ω = 1
Equilibrium plasmid density
0
2
4
6
8

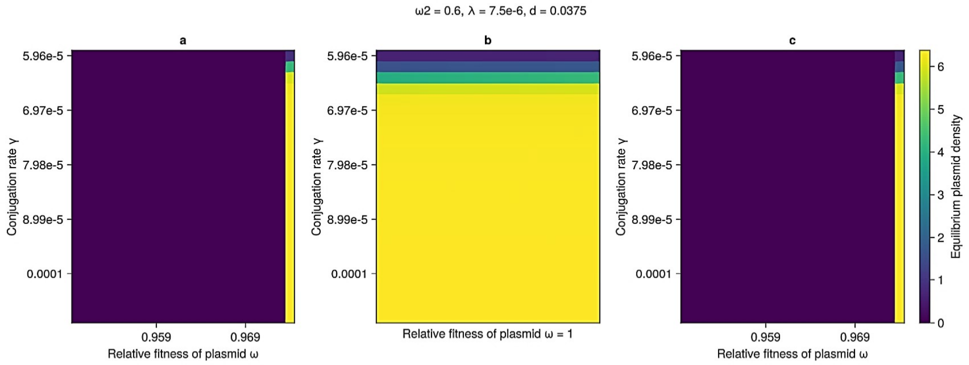
ω2 = 0.6, λ = 7.5e-6, d = 0.0375
a
b
c
Conjugation rate γ
5.96e-5
6.97e-5
7.98e-5
8.99e-5
0.0001
0.959
0.969
Relative fitness of plasmid ω
Relative fitness of plasmid ω = 1
Equilibrium plasmid density
0
1
2
3
4
5
6

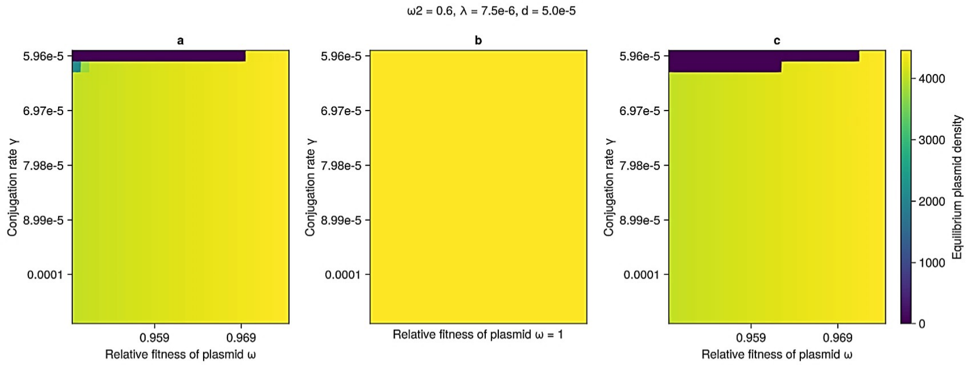
ω2 = 0.6, λ = 7.5e-6, d = 5.0e-5
a
b
c
Conjugation rate γ
5.96e-5
6.97e-5
7.98e-5
8.99e-5
0.0001
0.959
0.969
Relative fitness of plasmid ω
Relative fitness of plasmid ω = 1
Equilibrium plasmid density
0
1000
2000
3000
4000

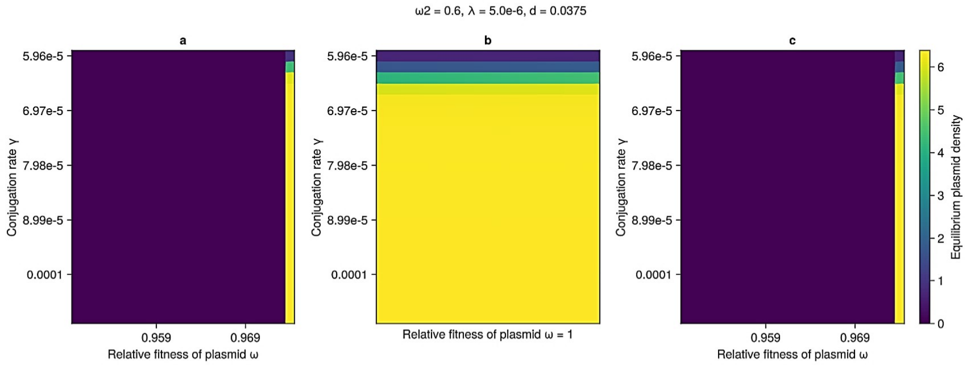
ω2 = 0.6, λ = 5.0e-6, d = 0.0375
a
b
c
Conjugation rate γ
5.96e-5
6.97e-5
7.98e-5
8.99e-5
0.0001
0.959
0.969
Relative fitness of plasmid ω
Relative fitness of plasmid ω = 1
Equilibrium plasmid density
0
1
2
3
4
5
6

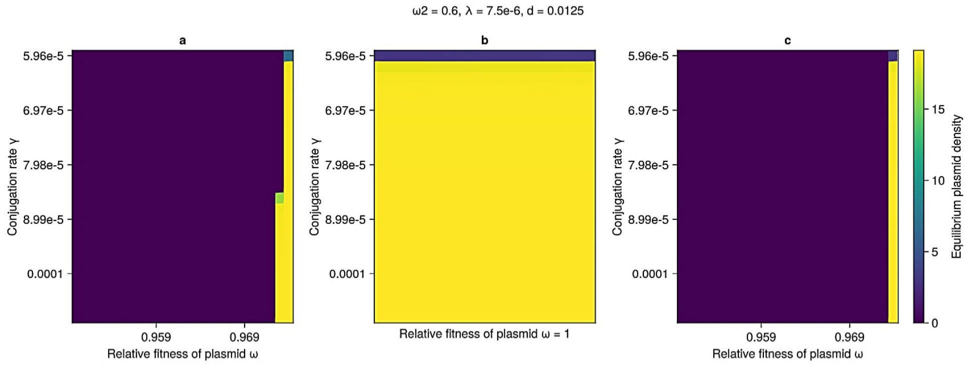
ω2 = 0.6, λ = 7.5e-6, d = 0.0125
a
b
c
Conjugation rate γ
5.96e-5
6.97e-5
7.98e-5
8.99e-5
0.0001
0.959
0.969
Relative fitness of plasmid ω
Relative fitness of plasmid ω = 1
Equilibrium plasmid density
0
5
10
15

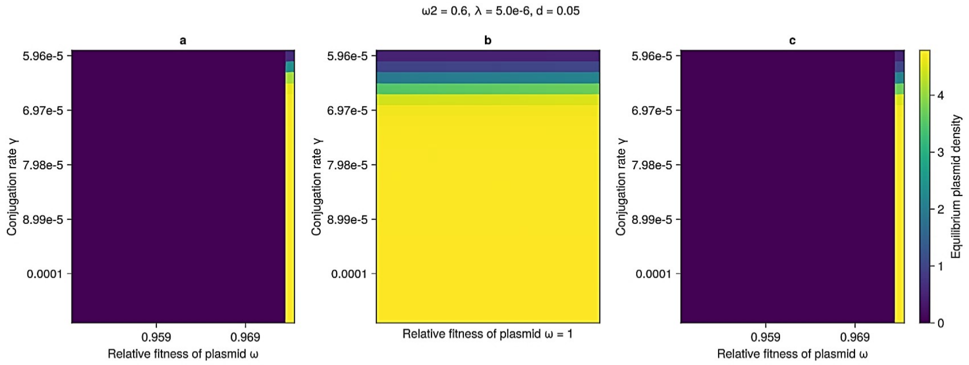
ω2 = 0.6, λ = 5.0e-6, d = 0.05
a
b
c
Conjugation rate γ
5.96e-5
6.97e-5
7.98e-5
8.99e-5
0.0001
0.959
0.969
Relative fitness of plasmid ω
Relative fitness of plasmid ω = 1
Equilibrium plasmid density
0
1
2
3
4

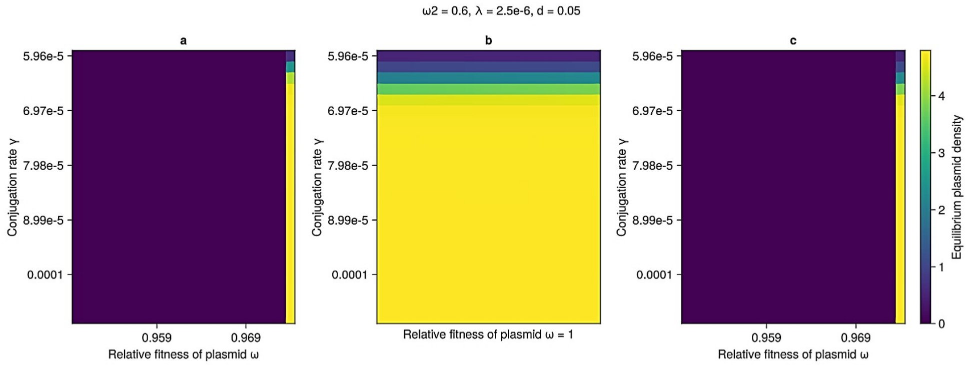
ω2 = 0.6, λ = 2.5e-6, d = 0.05
a
b
c
Conjugation rate γ
5.96e-5
6.97e-5
7.98e-5
8.99e-5
0.0001
0.959
0.969
Relative fitness of plasmid ω
Relative fitness of plasmid ω = 1
Equilibrium plasmid density
0
1
2
3
4

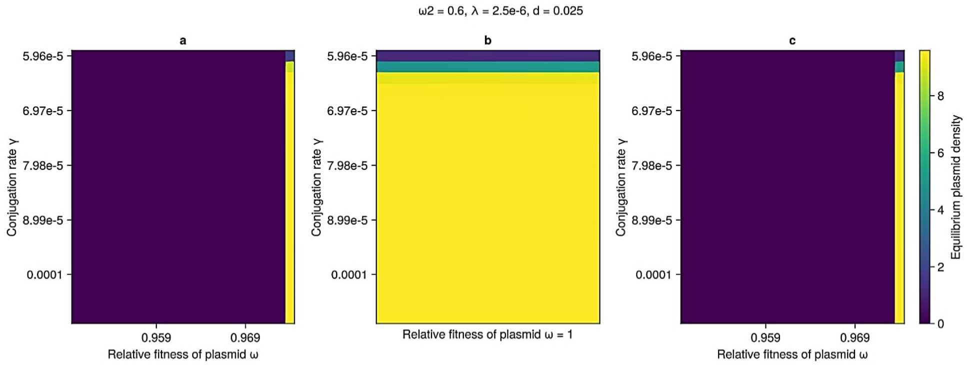
ω2 = 0.6, λ = 2.5e-6, d = 0.025
a
b
c
Conjugation rate γ
5.96e-5
6.97e-5
7.98e-5
8.99e-5
0.0001
0.959
0.969
Relative fitness of plasmid ω
Relative fitness of plasmid ω = 1
Equilibrium plasmid density
0
2
4
6
8

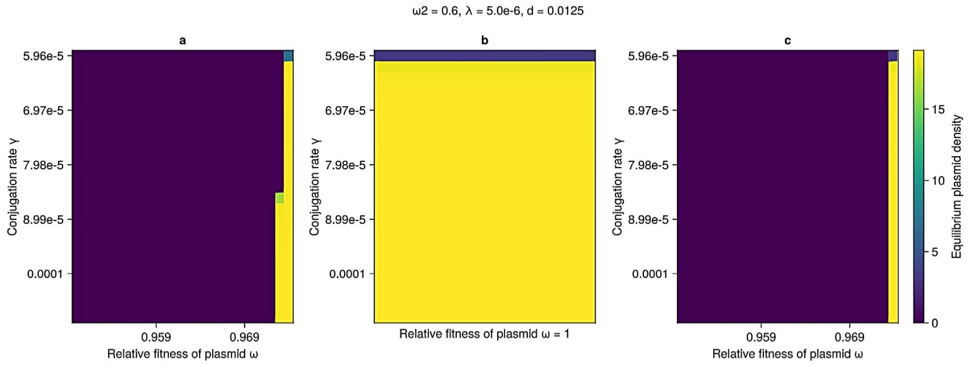
ω2 = 0.6, λ = 5.0e-6, d = 0.0125
a
b
c
Conjugation rate γ
5.96e-5
6.97e-5
7.98e-5
8.99e-5
0.0001
0.959
0.969
Relative fitness of plasmid ω
Relative fitness of plasmid ω = 1
Equilibrium plasmid density
0
5
10
15

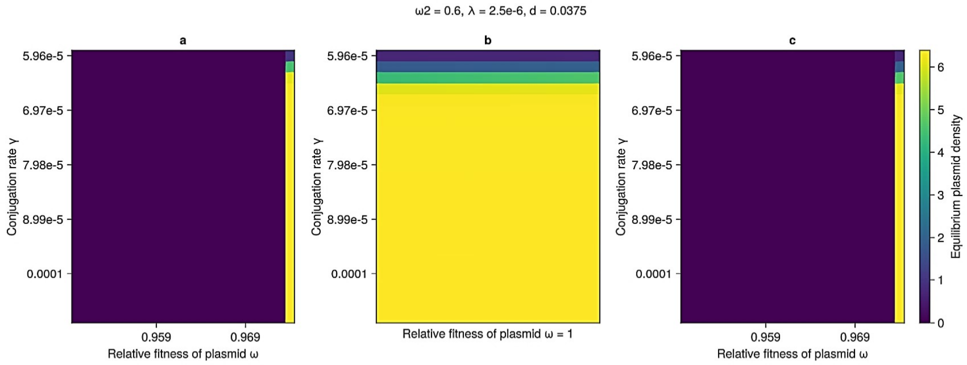
ω2 = 0.6, λ = 2.5e-6, d = 0.0375
a
b
c
Conjugation rate γ
5.96e-5
6.97e-5
7.98e-5
8.99e-5
0.0001
0.959
0.969
Relative fitness of plasmid ω
Relative fitness of plasmid ω = 1
Equilibrium plasmid density
0
1
2
3
4
5
6

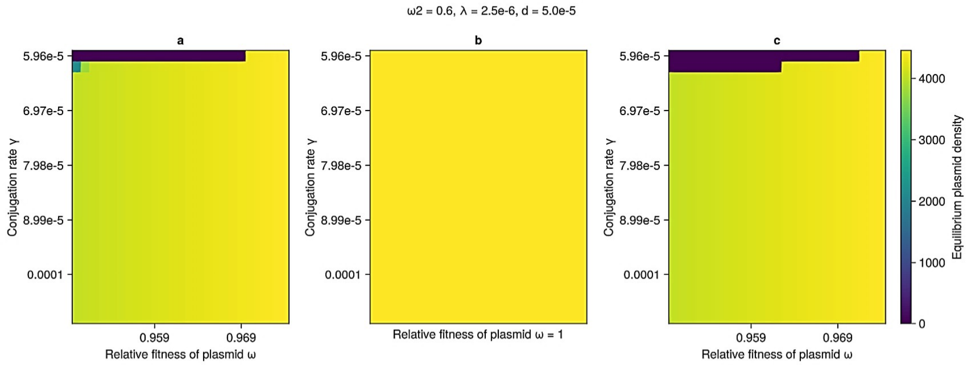
ω2 = 0.6, λ = 2.5e-6, d = 5.0e-5
a
b
c
Conjugation rate γ
5.96e-5
6.97e-5
7.98e-5
8.99e-5
0.0001
0.959
0.969
Relative fitness of plasmid ω
Relative fitness of plasmid ω = 1
Equilibrium plasmid density
0
1000
2000
3000
4000
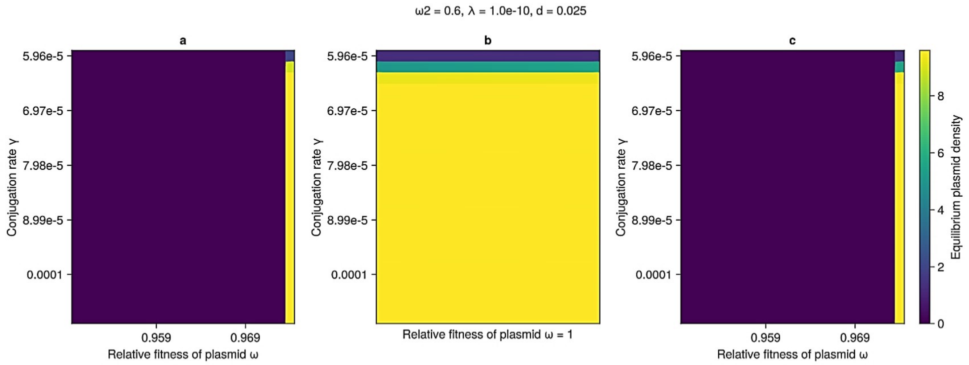
ω2 = 0.6, λ = 1.0e-10, d = 0.025
a
b
c
Conjugation rate γ
5.96e-5
6.97e-5
7.98e-5
8.99e-5
0.0001
0.959
0.969
Relative fitness of plasmid ω
Relative fitness of plasmid ω = 1
Equilibrium plasmid density
0
2
4
6
8
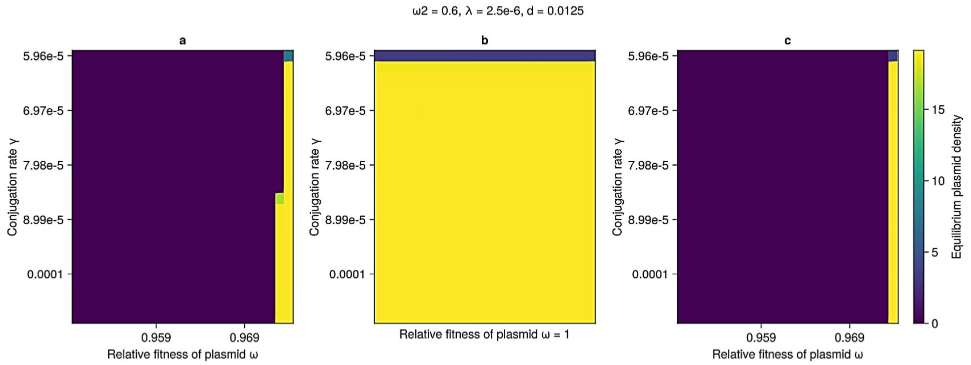
ω2 = 0.6, λ = 2.5e-6, d = 0.0125
a
b
c
Conjugation rate γ
5.96e-5
6.97e-5
7.98e-5
8.99e-5
0.0001
0.959
0.969
Relative fitness of plasmid ω
Relative fitness of plasmid ω = 1
Equilibrium plasmid density
0
5
10
15
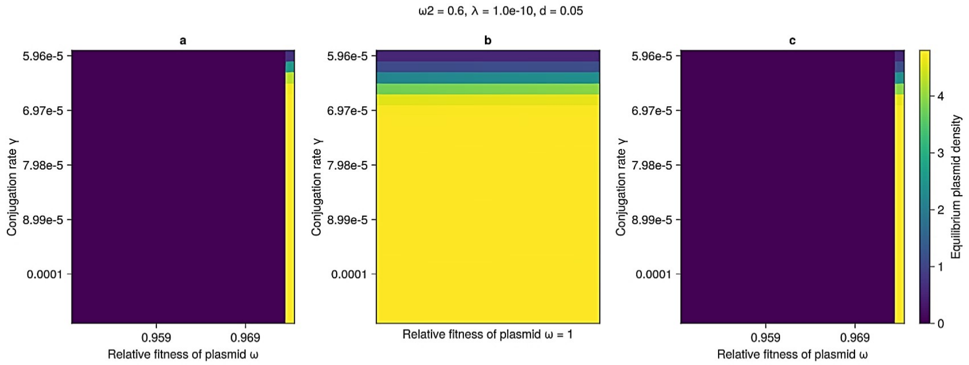
ω2 = 0.6, λ = 1.0e-10, d = 0.05
a
b
c
Conjugation rate γ
5.96e-5
6.97e-5
7.98e-5
8.99e-5
0.0001
0.959
0.969
Relative fitness of plasmid ω
Relative fitness of plasmid ω = 1
Equilibrium plasmid density
0
1
2
3
4

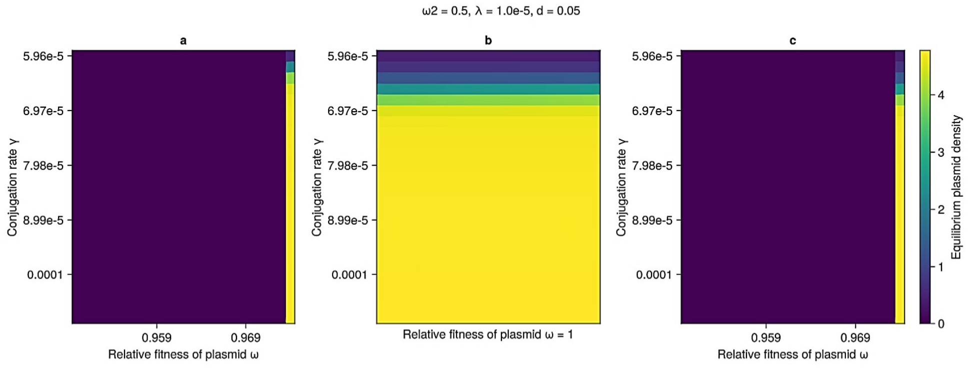
ω2 = 0.5, λ = 1.0e-5, d = 0.05
a
b
c
Conjugation rate γ
5.96e-5
6.97e-5
7.98e-5
8.99e-5
0.0001
0.959
0.969
Relative fitness of plasmid ω
Relative fitness of plasmid ω = 1
Equilibrium plasmid density
0
1
2
3
4

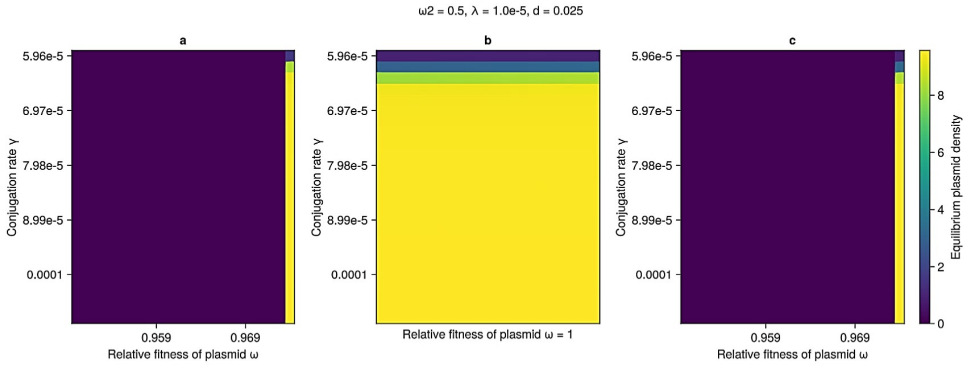
ω2 = 0.5, λ = 1.0e-5, d = 0.025
a
b
c
Conjugation rate γ
5.96e-5
6.97e-5
7.98e-5
8.99e-5
0.0001
0.959
0.969
Relative fitness of plasmid ω
Relative fitness of plasmid ω = 1
Equilibrium plasmid density
0
2
4
6
8

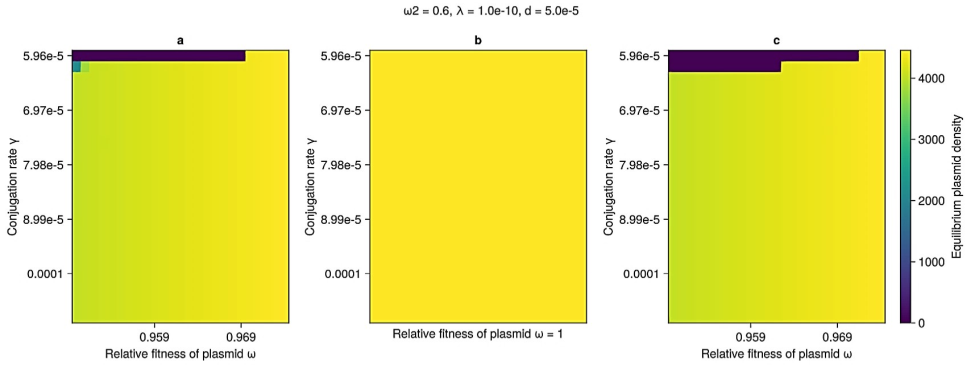
ω2 = 0.6, λ = 1.0e-10, d = 5.0e-5
a
b
c
Conjugation rate γ
5.96e-5
6.97e-5
7.98e-5
8.99e-5
0.0001
0.959
0.969
Relative fitness of plasmid ω
Relative fitness of plasmid ω = 1
Equilibrium plasmid density
0
1000
2000
3000
4000

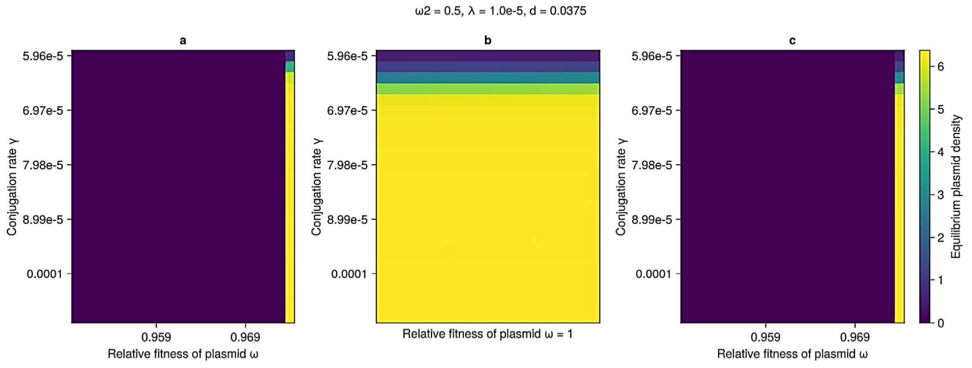
ω2 = 0.5, λ = 1.0e-5, d = 0.0375
a
b
c
Conjugation rate γ
5.96e-5
6.97e-5
7.98e-5
8.99e-5
0.0001
0.959
0.969
Relative fitness of plasmid ω
Relative fitness of plasmid ω = 1
Equilibrium plasmid density
0
1
2
3
4
5
6

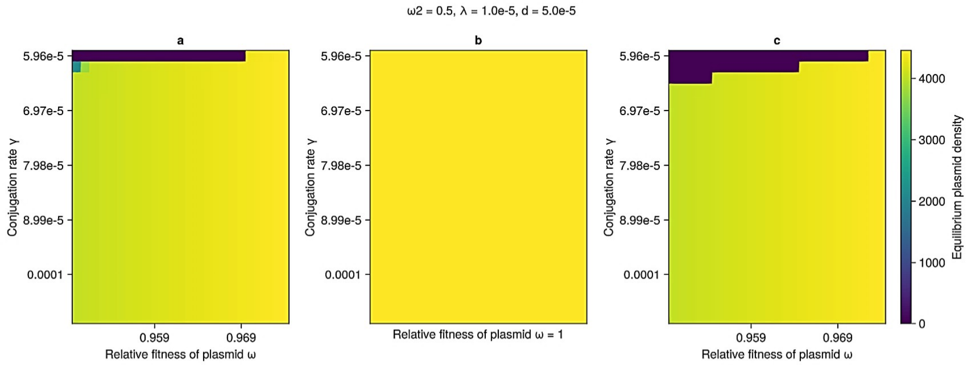
ω2 = 0.5, λ = 1.0e-5, d = 5.0e-5
a
b
c
Conjugation rate γ
5.96e-5
6.97e-5
7.98e-5
8.99e-5
0.0001
0.959
0.969
Relative fitness of plasmid ω
Relative fitness of plasmid ω = 1
0
1000
2000
3000
4000
Equilibrium plasmid density

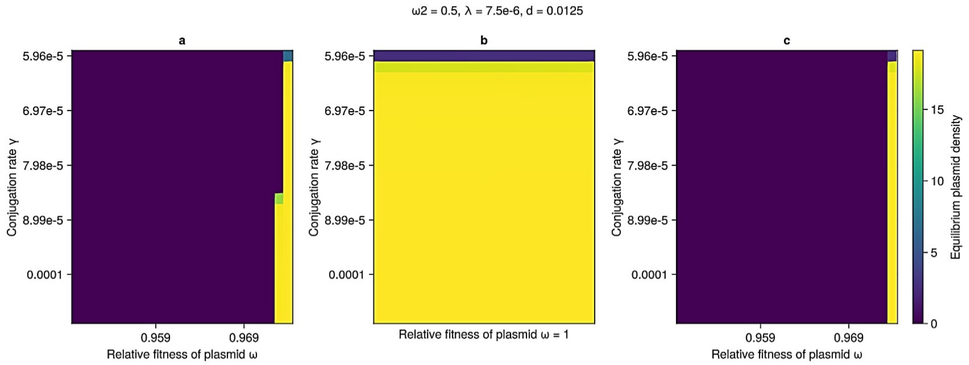
ω2 = 0.5, λ = 7.5e-6, d = 0.0125
a
b
c
Conjugation rate γ
5.96e-5
6.97e-5
7.98e-5
8.99e-5
0.0001
0.959
0.969
Relative fitness of plasmid ω
Relative fitness of plasmid ω = 1
0
5
10
15
Equilibrium plasmid density

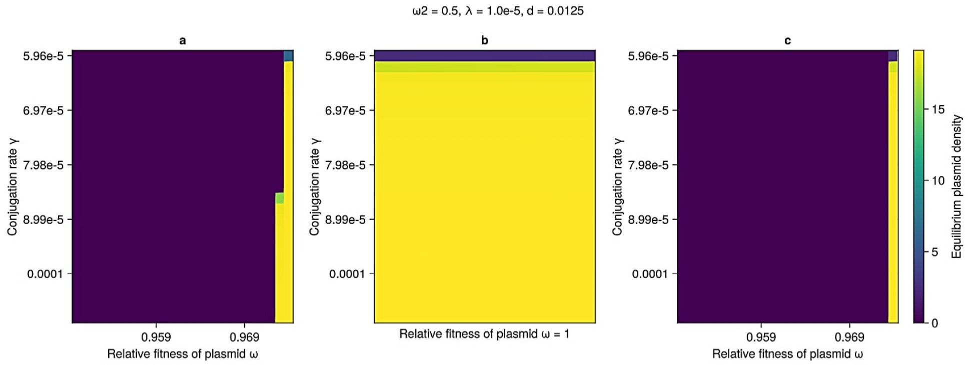
ω2 = 0.5, λ = 1.0e-5, d = 0.0125
a
b
c
Conjugation rate γ
5.96e-5
6.97e-5
7.98e-5
8.99e-5
0.0001
0.959
0.969
Relative fitness of plasmid ω
Relative fitness of plasmid ω = 1
0
5
10
15
Equilibrium plasmid density

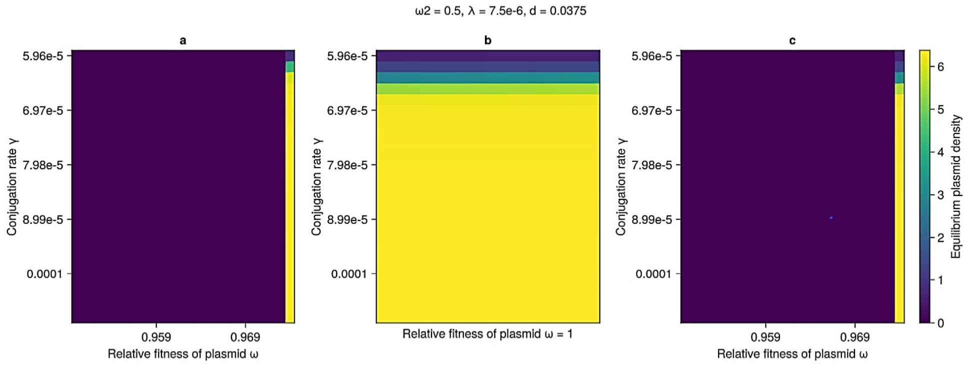
ω2 = 0.5, λ = 7.5e-6, d = 0.0375
a
b
c
Conjugation rate γ
5.96e-5
6.97e-5
7.98e-5
8.99e-5
0.0001
0.959
0.969
Relative fitness of plasmid ω
Relative fitness of plasmid ω = 1
0
1
2
3
4
5
6
Equilibrium plasmid density

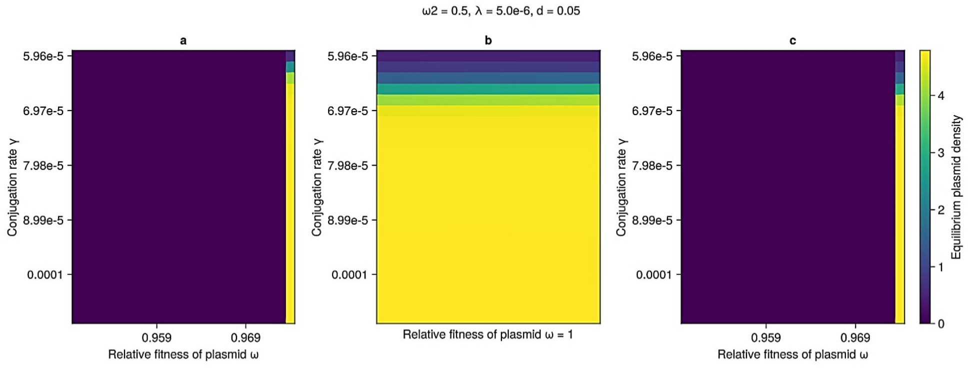
ω2 = 0.5, λ = 5.0e-6, d = 0.05
a
b
c
Conjugation rate γ
5.96e-5
6.97e-5
7.98e-5
8.99e-5
0.0001
0.959
0.969
Relative fitness of plasmid ω
Relative fitness of plasmid ω = 1
Equilibrium plasmid density
0
1
2
3
4

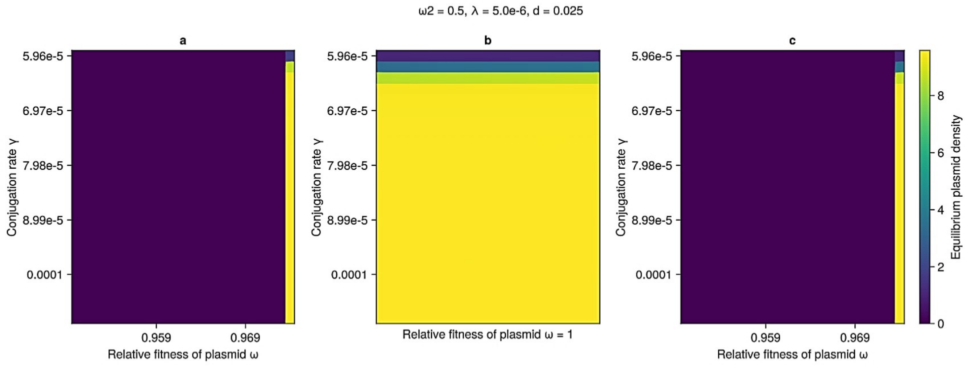
ω2 = 0.5, λ = 5.0e-6, d = 0.025
a
b
c
Conjugation rate γ
5.96e-5
6.97e-5
7.98e-5
8.99e-5
0.0001
0.959
0.969
Relative fitness of plasmid ω
Relative fitness of plasmid ω = 1
Equilibrium plasmid density
0
2
4
6
8

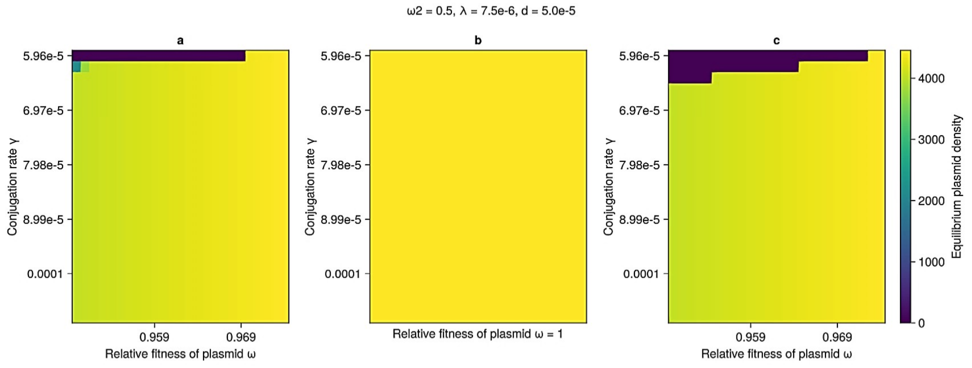
ω2 = 0.5, λ = 7.5e-6, d = 5.0e-5
a
b
c
Conjugation rate γ
5.96e-5
6.97e-5
7.98e-5
8.99e-5
0.0001
0.959
0.969
Relative fitness of plasmid ω
Relative fitness of plasmid ω = 1
Equilibrium plasmid density
0
1000
2000
3000
4000

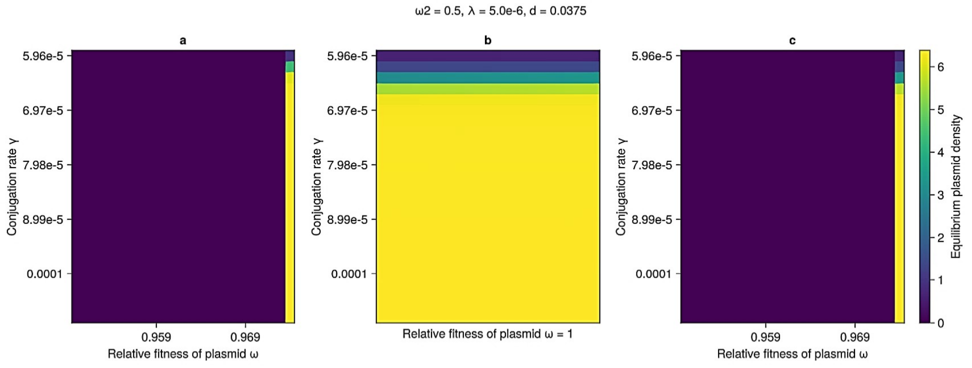
ω2 = 0.5, λ = 5.0e-6, d = 0.0375
a
b
c
Conjugation rate γ
5.96e-5
6.97e-5
7.98e-5
8.99e-5
0.0001
0.959
0.969
Relative fitness of plasmid ω
Relative fitness of plasmid ω = 1
Equilibrium plasmid density
0
1
2
3
4
5
6

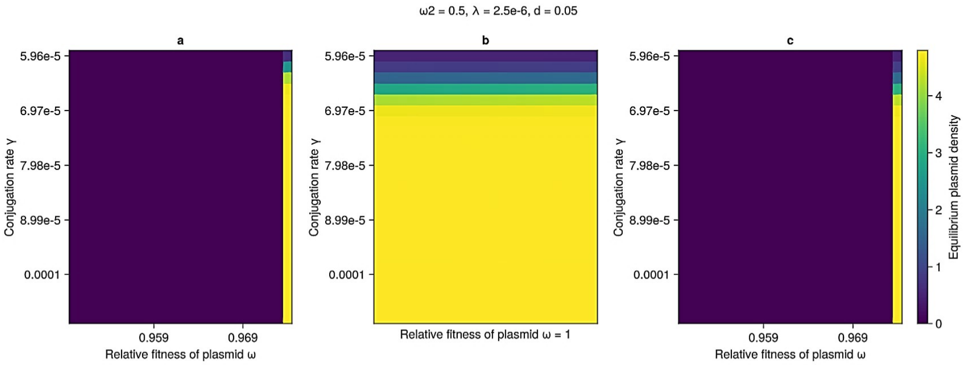
ω2 = 0.5, λ = 2.5e-6, d = 0.05
a
b
c
5.96e-5
6.97e-5
7.98e-5
8.99e-5
0.0001
Conjugation rate γ
0.959
0.969
Relative fitness of plasmid ω
Relative fitness of plasmid ω = 1
Equilibrium plasmid density
0
1
2
3
4

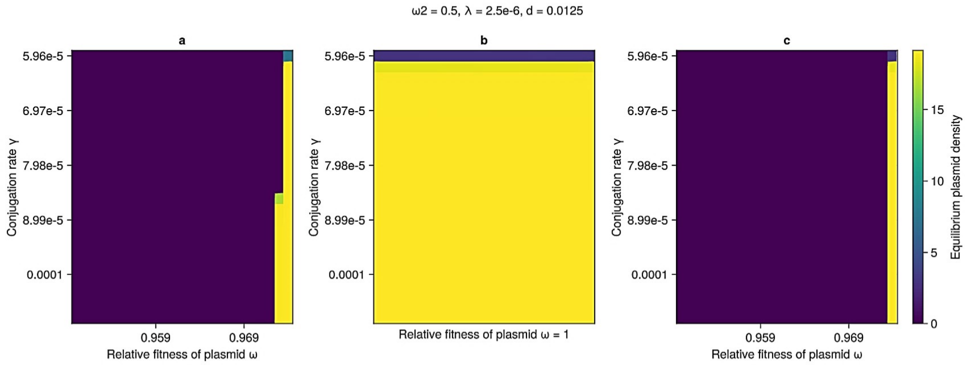
ω2 = 0.5, λ = 2.5e-6, d = 0.0125
a
b
c
5.96e-5
6.97e-5
7.98e-5
8.99e-5
0.0001
Conjugation rate γ
0.959
0.969
Relative fitness of plasmid ω
Relative fitness of plasmid ω = 1
Equilibrium plasmid density
0
5
10
15

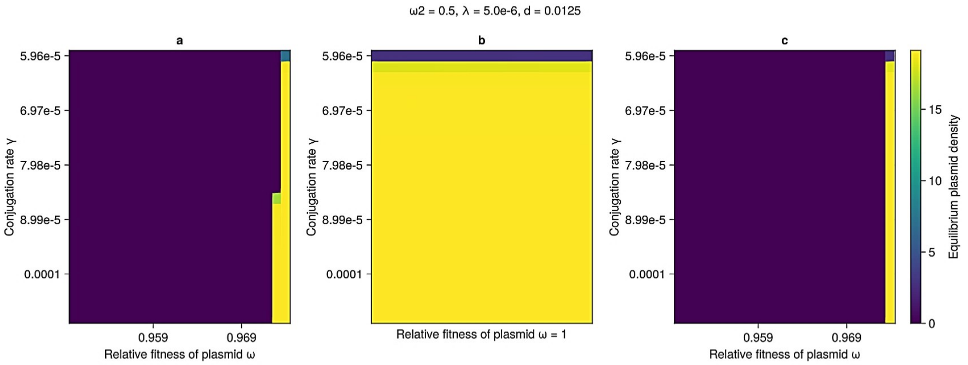
ω2 = 0.5, λ = 5.0e-6, d = 0.0125
a
b
c
5.96e-5
6.97e-5
7.98e-5
8.99e-5
0.0001
Conjugation rate γ
0.959
0.969
Relative fitness of plasmid ω
Relative fitness of plasmid ω = 1
Equilibrium plasmid density
0
5
10
15

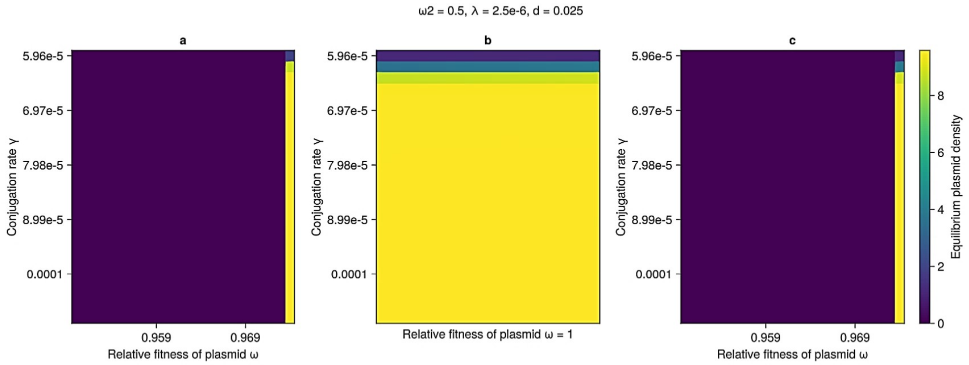
ω2 = 0.5, λ = 2.5e-6, d = 0.025
a
b
c
5.96e-5
6.97e-5
7.98e-5
8.99e-5
0.0001
Conjugation rate γ
0.959
0.969
Relative fitness of plasmid ω
Relative fitness of plasmid ω = 1
Equilibrium plasmid density
0
2
4
6
8

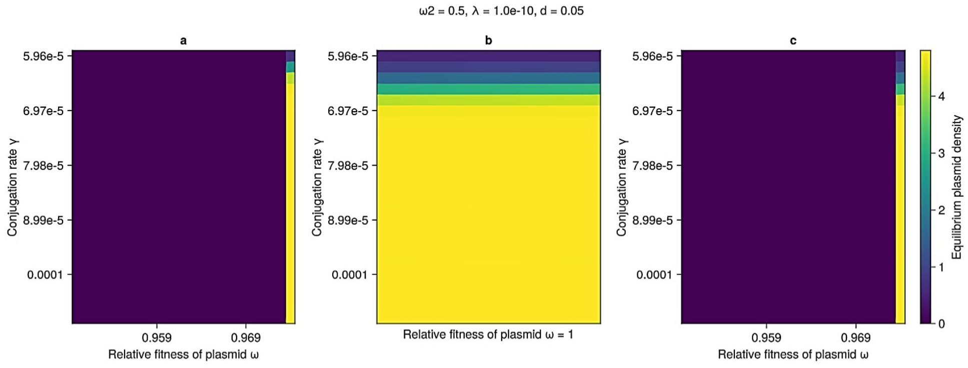
ω2 = 0.5, λ = 1.0e-10, d = 0.05
a
b
c
Conjugation rate γ
5.96e-5
6.97e-5
7.98e-5
8.99e-5
0.0001
0.959
0.969
Relative fitness of plasmid ω
Relative fitness of plasmid ω = 1
Equilibrium plasmid density
0
1
2
3
4

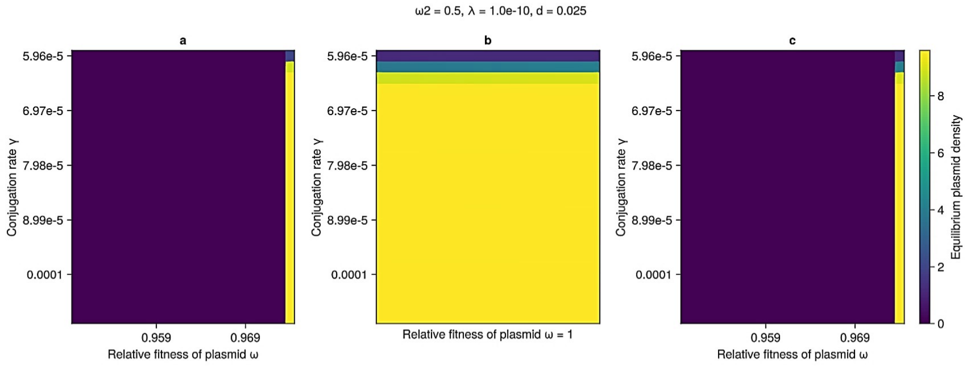
ω2 = 0.5, λ = 1.0e-10, d = 0.025
a
b
c
Conjugation rate γ
5.96e-5
6.97e-5
7.98e-5
8.99e-5
0.0001
0.959
0.969
Relative fitness of plasmid ω
Relative fitness of plasmid ω = 1
Equilibrium plasmid density
0
2
4
6
8

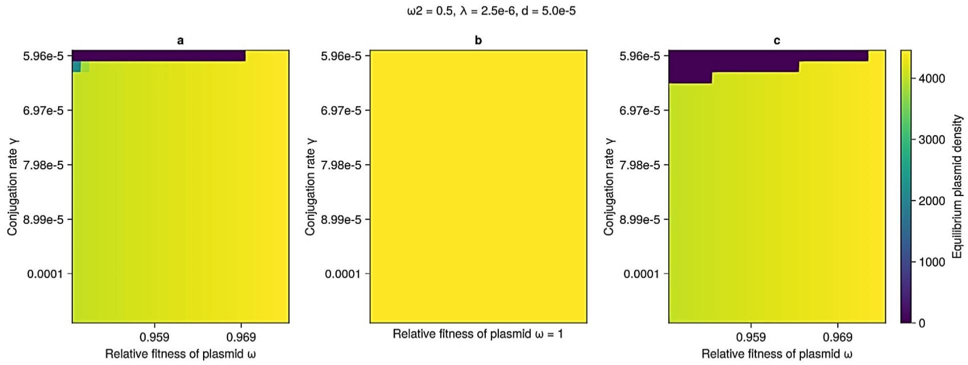
ω2 = 0.5, λ = 2.5e-6, d = 5.0e-5
a
b
c
Conjugation rate γ
5.96e-5
6.97e-5
7.98e-5
8.99e-5
0.0001
0.959
0.969
Relative fitness of plasmid ω
Relative fitness of plasmid ω = 1
Equilibrium plasmid density
0
1000
2000
3000
4000

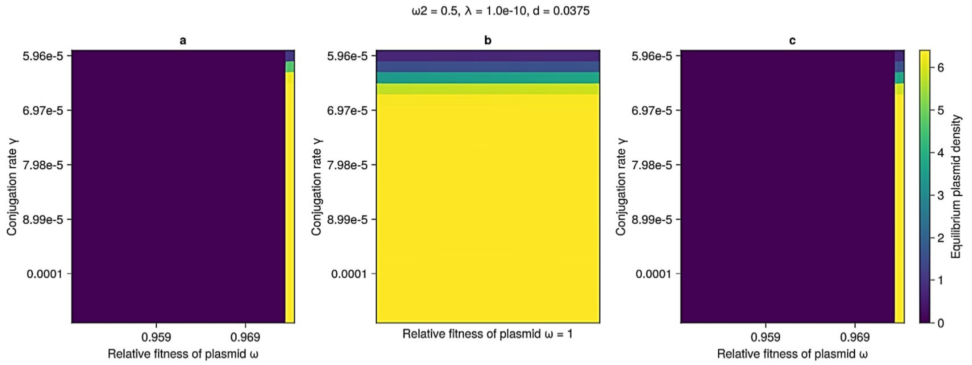
ω2 = 0.5, λ = 1.0e-10, d = 0.0375
a
b
c
Conjugation rate γ
5.96e-5
6.97e-5
7.98e-5
8.99e-5
0.0001
0.959
0.969
Relative fitness of plasmid ω
Relative fitness of plasmid ω = 1
Equilibrium plasmid density
0
1
2
3
4
5
6

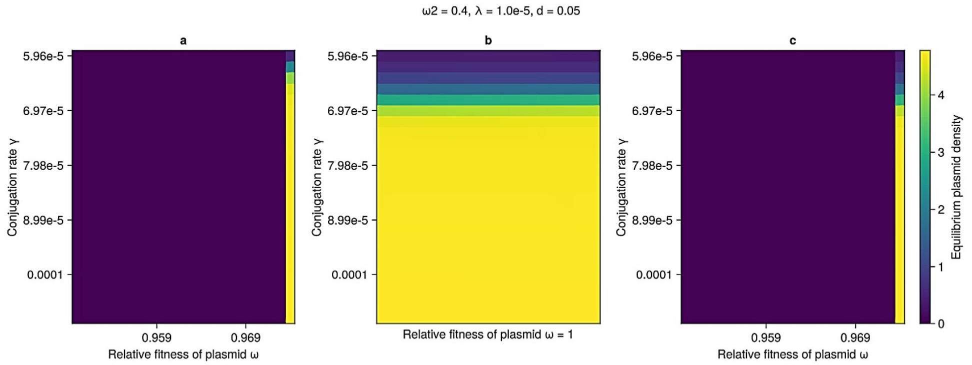
ω2 = 0.4, λ = 1.0e-5, d = 0.05
a
b
c
Conjugation rate γ
5.96e-5
6.97e-5
7.98e-5
8.99e-5
0.0001
0.959
0.969
Relative fitness of plasmid ω
Relative fitness of plasmid ω = 1
Equilibrium plasmid density
0
1
2
3
4

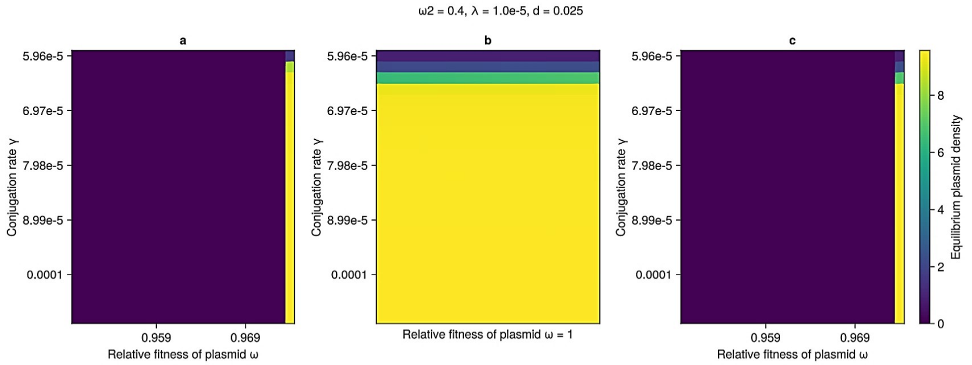
ω2 = 0.4, λ = 1.0e-5, d = 0.025
a
b
c
Conjugation rate γ
5.96e-5
6.97e-5
7.98e-5
8.99e-5
0.0001
0.959
0.969
Relative fitness of plasmid ω
Relative fitness of plasmid ω = 1
Equilibrium plasmid density
0
2
4
6
8

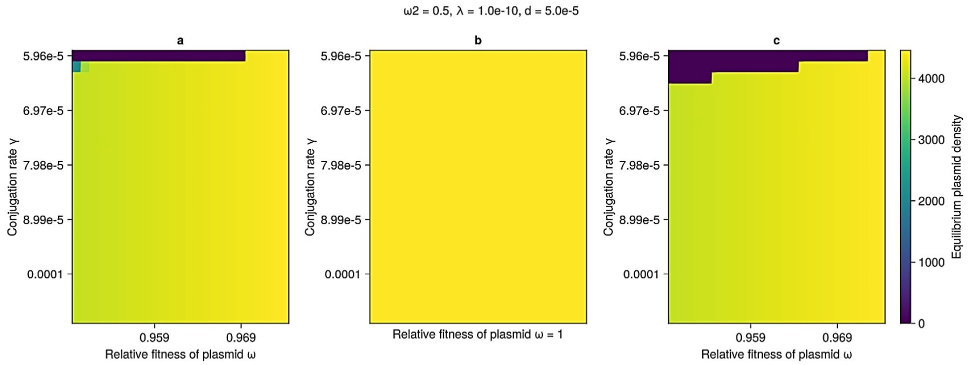
ω2 = 0.5, λ = 1.0e-10, d = 5.0e-5
a
b
c
Conjugation rate γ
5.96e-5
6.97e-5
7.98e-5
8.99e-5
0.0001
0.959
0.969
Relative fitness of plasmid ω
Relative fitness of plasmid ω = 1
Equilibrium plasmid density
0
1000
2000
3000
4000

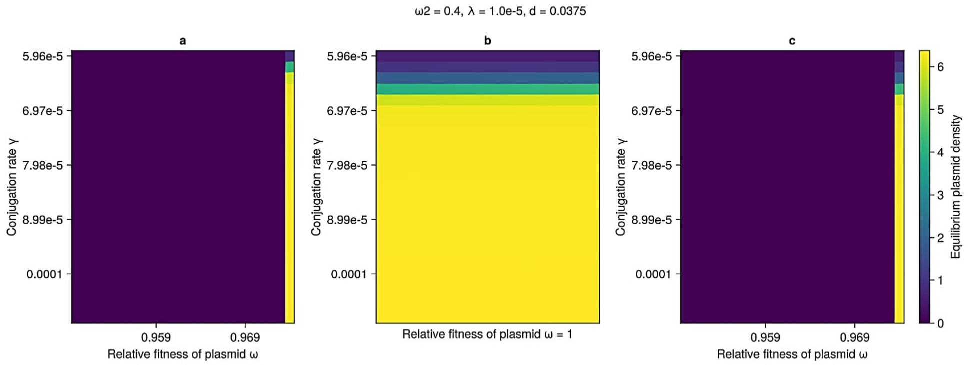
ω2 = 0.4, λ = 1.0e-5, d = 0.0375
a
b
c
Conjugation rate γ
5.96e-5
6.97e-5
7.98e-5
8.99e-5
0.0001
0.959
0.969
Relative fitness of plasmid ω
Relative fitness of plasmid ω = 1
Equilibrium plasmid density
0
1
2
3
4
5
6

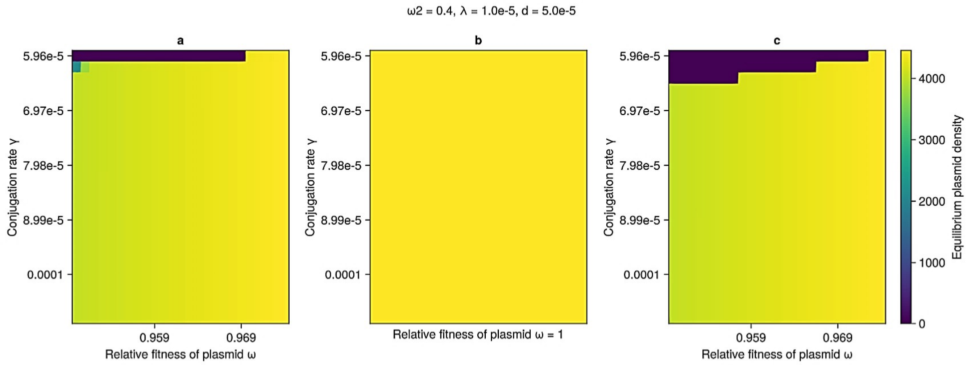
ω2 = 0.4, λ = 1.0e-5, d = 5.0e-5
a
b
c
Conjugation rate γ
5.96e-5
6.97e-5
7.98e-5
8.99e-5
0.0001
0.959
0.969
Relative fitness of plasmid ω
Relative fitness of plasmid ω = 1
Equilibrium plasmid density
0
1000
2000
3000
4000

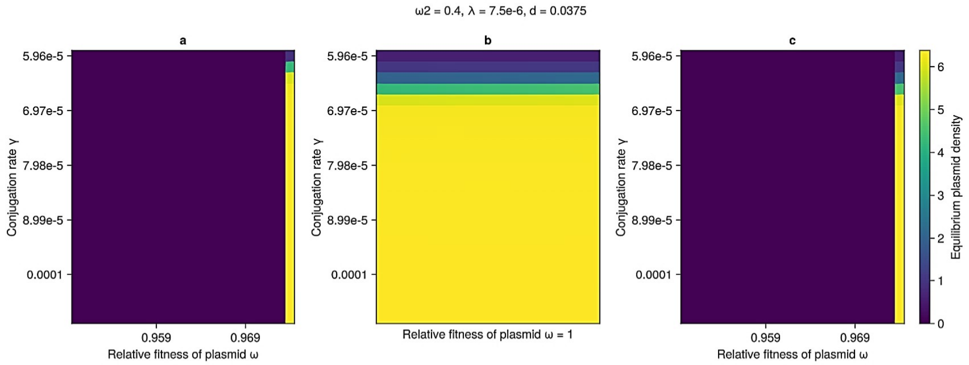
ω2 = 0.4, λ = 7.5e-6, d = 0.0375
a
b
c
Conjugation rate γ
5.96e-5
6.97e-5
7.98e-5
8.99e-5
0.0001
0.959
0.969
Relative fitness of plasmid ω
Relative fitness of plasmid ω = 1
Equilibrium plasmid density
0
1
2
3
4
5
6

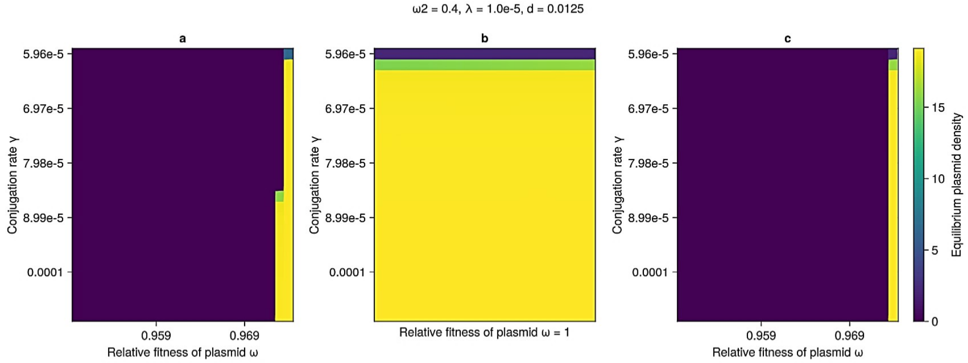
ω2 = 0.4, λ = 1.0e-5, d = 0.0125
a
b
c
Conjugation rate γ
5.96e-5
6.97e-5
7.98e-5
8.99e-5
0.0001
0.959
0.969
Relative fitness of plasmid ω
Relative fitness of plasmid ω = 1
Equilibrium plasmid density
0
5
10
15

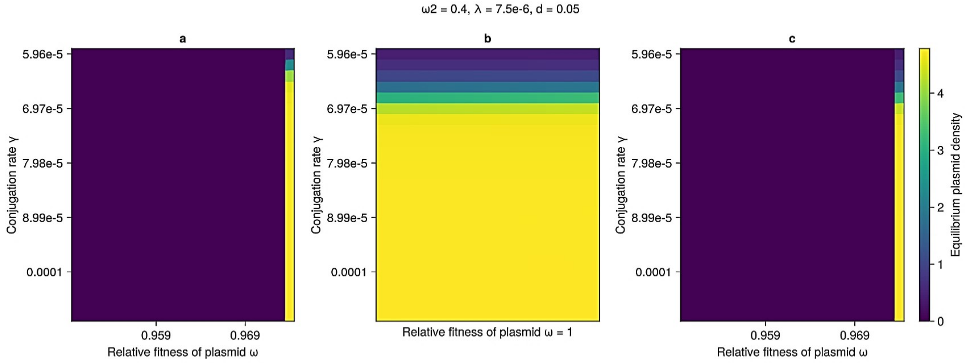
ω2 = 0.4, λ = 7.5e-6, d = 0.05
a
b
c
Conjugation rate γ
5.96e-5
6.97e-5
7.98e-5
8.99e-5
0.0001
0.959
0.969
Relative fitness of plasmid ω
Relative fitness of plasmid ω = 1
Equilibrium plasmid density
0
1
2
3
4

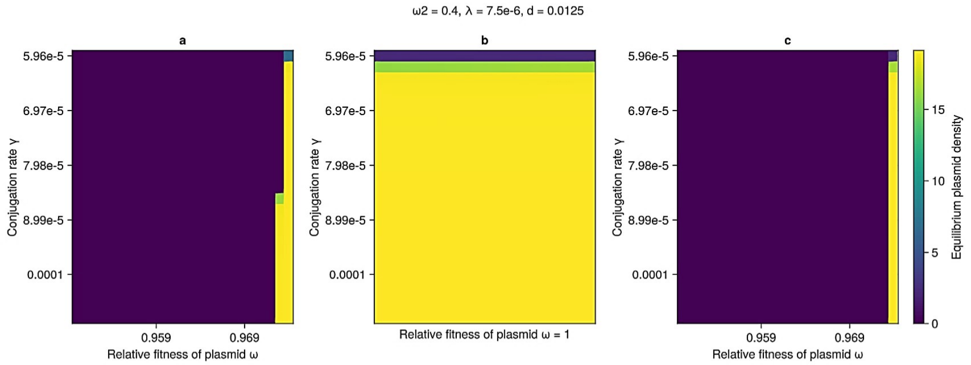
ω2 = 0.4, λ = 7.5e-6, d = 0.0125
a
b
c
Conjugation rate γ
5.96e-5
6.97e-5
7.98e-5
8.99e-5
0.0001
0.959
0.969
Relative fitness of plasmid ω
Relative fitness of plasmid ω = 1
Equilibrium plasmid density
0
5
10
15

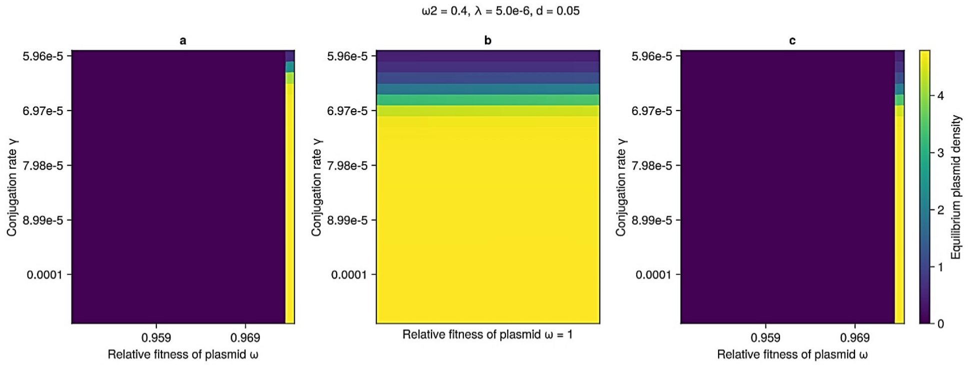
ω2 = 0.4, λ = 5.0e-6, d = 0.05
a
b
c
Conjugation rate γ
5.96e-5
6.97e-5
7.98e-5
8.99e-5
0.0001
0.959
0.969
Relative fitness of plasmid ω
Relative fitness of plasmid ω = 1
Equilibrium plasmid density
0
1
2
3
4

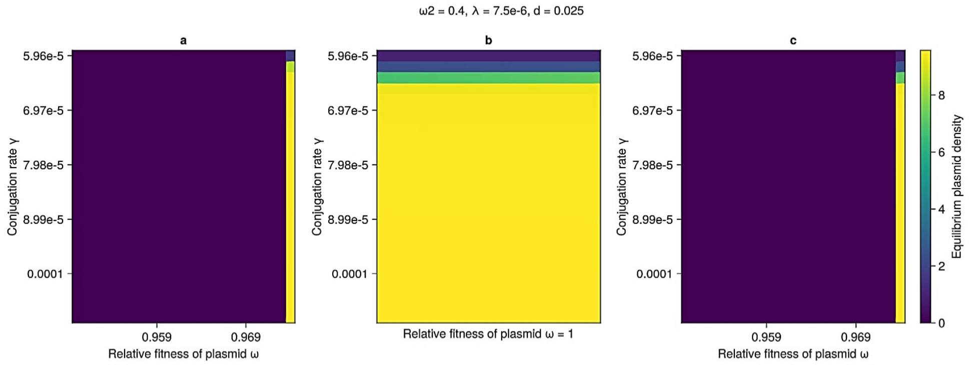
ω2 = 0.4, λ = 7.5e-6, d = 0.025
a
b
c
Conjugation rate γ
5.96e-5
6.97e-5
7.98e-5
8.99e-5
0.0001
0.959
0.969
Relative fitness of plasmid ω
Relative fitness of plasmid ω = 1
Equilibrium plasmid density
0
2
4
6
8

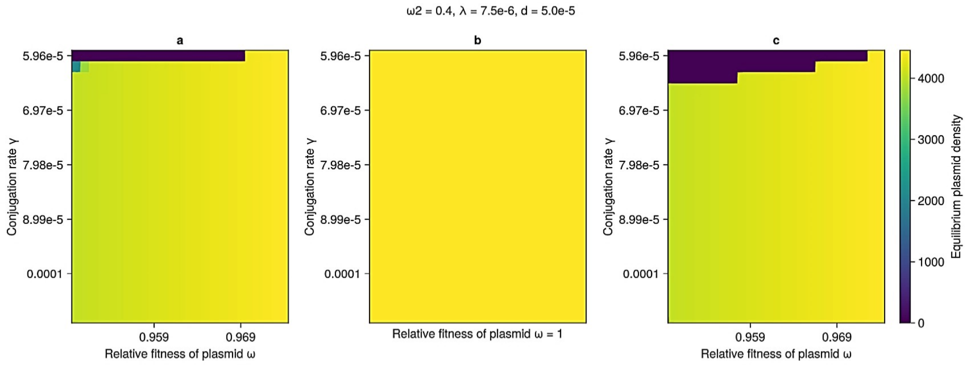
ω2 = 0.4, λ = 7.5e-6, d = 5.0e-5
a
b
c
Conjugation rate γ
5.96e-5
6.97e-5
7.98e-5
8.99e-5
0.0001
0.959
0.969
Relative fitness of plasmid ω
Relative fitness of plasmid ω = 1
Equilibrium plasmid density
0
1000
2000
3000
4000

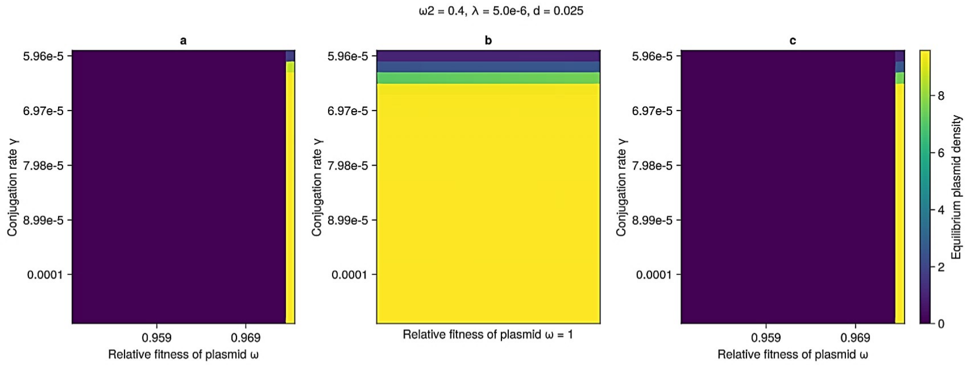
ω2 = 0.4, λ = 5.0e-6, d = 0.025
a
b
c
Conjugation rate γ
5.96e-5
6.97e-5
7.98e-5
8.99e-5
0.0001
0.959
0.969
Relative fitness of plasmid ω
Relative fitness of plasmid ω = 1
Equilibrium plasmid density
0
2
4
6
8

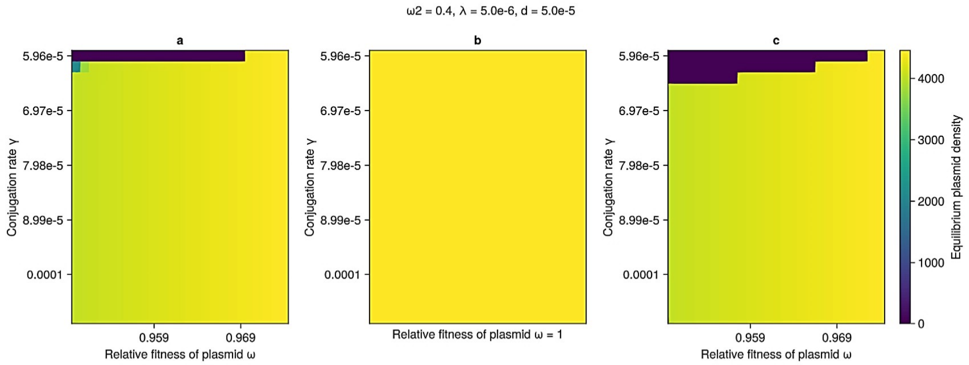
ω2 = 0.4, λ = 5.0e-6, d = 5.0e-5
a
b
c
Conjugation rate γ
5.96e-5
6.97e-5
7.98e-5
8.99e-5
0.0001
0.959
0.969
Relative fitness of plasmid ω
Relative fitness of plasmid ω = 1
Equilibrium plasmid density
0
1000
2000
3000
4000

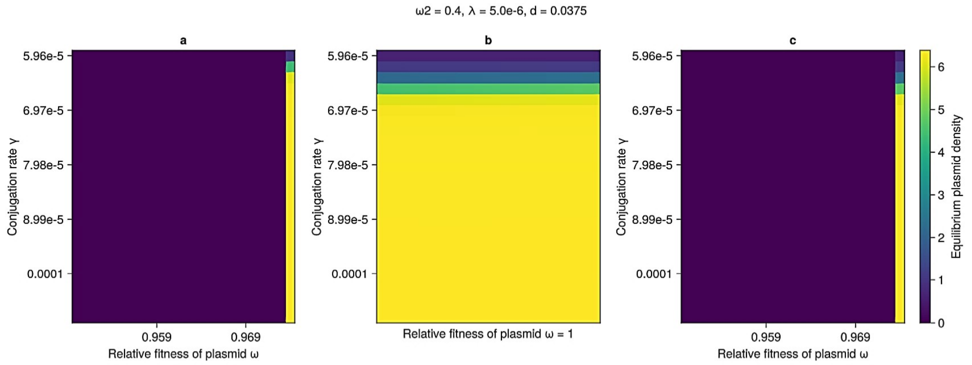
ω2 = 0.4, λ = 5.0e-6, d = 0.0375
a
b
c
Conjugation rate γ
5.96e-5
6.97e-5
7.98e-5
8.99e-5
0.0001
0.959
0.969
Relative fitness of plasmid ω
Relative fitness of plasmid ω = 1
Equilibrium plasmid density
0
1
2
3
4
5
6

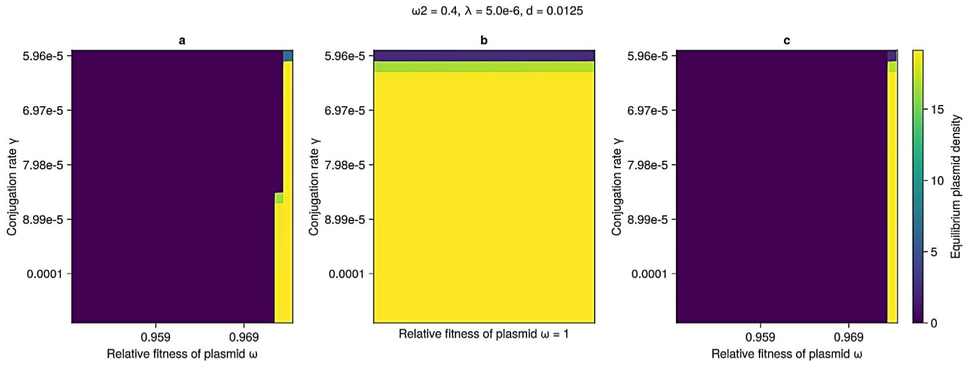
ω2 = 0.4, λ = 5.0e-6, d = 0.0125
a
b
c
Conjugation rate γ
5.96e-5
6.97e-5
7.98e-5
8.99e-5
0.0001
0.959
0.969
Relative fitness of plasmid ω
Relative fitness of plasmid ω = 1
Equilibrium plasmid density
0
5
10
15

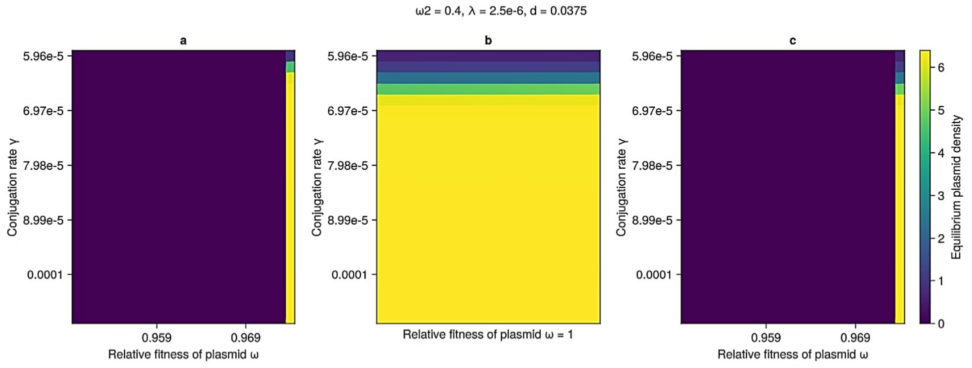
ω2 = 0.4, λ = 2.5e-6, d = 0.0375
a
b
c
Conjugation rate γ
5.96e-5
6.97e-5
7.98e-5
8.99e-5
0.0001
0.959
0.969
Relative fitness of plasmid ω
Relative fitness of plasmid ω = 1
Equilibrium plasmid density
0
1
2
3
4
5
6

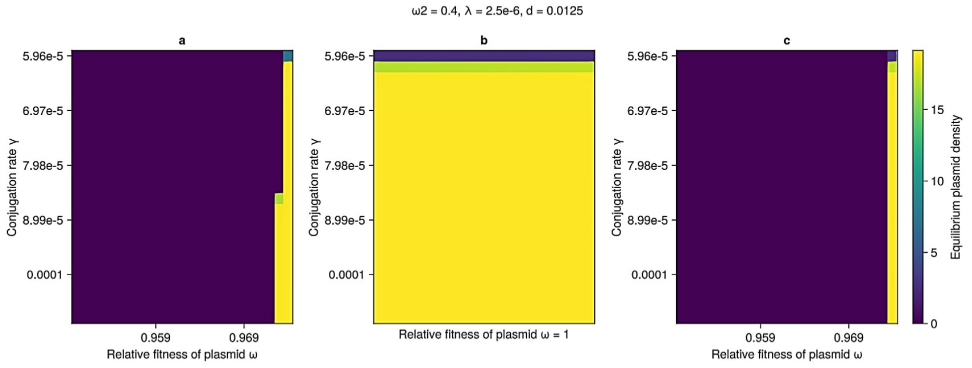
ω2 = 0.4, λ = 2.5e-6, d = 0.0125
a
b
c
Conjugation rate γ
5.96e-5
6.97e-5
7.98e-5
8.99e-5
0.0001
0.959
0.969
Relative fitness of plasmid ω
Relative fitness of plasmid ω = 1
Equilibrium plasmid density
0
5
10
15

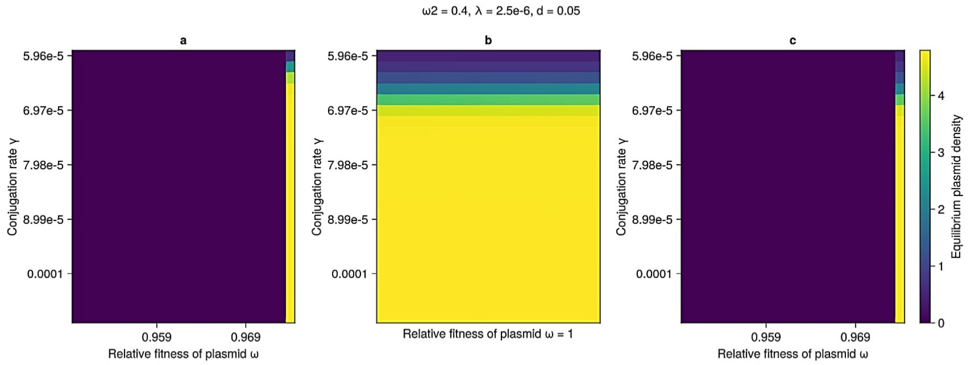
ω2 = 0.4, λ = 2.5e-6, d = 0.05
a
b
c
Conjugation rate γ
5.96e-5
6.97e-5
7.98e-5
8.99e-5
0.0001
0.959
0.969
Relative fitness of plasmid ω
Relative fitness of plasmid ω = 1
Equilibrium plasmid density
0
1
2
3
4

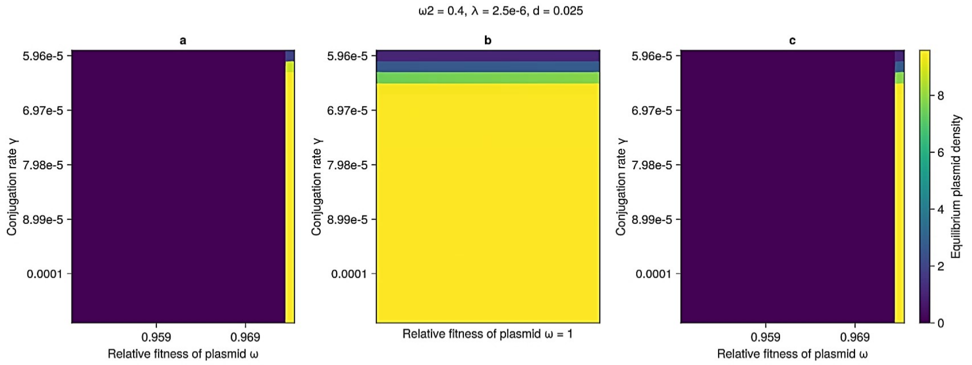
ω2 = 0.4, λ = 2.5e-6, d = 0.025
a
b
c
Conjugation rate γ
5.96e-5
6.97e-5
7.98e-5
8.99e-5
0.0001
0.959
0.969
Relative fitness of plasmid ω
Relative fitness of plasmid ω = 1
Equilibrium plasmid density
0
2
4
6
8

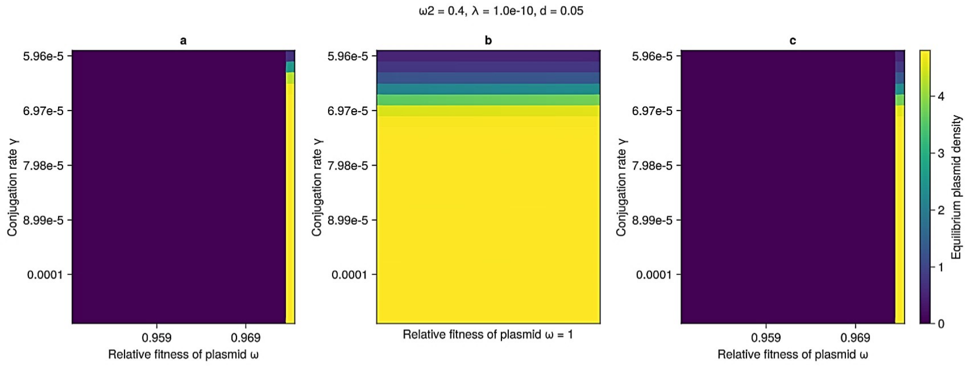
ω2 = 0.4, λ = 1.0e-10, d = 0.05
a
b
c
Conjugation rate γ
5.96e-5
6.97e-5
7.98e-5
8.99e-5
0.0001
0.959
0.969
Relative fitness of plasmid ω
Relative fitness of plasmid ω = 1
Equilibrium plasmid density
0
1
2
3
4

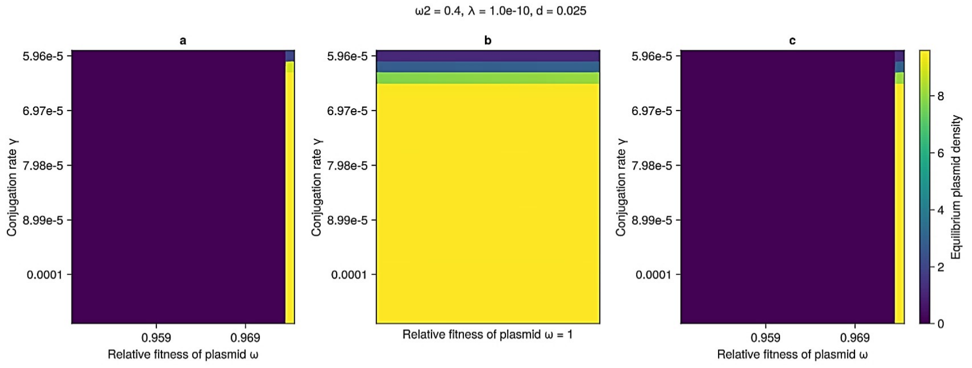
ω2 = 0.4, λ = 1.0e-10, d = 0.025
a
b
c
Conjugation rate γ
5.96e-5
6.97e-5
7.98e-5
8.99e-5
0.0001
0.959
0.969
Relative fitness of plasmid ω
Relative fitness of plasmid ω = 1
Equilibrium plasmid density
0
2
4
6
8

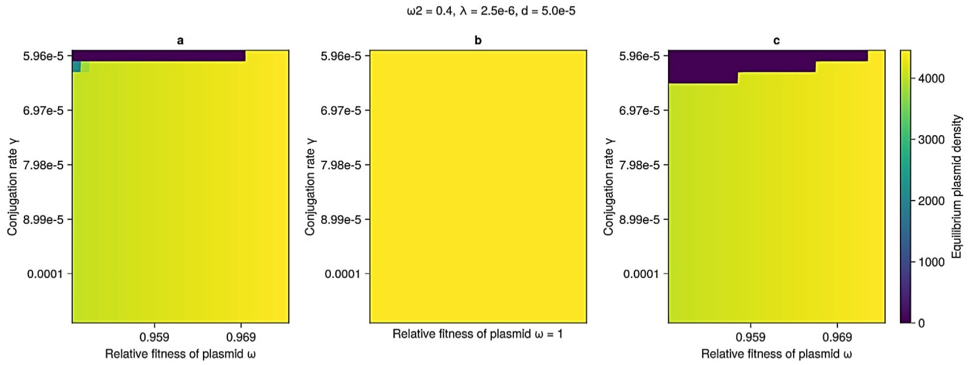
ω2 = 0.4, λ = 2.5e-6, d = 5.0e-5
a
b
c
Conjugation rate γ
5.96e-5
6.97e-5
7.98e-5
8.99e-5
0.0001
0.959
0.969
Relative fitness of plasmid ω
Relative fitness of plasmid ω = 1
Equilibrium plasmid density
0
1000
2000
3000
4000

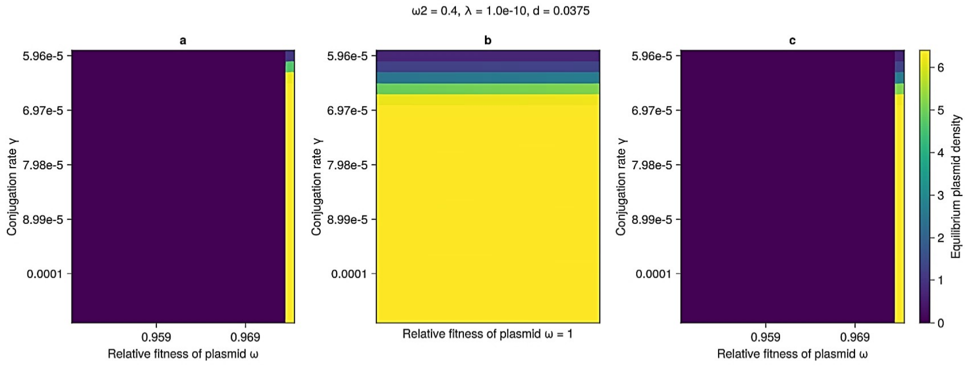
ω2 = 0.4, λ = 1.0e-10, d = 0.0375
a
b
c
Conjugation rate γ
5.96e-5
6.97e-5
7.98e-5
8.99e-5
0.0001
0.959
0.969
Relative fitness of plasmid ω
Relative fitness of plasmid ω = 1
Equilibrium plasmid density
0
1
2
3
4
5
6

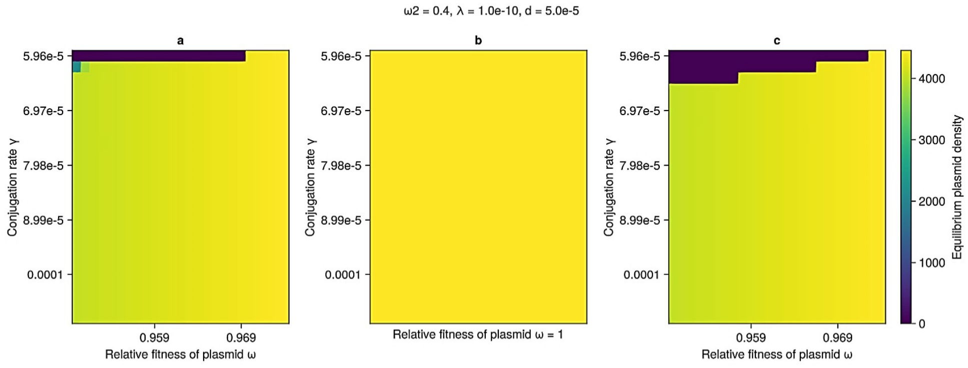
ω2 = 0.4, λ = 1.0e-10, d = 5.0e-5
a
b
c
Conjugation rate γ
5.96e-5
6.97e-5
7.98e-5
8.99e-5
0.0001
0.959
0.969
Relative fitness of plasmid ω
Relative fitness of plasmid ω = 1
Equilibrium plasmid density
0
1000
2000
3000
4000

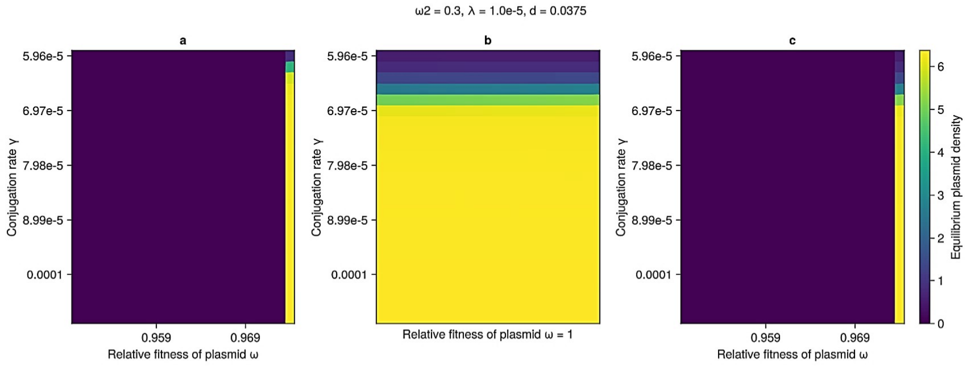
ω2 = 0.3, λ = 1.0e-5, d = 0.0375
a
b
c
Conjugation rate γ
5.96e-5
6.97e-5
7.98e-5
8.99e-5
0.0001
0.959
0.969
Relative fitness of plasmid ω
Relative fitness of plasmid ω = 1
Equilibrium plasmid density
0
1
2
3
4
5
6

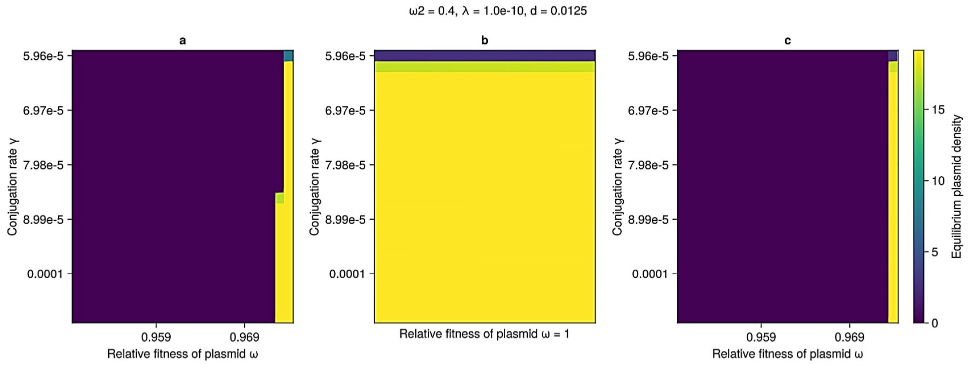
ω2 = 0.4, λ = 1.0e-10, d = 0.0125
a
b
c
Conjugation rate γ
5.96e-5
6.97e-5
7.98e-5
8.99e-5
0.0001
0.959
0.969
Relative fitness of plasmid ω
Relative fitness of plasmid ω = 1
Equilibrium plasmid density
0
5
10
15

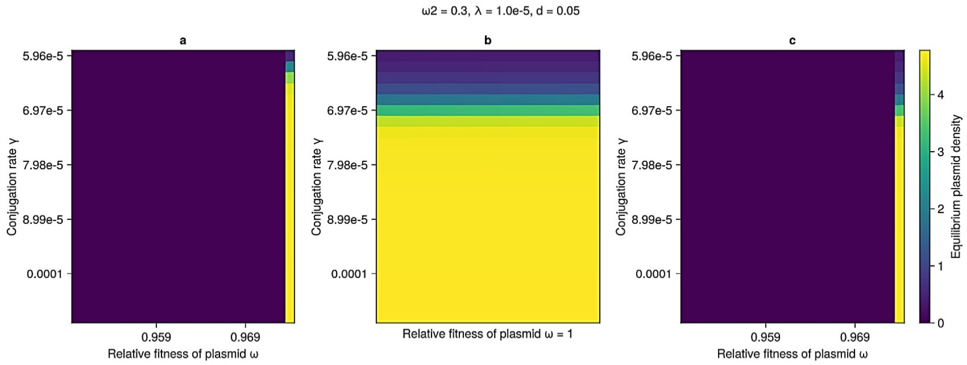
ω2 = 0.3, λ = 1.0e-5, d = 0.05
a
b
c
Conjugation rate γ
5.96e-5
6.97e-5
7.98e-5
8.99e-5
0.0001
0.959
0.969
Relative fitness of plasmid ω
Relative fitness of plasmid ω = 1
Equilibrium plasmid density
0
1
2
3
4

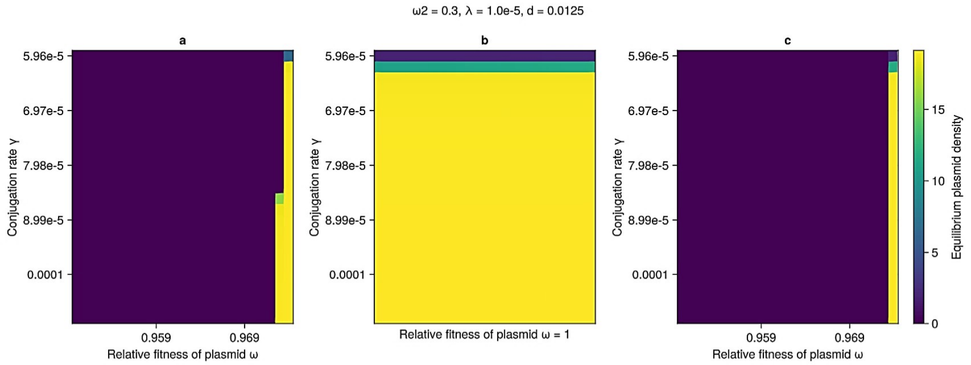
ω2 = 0.3, λ = 1.0e-5, d = 0.0125
a
b
c
Conjugation rate γ
5.96e-5
6.97e-5
7.98e-5
8.99e-5
0.0001
0.959
0.969
Relative fitness of plasmid ω
Relative fitness of plasmid ω = 1
Equilibrium plasmid density
0
5
10
15

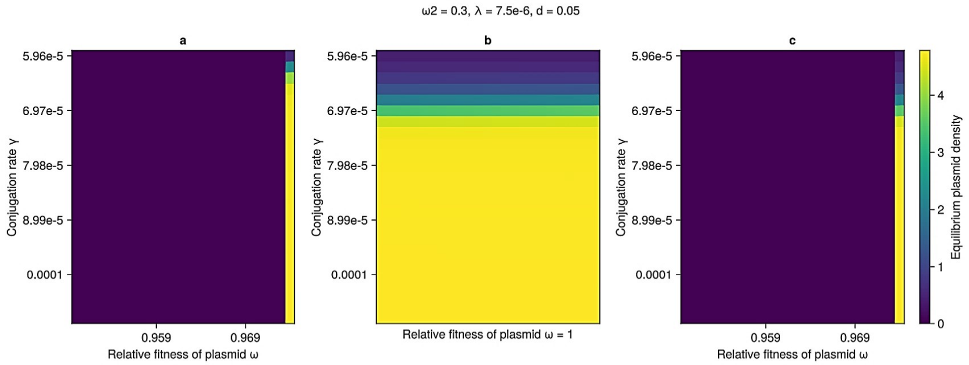
ω2 = 0.3, λ = 7.5e-6, d = 0.05
a
b
c
Conjugation rate γ
5.96e-5
6.97e-5
7.98e-5
8.99e-5
0.0001
0.959
0.969
Relative fitness of plasmid ω
Relative fitness of plasmid ω = 1
Equilibrium plasmid density
0
1
2
3
4

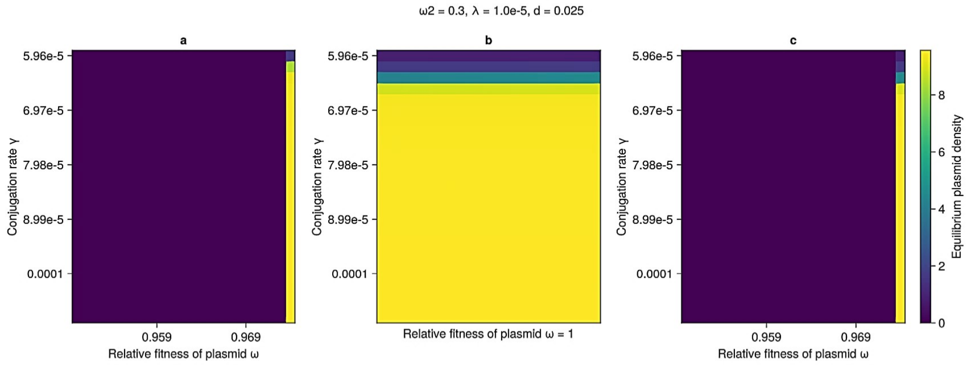
ω2 = 0.3, λ = 1.0e-5, d = 0.025
a
b
c
Conjugation rate γ
5.96e-5
6.97e-5
7.98e-5
8.99e-5
0.0001
0.959
0.969
Relative fitness of plasmid ω
Relative fitness of plasmid ω = 1
Equilibrium plasmid density
0
2
4
6
8

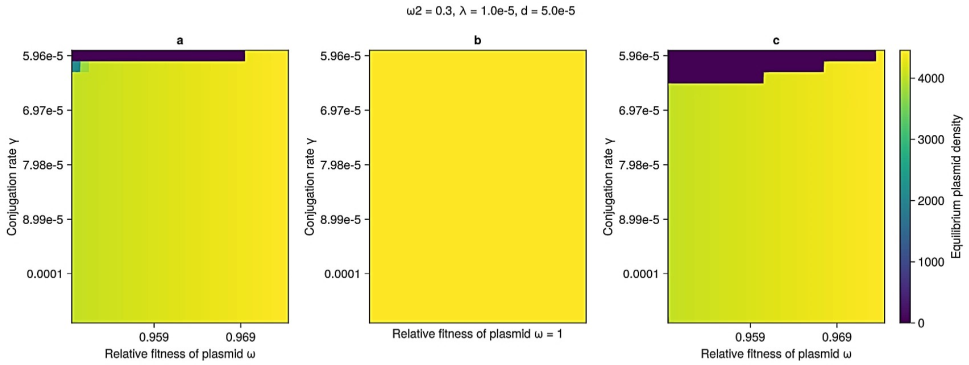
ω2 = 0.3, λ = 1.0e-5, d = 5.0e-5
a
b
c
Conjugation rate γ
5.96e-5
6.97e-5
7.98e-5
8.99e-5
0.0001
0.959
0.969
Relative fitness of plasmid ω
Relative fitness of plasmid ω = 1
Equilibrium plasmid density
0
1000
2000
3000
4000

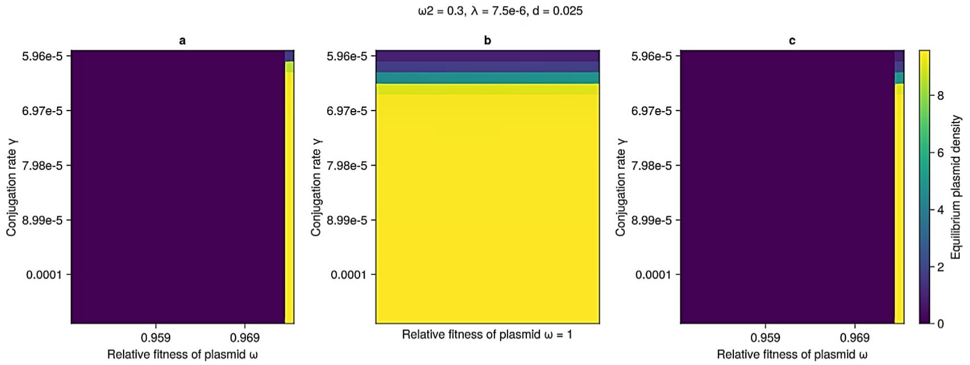
ω2 = 0.3, λ = 7.5e-6, d = 0.025
a
b
c
5.96e-5
6.97e-5
7.98e-5
8.99e-5
0.0001
Conjugation rate γ
0.959
0.969
Relative fitness of plasmid ω
Relative fitness of plasmid ω = 1
0
2
4
6
8
Equilibrium plasmid density

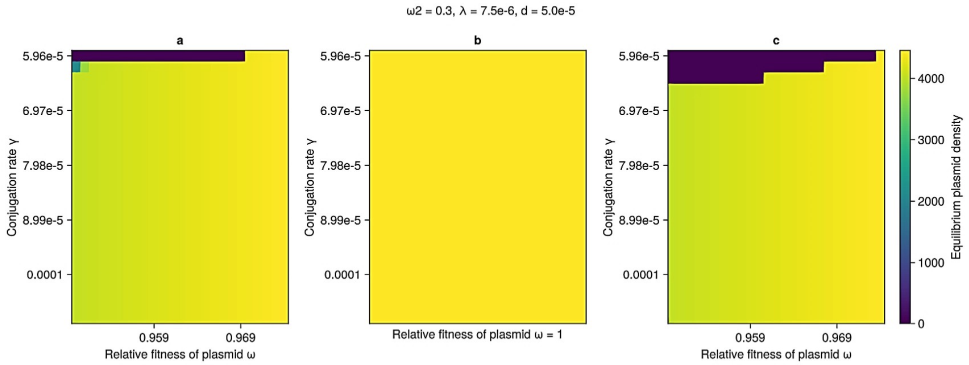
ω2 = 0.3, λ = 7.5e-6, d = 5.0e-5
a
b
c
5.96e-5
6.97e-5
7.98e-5
8.99e-5
0.0001
Conjugation rate γ
0.959
0.969
Relative fitness of plasmid ω
Relative fitness of plasmid ω = 1
0
1000
2000
3000
4000
Equilibrium plasmid density

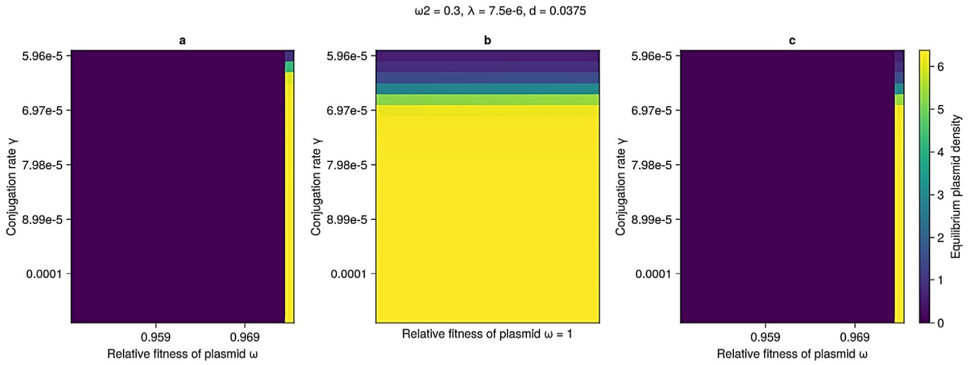
ω2 = 0.3, λ = 7.5e-6, d = 0.0375
a
b
c
5.96e-5
6.97e-5
7.98e-5
8.99e-5
0.0001
Conjugation rate γ
0.959
0.969
Relative fitness of plasmid ω
Relative fitness of plasmid ω = 1
0
1
2
3
4
5
6
Equilibrium plasmid density

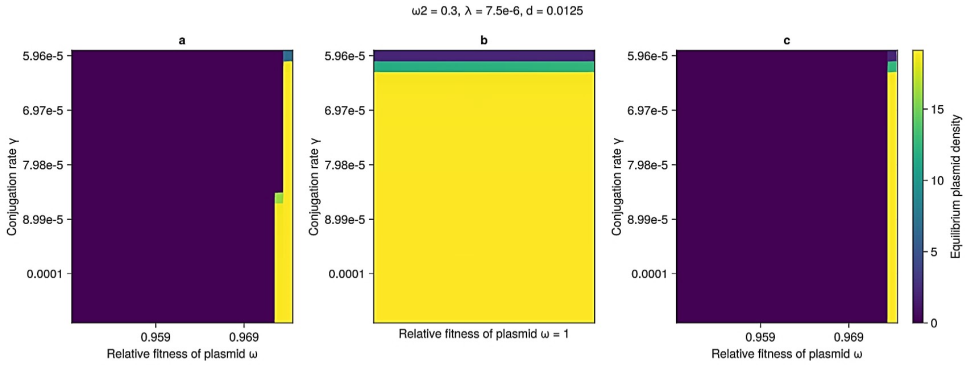
ω2 = 0.3, λ = 7.5e-6, d = 0.0125
a
b
c
5.96e-5
6.97e-5
7.98e-5
8.99e-5
0.0001
Conjugation rate γ
0.959
0.969
Relative fitness of plasmid ω
Relative fitness of plasmid ω = 1
0
5
10
15
Equilibrium plasmid density

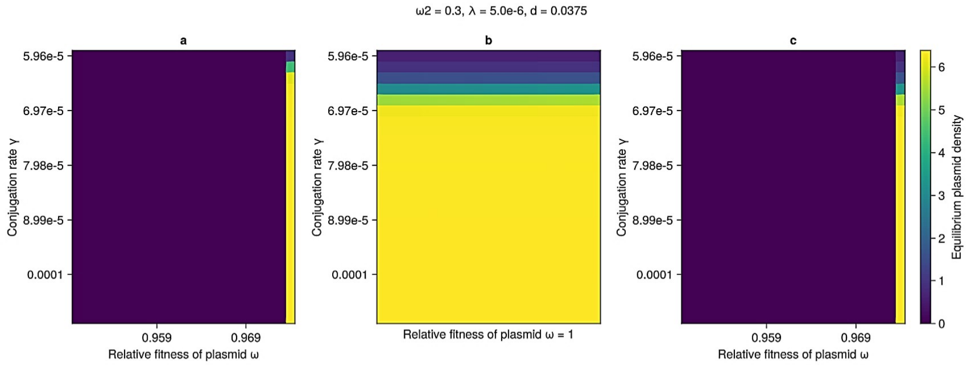
ω2 = 0.3, λ = 5.0e-6, d = 0.0375
a
b
c
Conjugation rate γ
5.96e-5
6.97e-5
7.98e-5
8.99e-5
0.0001
0.959
0.969
Relative fitness of plasmid ω
Relative fitness of plasmid ω = 1
Equilibrium plasmid density
0
1
2
3
4
5
6

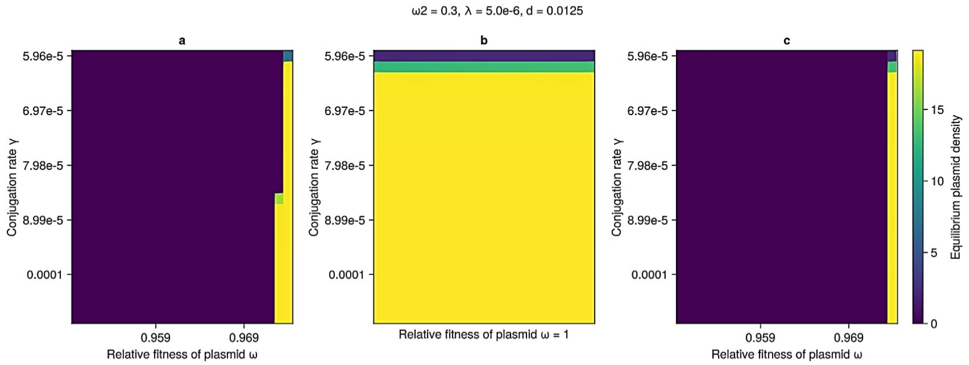
ω2 = 0.3, λ = 5.0e-6, d = 0.0125
a
b
c
Conjugation rate γ
5.96e-5
6.97e-5
7.98e-5
8.99e-5
0.0001
0.959
0.969
Relative fitness of plasmid ω
Relative fitness of plasmid ω = 1
Equilibrium plasmid density
0
5
10
15

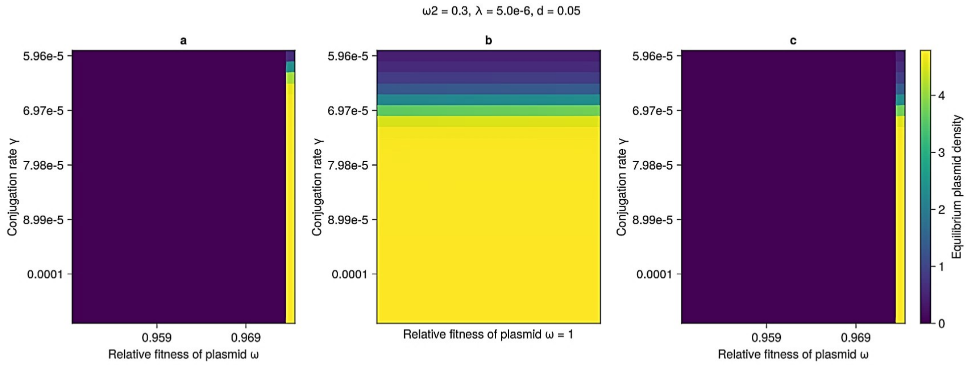
ω2 = 0.3, λ = 5.0e-6, d = 0.05
a
b
c
Conjugation rate γ
5.96e-5
6.97e-5
7.98e-5
8.99e-5
0.0001
0.959
0.969
Relative fitness of plasmid ω
Relative fitness of plasmid ω = 1
Equilibrium plasmid density
0
1
2
3
4

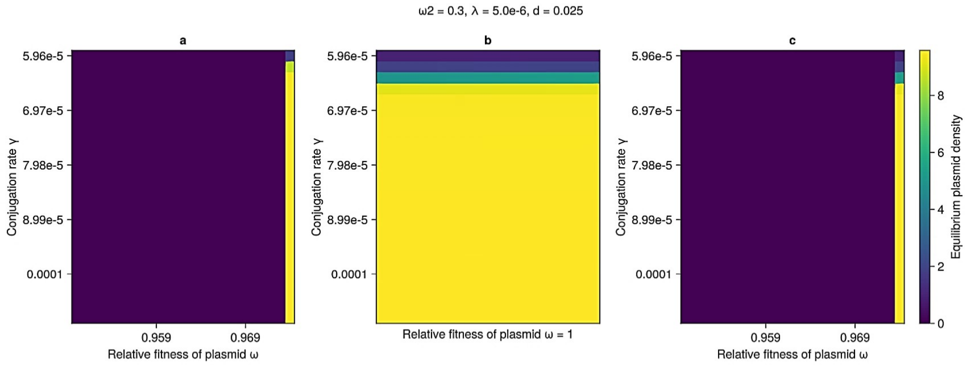
ω2 = 0.3, λ = 5.0e-6, d = 0.025
a
b
c
Conjugation rate γ
5.96e-5
6.97e-5
7.98e-5
8.99e-5
0.0001
0.959
0.969
Relative fitness of plasmid ω
Relative fitness of plasmid ω = 1
Equilibrium plasmid density
0
2
4
6
8

ω2 = 0.3, λ = 2.5e-6, d = 0.05

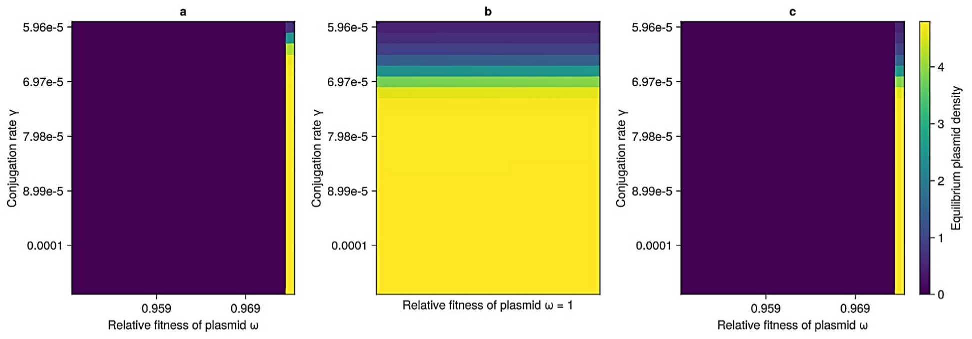


ω2 = 0.3, λ = 2.5e-6, d = 0.025

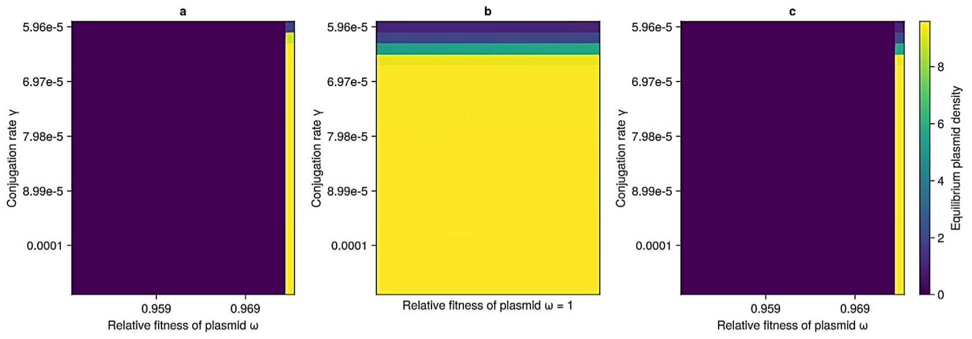


ω2 = 0.3, λ = 5.0e-6, d = 5.0e-5

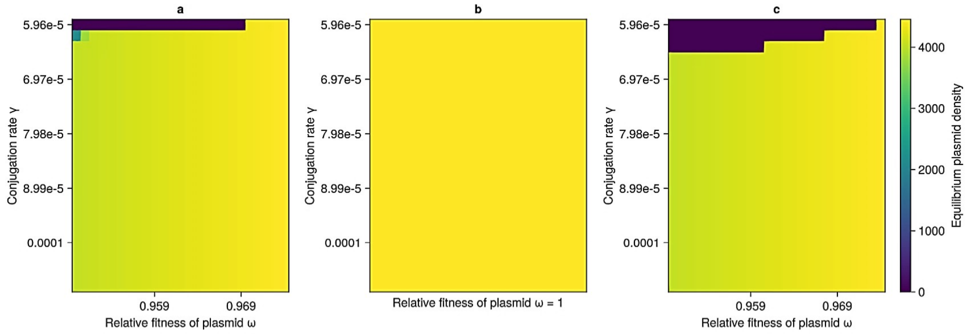


ω2 = 0.3, λ = 2.5e-6, d = 0.0375

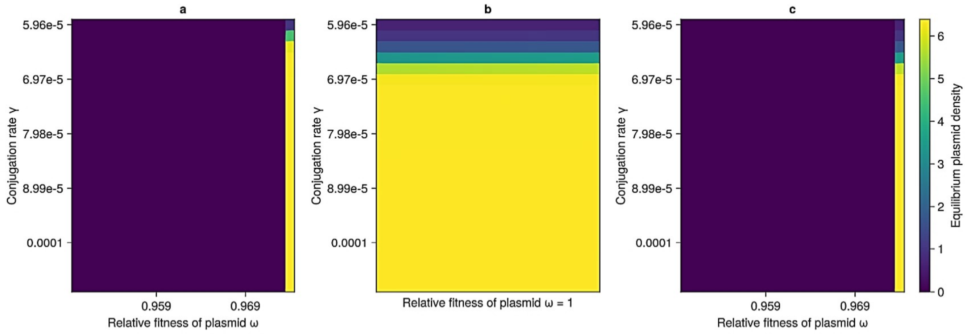

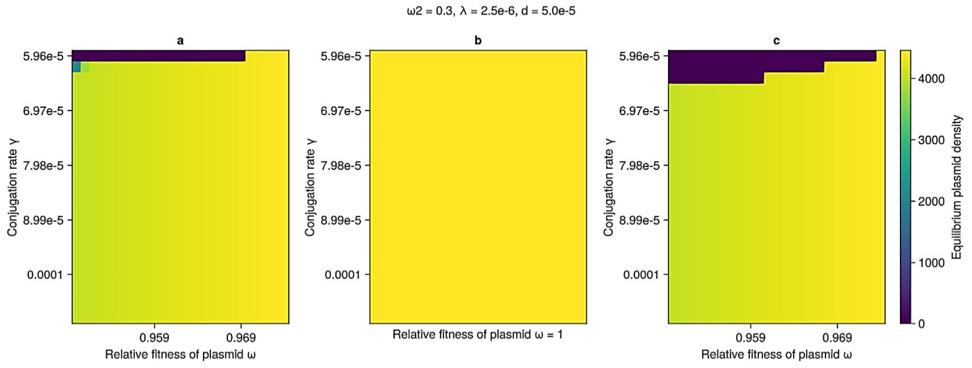
ω2 = 0.3, λ = 2.5e-6, d = 5.0e-5
a
b
c
Conjugation rate γ
5.96e-5
6.97e-5
7.98e-5
8.99e-5
0.0001
0.959
0.969
Relative fitness of plasmid ω
Relative fitness of plasmid ω = 1
Equilibrium plasmid density
0
1000
2000
3000
4000

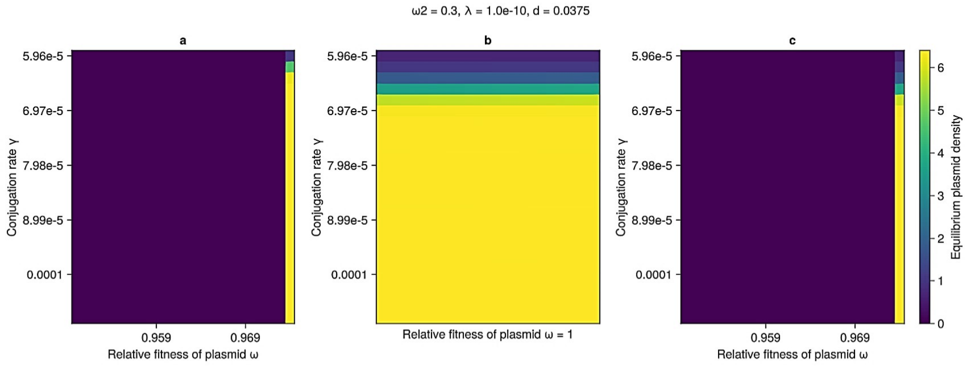
ω2 = 0.3, λ = 1.0e-10, d = 0.0375
a
b
c
Conjugation rate γ
5.96e-5
6.97e-5
7.98e-5
8.99e-5
0.0001
0.959
0.969
Relative fitness of plasmid ω
Relative fitness of plasmid ω = 1
Equilibrium plasmid density
0
1
2
3
4
5
6

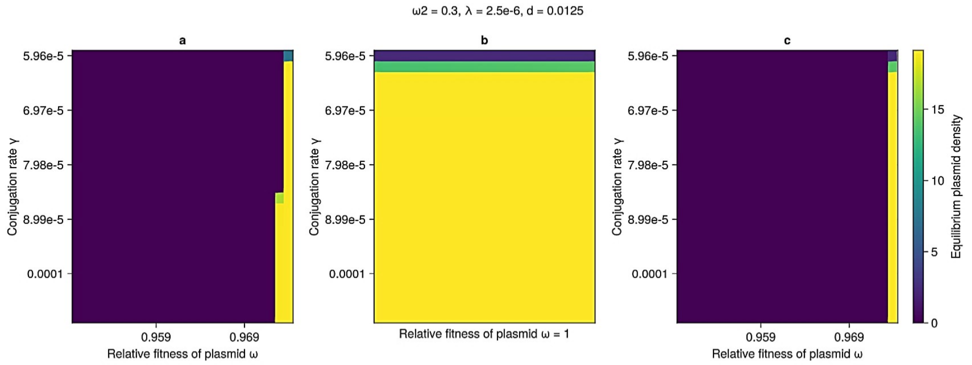
ω2 = 0.3, λ = 2.5e-6, d = 0.0125
a
b
c
Conjugation rate γ
5.96e-5
6.97e-5
7.98e-5
8.99e-5
0.0001
0.959
0.969
Relative fitness of plasmid ω
Relative fitness of plasmid ω = 1
Equilibrium plasmid density
0
5
10
15

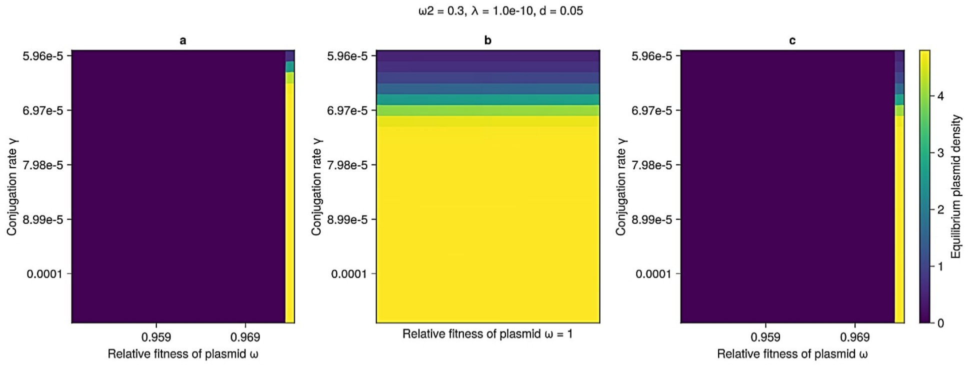
ω2 = 0.3, λ = 1.0e-10, d = 0.05
a
b
c
Conjugation rate γ
5.96e-5
6.97e-5
7.98e-5
8.99e-5
0.0001
0.959
0.969
Relative fitness of plasmid ω
Relative fitness of plasmid ω = 1
Equilibrium plasmid density
0
1
2
3
4

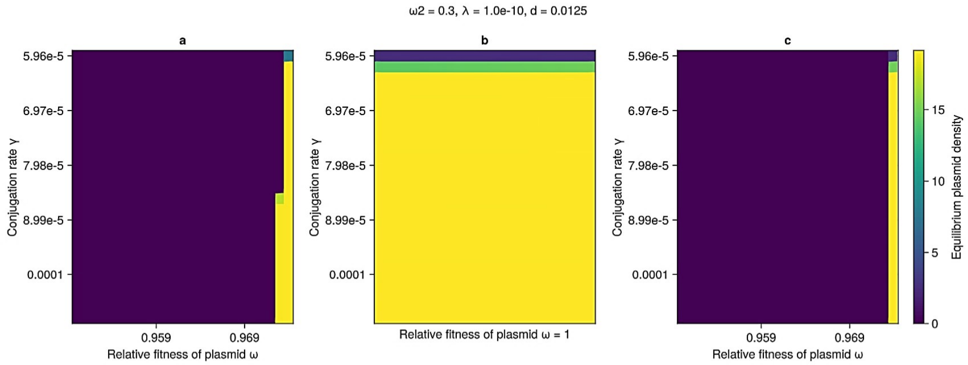
ω2 = 0.3, λ = 1.0e-10, d = 0.0125
a
b
c
Conjugation rate γ
5.96e-5
6.97e-5
7.98e-5
8.99e-5
0.0001
0.959
0.969
Relative fitness of plasmid ω
Relative fitness of plasmid ω = 1
Equilibrium plasmid density
0
5
10
15

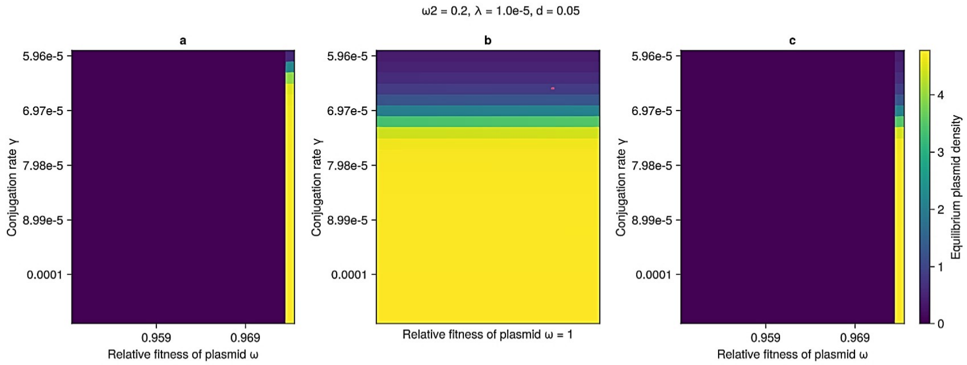
ω2 = 0.2, λ = 1.0e-5, d = 0.05
a
b
c
Conjugation rate γ
5.96e-5
6.97e-5
7.98e-5
8.99e-5
0.0001
0.959
0.969
Relative fitness of plasmid ω
Relative fitness of plasmid ω = 1
Equilibrium plasmid density
0
1
2
3
4

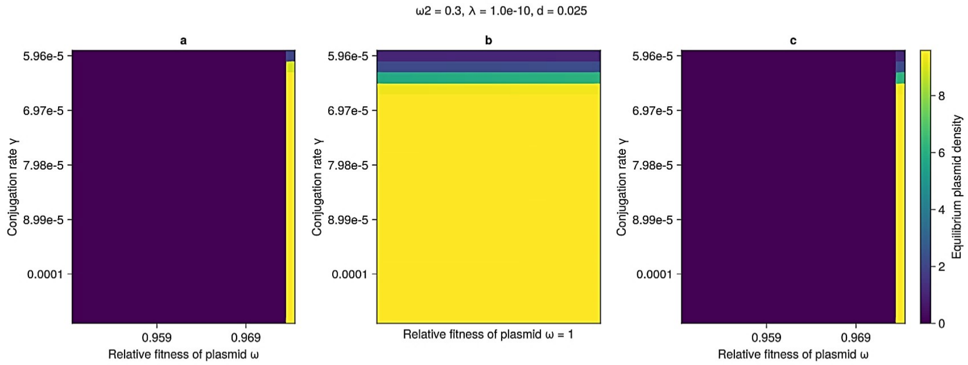
ω2 = 0.3, λ = 1.0e-10, d = 0.025
a
b
c
Conjugation rate γ
5.96e-5
6.97e-5
7.98e-5
8.99e-5
0.0001
0.959
0.969
Relative fitness of plasmid ω
Relative fitness of plasmid ω = 1
Equilibrium plasmid density
0
2
4
6
8

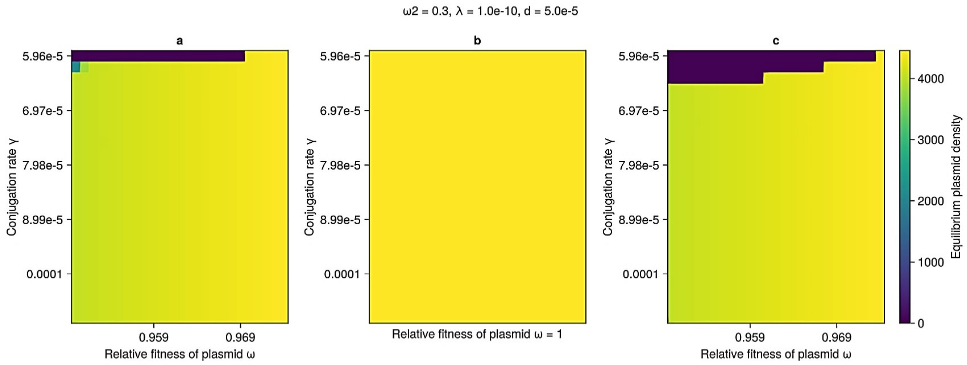
ω2 = 0.3, λ = 1.0e-10, d = 5.0e-5
a
b
c
Conjugation rate γ
5.96e-5
6.97e-5
7.98e-5
8.99e-5
0.0001
0.959
0.969
Relative fitness of plasmid ω
Relative fitness of plasmid ω = 1
Equilibrium plasmid density
0
1000
2000
3000
4000

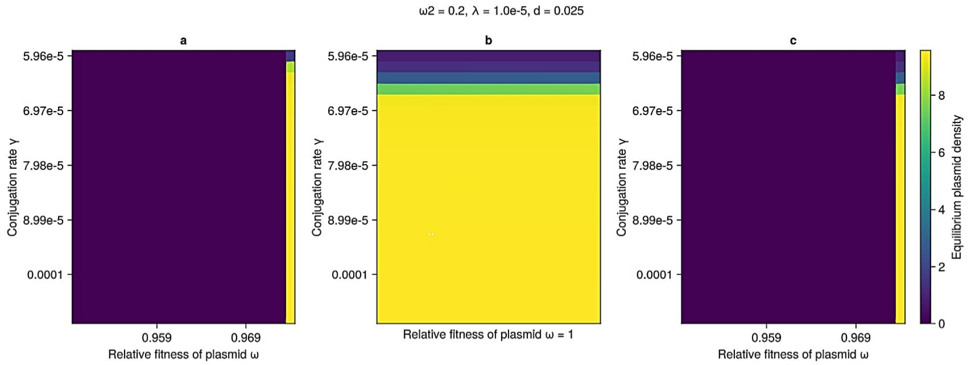
ω2 = 0.2, λ = 1.0e-5, d = 0.025
a
b
c
Conjugation rate γ
5.96e-5
6.97e-5
7.98e-5
8.99e-5
0.0001
0.959
0.969
Relative fitness of plasmid ω
Relative fitness of plasmid ω = 1
Equilibrium plasmid density
0
2
4
6
8

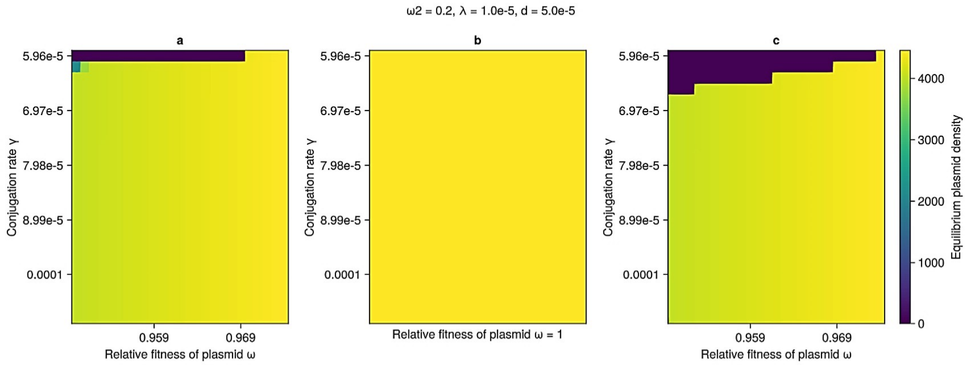
ω2 = 0.2, λ = 1.0e-5, d = 5.0e-5
a
b
c
Conjugation rate γ
5.96e-5
6.97e-5
7.98e-5
8.99e-5
0.0001
0.959
0.969
Relative fitness of plasmid ω
Relative fitness of plasmid ω = 1
Equilibrium plasmid density
0
1000
2000
3000
4000

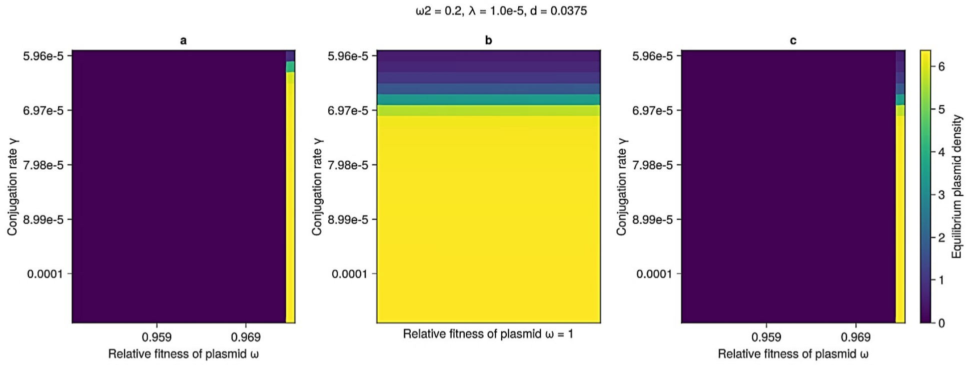
ω2 = 0.2, λ = 1.0e-5, d = 0.0375
a
b
c
Conjugation rate γ
5.96e-5
6.97e-5
7.98e-5
8.99e-5
0.0001
0.959
0.969
Relative fitness of plasmid ω
Relative fitness of plasmid ω = 1
Equilibrium plasmid density
0
1
2
3
4
5
6

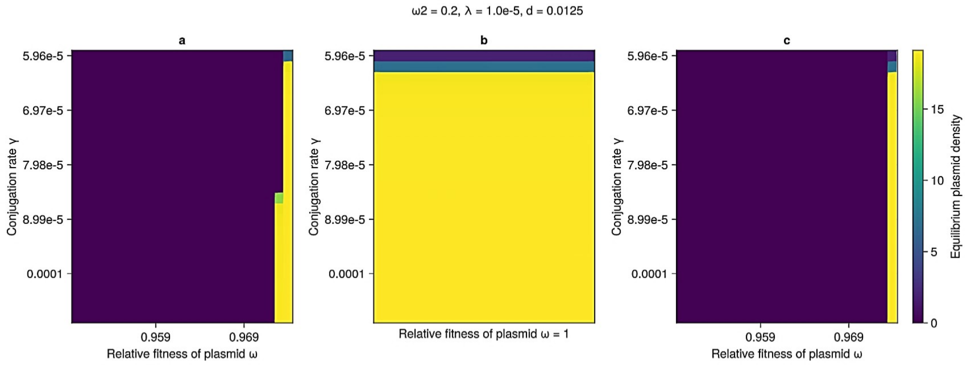
ω2 = 0.2, λ = 1.0e-5, d = 0.0125
a
b
c
Conjugation rate γ
5.96e-5
6.97e-5
7.98e-5
8.99e-5
0.0001
0.959
0.969
Relative fitness of plasmid ω
Relative fitness of plasmid ω = 1
Equilibrium plasmid density
0
5
10
15

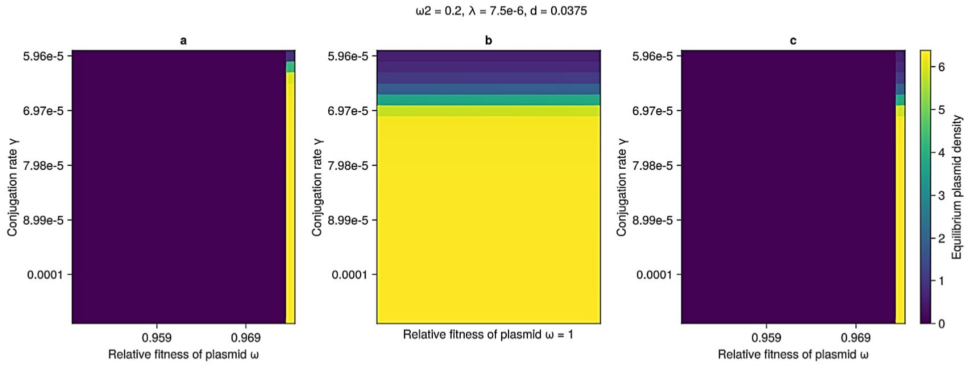
ω2 = 0.2, λ = 7.5e-6, d = 0.0375
a
b
c
Conjugation rate γ
5.96e-5
6.97e-5
7.98e-5
8.99e-5
0.0001
0.959
0.969
Relative fitness of plasmid ω
Relative fitness of plasmid ω = 1
Equilibrium plasmid density
0
1
2
3
4
5
6

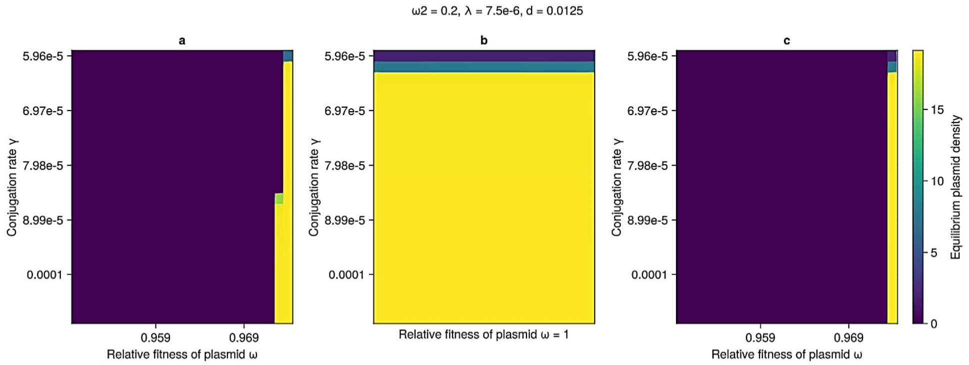
ω2 = 0.2, λ = 7.5e-6, d = 0.0125
a
b
c
Conjugation rate γ
5.96e-5
6.97e-5
7.98e-5
8.99e-5
0.0001
0.959
0.969
Relative fitness of plasmid ω
Relative fitness of plasmid ω = 1
Equilibrium plasmid density
0
5
10
15

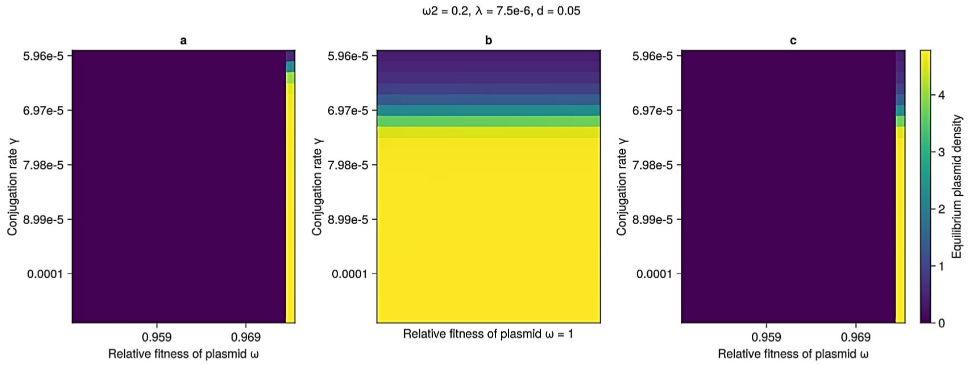
ω2 = 0.2, λ = 7.5e-6, d = 0.05
a
b
c
Conjugation rate γ
5.96e-5
6.97e-5
7.98e-5
8.99e-5
0.0001
0.959
0.969
Relative fitness of plasmid ω
Relative fitness of plasmid ω = 1
Equilibrium plasmid density
0
1
2
3
4

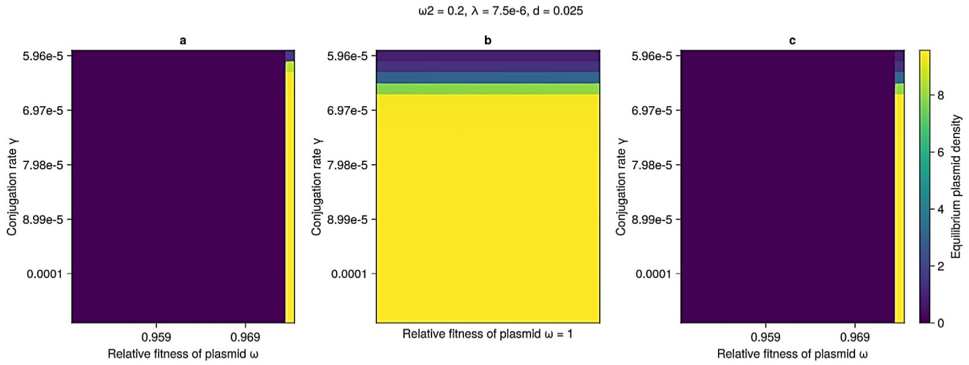
ω2 = 0.2, λ = 7.5e-6, d = 0.025
a
b
c
Conjugation rate γ
5.96e-5
6.97e-5
7.98e-5
8.99e-5
0.0001
0.959
0.969
Relative fitness of plasmid ω
Relative fitness of plasmid ω = 1
Equilibrium plasmid density
0
2
4
6
8

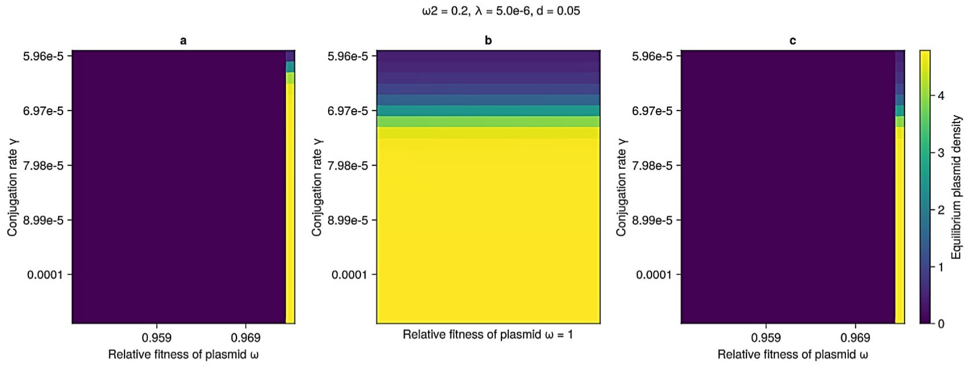
ω2 = 0.2, λ = 5.0e-6, d = 0.05
a
b
c
Conjugation rate γ
5.96e-5
6.97e-5
7.98e-5
8.99e-5
0.0001
0.959
0.969
Relative fitness of plasmid ω
Relative fitness of plasmid ω = 1
Equilibrium plasmid density

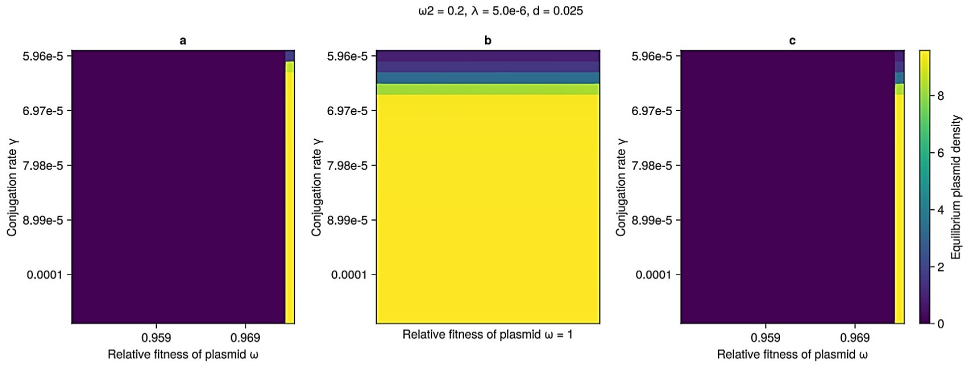
ω2 = 0.2, λ = 5.0e-6, d = 0.025
a
b
c
Conjugation rate γ
5.96e-5
6.97e-5
7.98e-5
8.99e-5
0.0001
0.959
0.969
Relative fitness of plasmid ω
Relative fitness of plasmid ω = 1
Equilibrium plasmid density

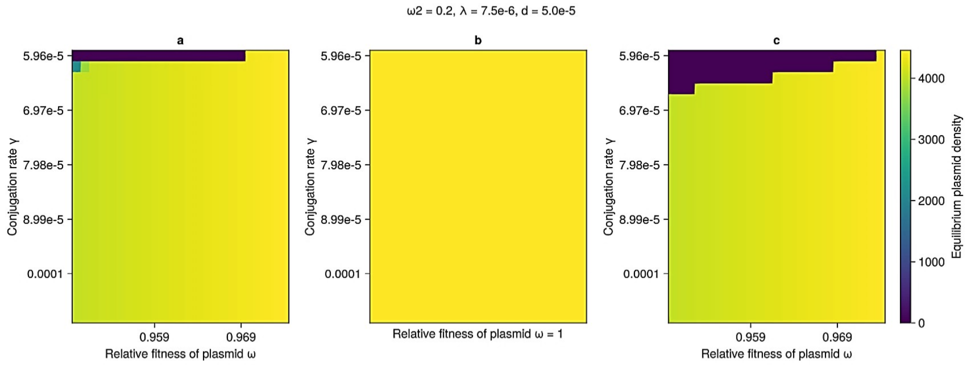
ω2 = 0.2, λ = 7.5e-6, d = 5.0e-5
a
b
c
Conjugation rate γ
5.96e-5
6.97e-5
7.98e-5
8.99e-5
0.0001
0.959
0.969
Relative fitness of plasmid ω
Relative fitness of plasmid ω = 1
Equilibrium plasmid density

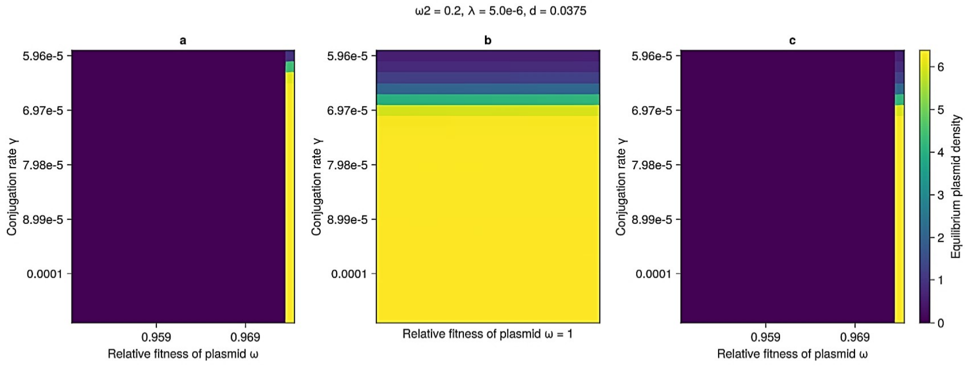
ω2 = 0.2, λ = 5.0e-6, d = 0.0375
a
b
c
Conjugation rate γ
5.96e-5
6.97e-5
7.98e-5
8.99e-5
0.0001
0.959
0.969
Relative fitness of plasmid ω
Relative fitness of plasmid ω = 1
Equilibrium plasmid density

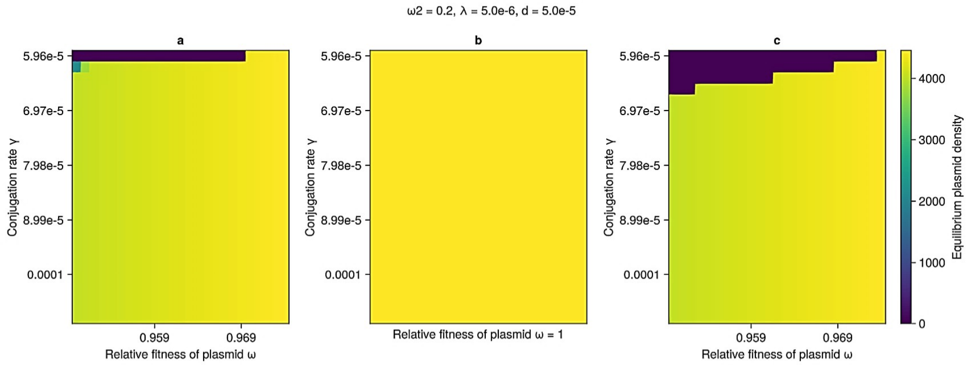
ω2 = 0.2, λ = 5.0e-6, d = 5.0e-5
a
b
c
Conjugation rate γ
5.96e-5
6.97e-5
7.98e-5
8.99e-5
0.0001
0.959
0.969
Relative fitness of plasmid ω
Relative fitness of plasmid ω = 1
Equilibrium plasmid density
0
1000
2000
3000
4000

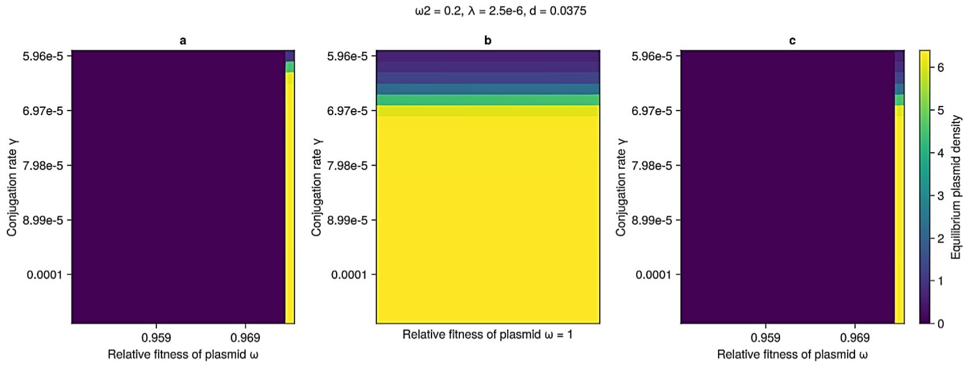
ω2 = 0.2, λ = 2.5e-6, d = 0.0375
a
b
c
Conjugation rate γ
5.96e-5
6.97e-5
7.98e-5
8.99e-5
0.0001
0.959
0.969
Relative fitness of plasmid ω
Relative fitness of plasmid ω = 1
Equilibrium plasmid density
0
1
2
3
4
5
6

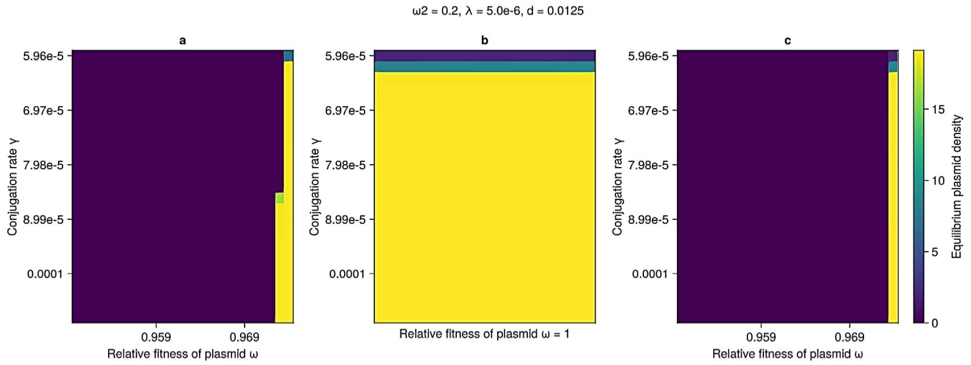
ω2 = 0.2, λ = 5.0e-6, d = 0.0125
a
b
c
Conjugation rate γ
5.96e-5
6.97e-5
7.98e-5
8.99e-5
0.0001
0.959
0.969
Relative fitness of plasmid ω
Relative fitness of plasmid ω = 1
Equilibrium plasmid density
0
5
10
15

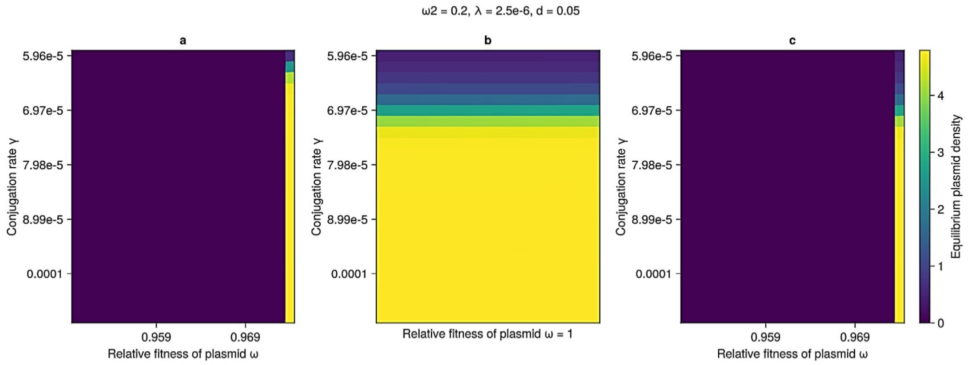
ω2 = 0.2, λ = 2.5e-6, d = 0.05
a
b
c
Conjugation rate γ
5.96e-5
6.97e-5
7.98e-5
8.99e-5
0.0001
0.959
0.969
Relative fitness of plasmid ω
Relative fitness of plasmid ω = 1
Equilibrium plasmid density
0
1
2
3
4

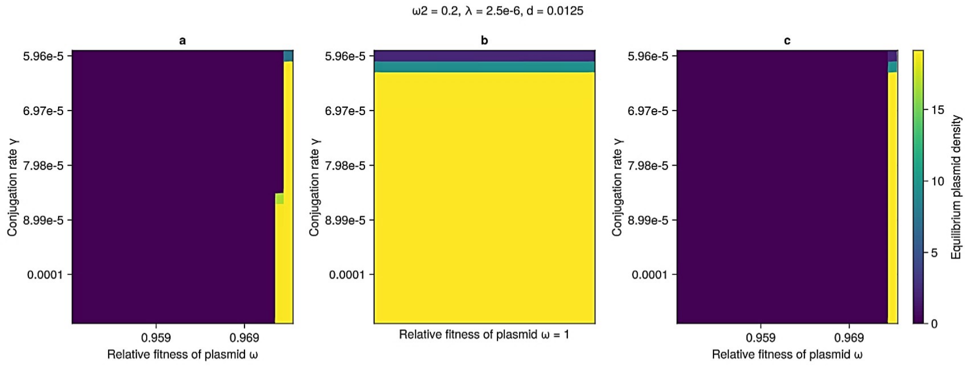
ω2 = 0.2, λ = 2.5e-6, d = 0.0125
a
b
c
Conjugation rate γ
5.96e-5
6.97e-5
7.98e-5
8.99e-5
0.0001
0.959
0.969
Relative fitness of plasmid ω
Relative fitness of plasmid ω = 1
Equilibrium plasmid density
0
5
10
15

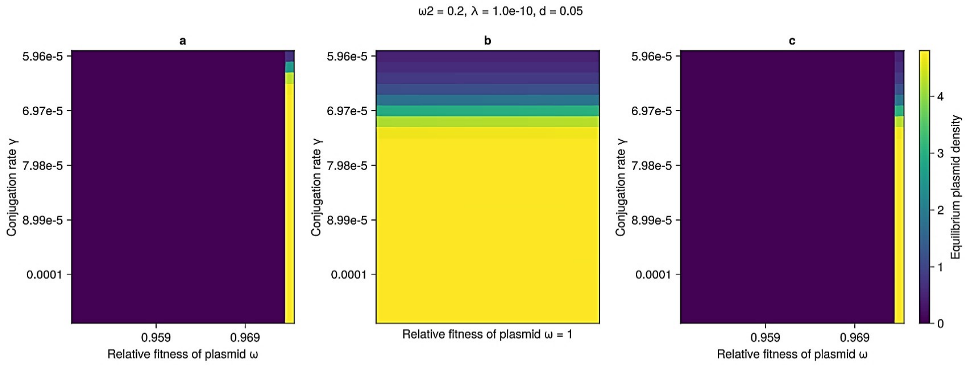
ω2 = 0.2, λ = 1.0e-10, d = 0.05
a
b
c
Conjugation rate γ
5.96e-5
6.97e-5
7.98e-5
8.99e-5
0.0001
0.959
0.969
Relative fitness of plasmid ω
Relative fitness of plasmid ω = 1
Equilibrium plasmid density
0
1
2
3
4

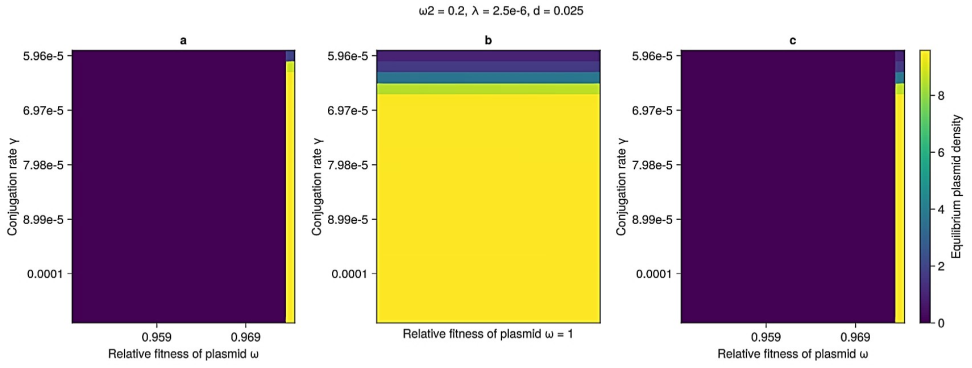
ω2 = 0.2, λ = 2.5e-6, d = 0.025
a
b
c
Conjugation rate γ
5.96e-5
6.97e-5
7.98e-5
8.99e-5
0.0001
0.959
0.969
Relative fitness of plasmid ω
Relative fitness of plasmid ω = 1
Equilibrium plasmid density
0
2
4
6
8

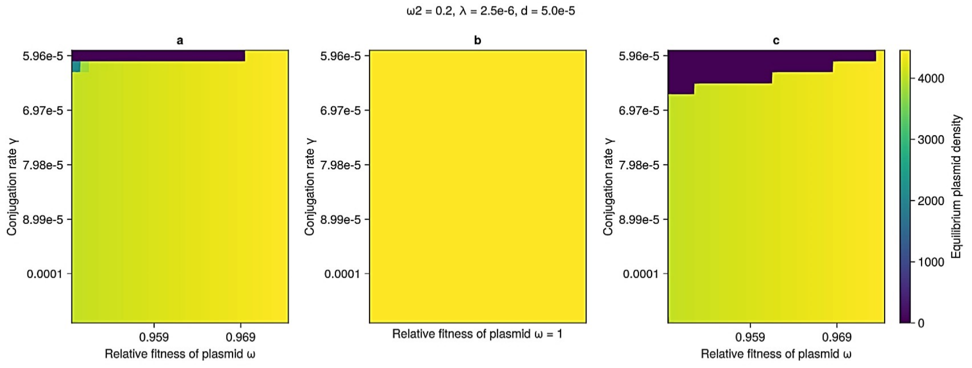
ω2 = 0.2, λ = 2.5e-6, d = 5.0e-5
a
b
c
Conjugation rate γ
5.96e-5
6.97e-5
7.98e-5
8.99e-5
0.0001
0.959
0.969
Relative fitness of plasmid ω
Relative fitness of plasmid ω = 1
Equilibrium plasmid density
0
1000
2000
3000
4000

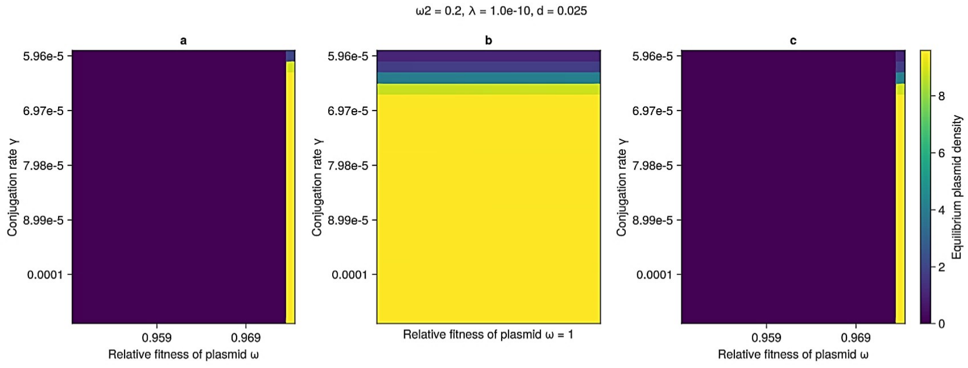
ω2 = 0.2, λ = 1.0e-10, d = 0.025
a
b
c
5.96e-5
6.97e-5
7.98e-5
8.99e-5
0.0001
Conjugation rate γ
0.959
0.969
Relative fitness of plasmid ω
Relative fitness of plasmid ω = 1
Equilibrium plasmid density
0
2
4
6
8

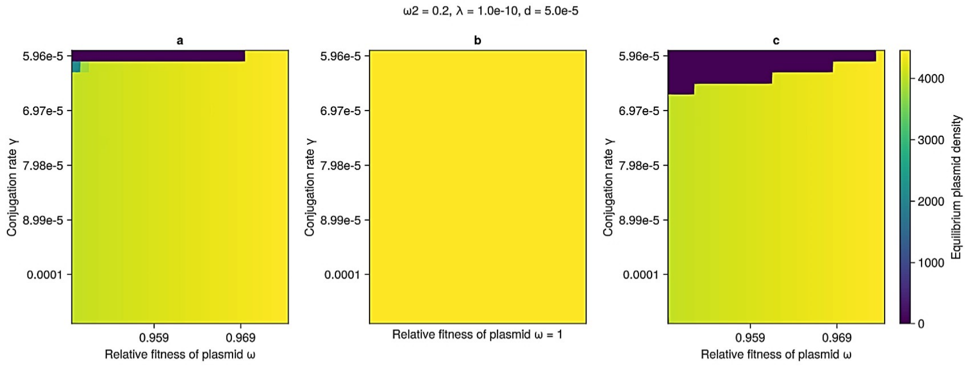
ω2 = 0.2, λ = 1.0e-10, d = 5.0e-5
a
b
c
5.96e-5
6.97e-5
7.98e-5
8.99e-5
0.0001
Conjugation rate γ
0.959
0.969
Relative fitness of plasmid ω
Relative fitness of plasmid ω = 1
Equilibrium plasmid density
0
1000
2000
3000
4000

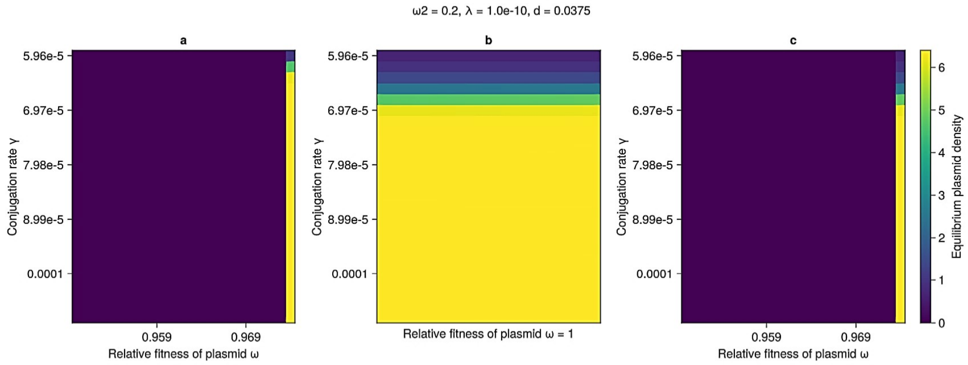
ω2 = 0.2, λ = 1.0e-10, d = 0.0375
a
b
c
5.96e-5
6.97e-5
7.98e-5
8.99e-5
0.0001
Conjugation rate γ
0.959
0.969
Relative fitness of plasmid ω
Relative fitness of plasmid ω = 1
Equilibrium plasmid density
0
1
2
3
4
5
6

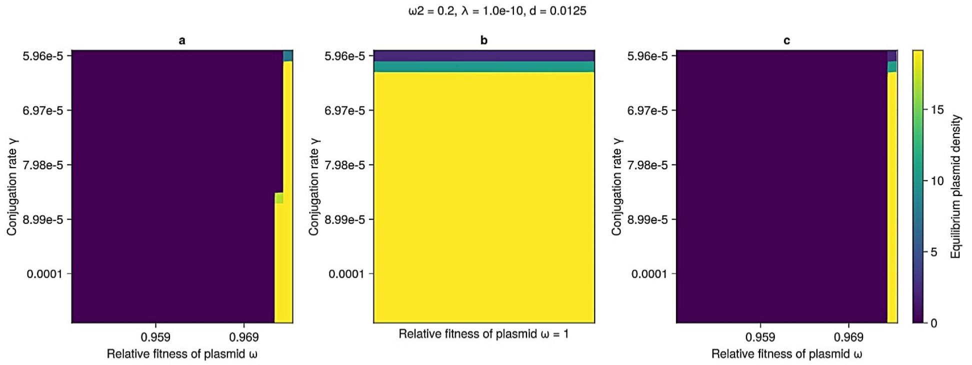
ω2 = 0.2, λ = 1.0e-10, d = 0.0125
a
b
c
5.96e-5
6.97e-5
7.98e-5
8.99e-5
0.0001
Conjugation rate γ
0.959
0.969
Relative fitness of plasmid ω
Relative fitness of plasmid ω = 1
Equilibrium plasmid density
0
5
10
15

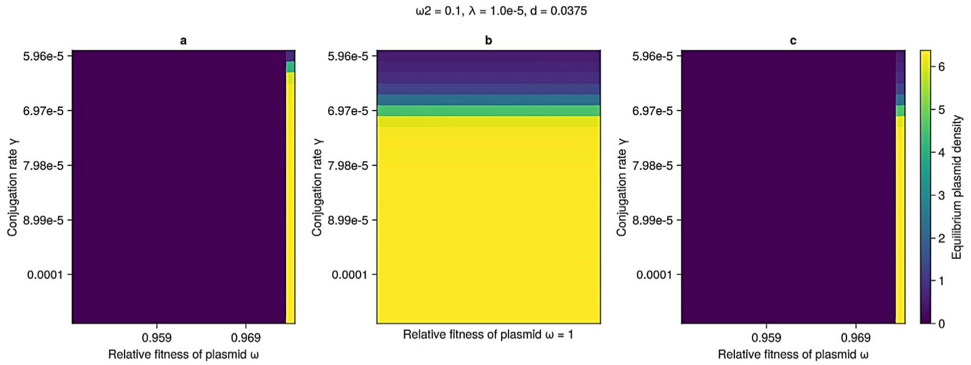
ω2 = 0.1, λ = 1.0e-5, d = 0.0375
a
b
c
Conjugation rate γ
5.96e-5
6.97e-5
7.98e-5
8.99e-5
0.0001
0.959
0.969
Relative fitness of plasmid ω
Relative fitness of plasmid ω = 1
Equilibrium plasmid density
0
1
2
3
4
5
6

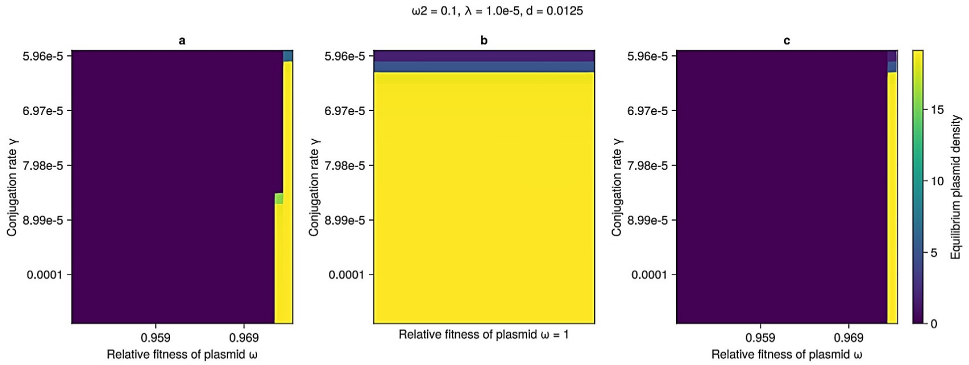
ω2 = 0.1, λ = 1.0e-5, d = 0.0125
a
b
c
Conjugation rate γ
5.96e-5
6.97e-5
7.98e-5
8.99e-5
0.0001
0.959
0.969
Relative fitness of plasmid ω
Relative fitness of plasmid ω = 1
Equilibrium plasmid density
0
5
10
15

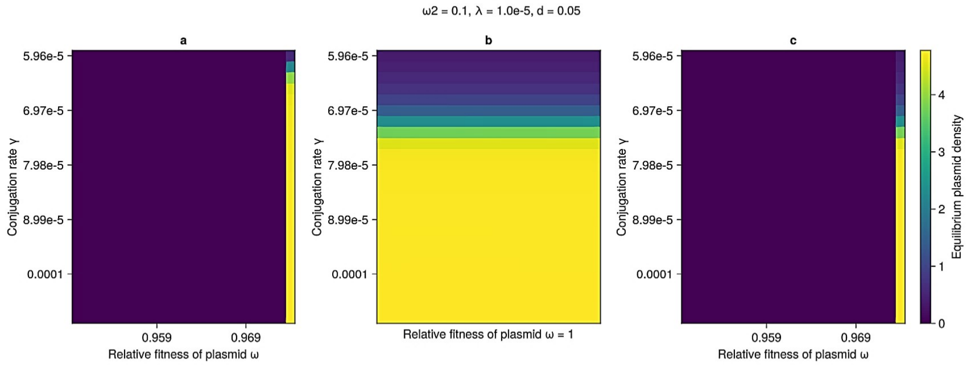
ω2 = 0.1, λ = 1.0e-5, d = 0.05
a
b
c
Conjugation rate γ
5.96e-5
6.97e-5
7.98e-5
8.99e-5
0.0001
0.959
0.969
Relative fitness of plasmid ω
Relative fitness of plasmid ω = 1
Equilibrium plasmid density
0
1
2
3
4

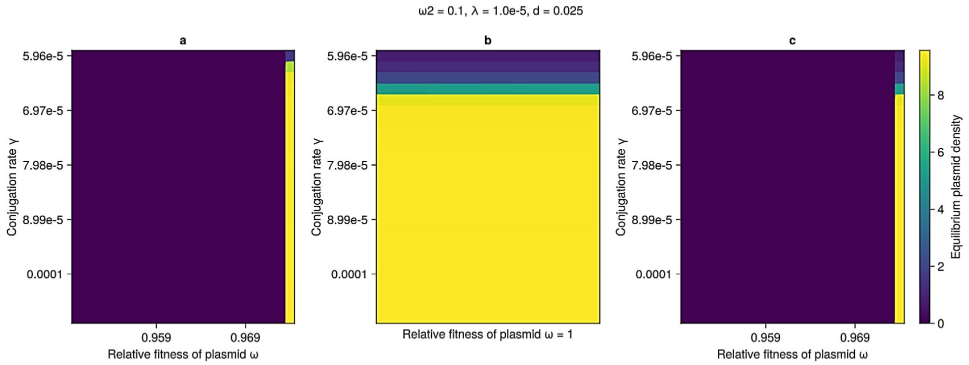
ω2 = 0.1, λ = 1.0e-5, d = 0.025
a
b
c
Conjugation rate γ
5.96e-5
6.97e-5
7.98e-5
8.99e-5
0.0001
0.959
0.969
Relative fitness of plasmid ω
Relative fitness of plasmid ω = 1
Equilibrium plasmid density
0
2
4
6
8

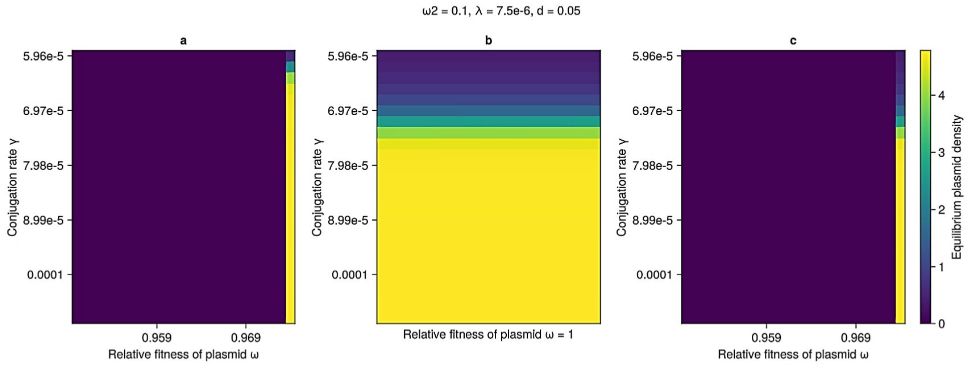
ω2 = 0.1, λ = 7.5e-6, d = 0.05
a
b
c
Conjugation rate γ
5.96e-5
6.97e-5
7.98e-5
8.99e-5
0.0001
0.959
0.969
Relative fitness of plasmid ω
Relative fitness of plasmid ω = 1
Equilibrium plasmid density
0
1
2
3
4

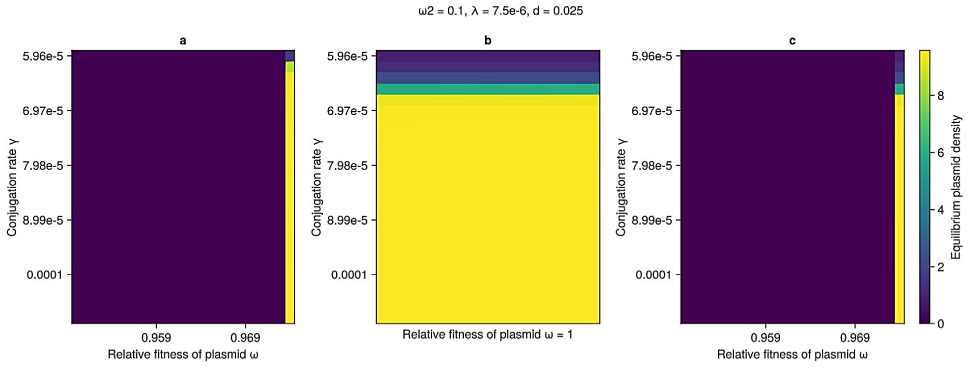
ω2 = 0.1, λ = 7.5e-6, d = 0.025
a
b
c
Conjugation rate γ
5.96e-5
6.97e-5
7.98e-5
8.99e-5
0.0001
0.959
0.969
Relative fitness of plasmid ω
Relative fitness of plasmid ω = 1
Equilibrium plasmid density
0
2
4
6
8

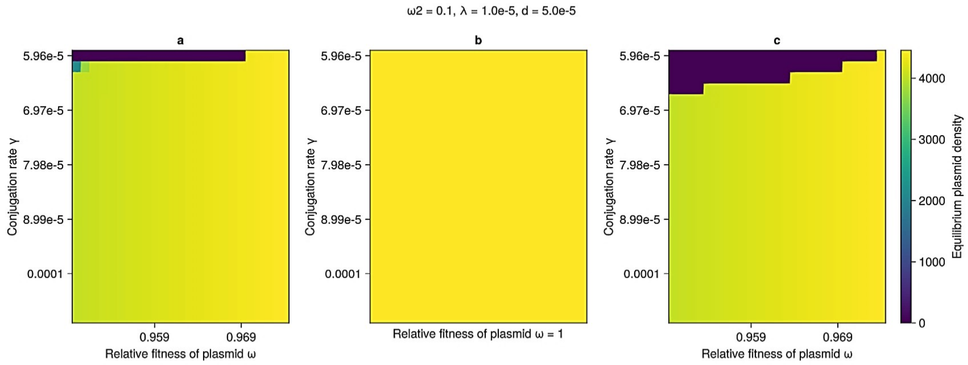
ω2 = 0.1, λ = 1.0e-5, d = 5.0e-5
a
b
c
Conjugation rate γ
5.96e-5
6.97e-5
7.98e-5
8.99e-5
0.0001
0.959
0.969
Relative fitness of plasmid ω
Relative fitness of plasmid ω = 1
Equilibrium plasmid density
0
1000
2000
3000
4000

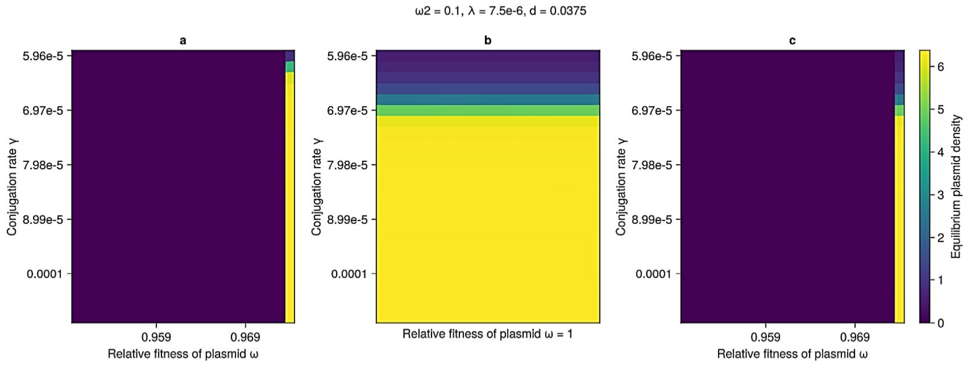
ω2 = 0.1, λ = 7.5e-6, d = 0.0375
a
b
c
Conjugation rate γ
5.96e-5
6.97e-5
7.98e-5
8.99e-5
0.0001
0.959
0.969
Relative fitness of plasmid ω
Relative fitness of plasmid ω = 1
Equilibrium plasmid density
0
1
2
3
4
5
6

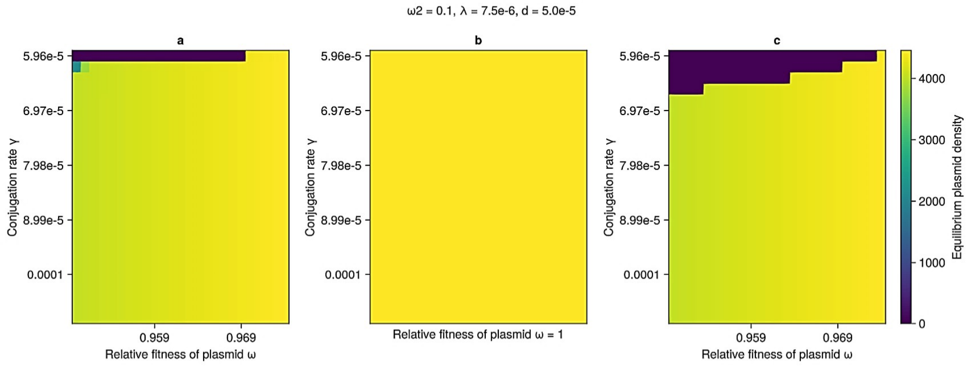
ω2 = 0.1, λ = 7.5e-6, d = 5.0e-5
a
b
c
5.96e-5
6.97e-5
7.98e-5
8.99e-5
0.0001
Conjugation rate γ
0.959
0.969
Relative fitness of plasmid ω
Relative fitness of plasmid ω = 1
0
1000
2000
3000
4000
Equilibrium plasmid density

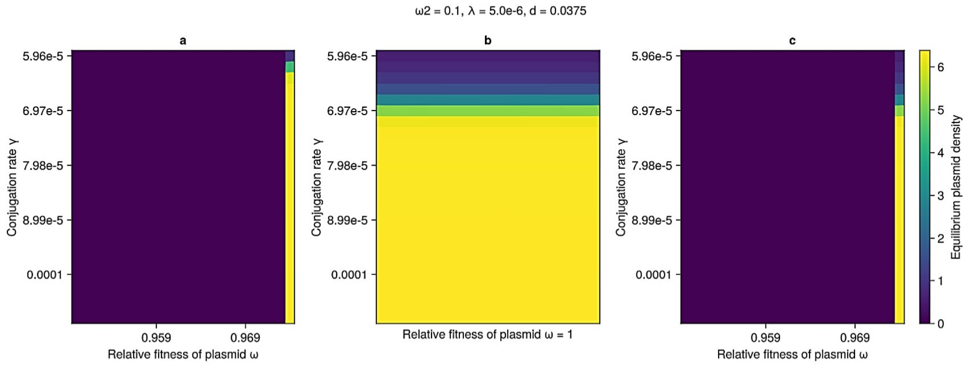
ω2 = 0.1, λ = 5.0e-6, d = 0.0375
a
b
c
5.96e-5
6.97e-5
7.98e-5
8.99e-5
0.0001
Conjugation rate γ
0.959
0.969
Relative fitness of plasmid ω
Relative fitness of plasmid ω = 1
0
1
2
3
4
5
6
Equilibrium plasmid density

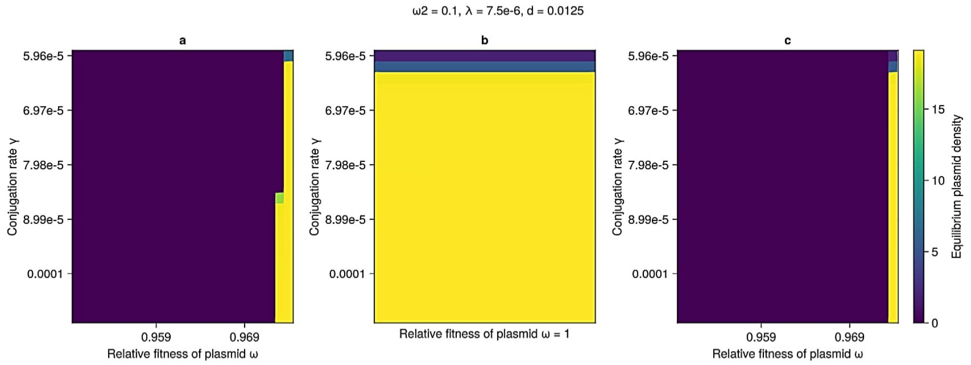
ω2 = 0.1, λ = 7.5e-6, d = 0.0125
a
b
c
5.96e-5
6.97e-5
7.98e-5
8.99e-5
0.0001
Conjugation rate γ
0.959
0.969
Relative fitness of plasmid ω
Relative fitness of plasmid ω = 1
0
5
10
15
Equilibrium plasmid density

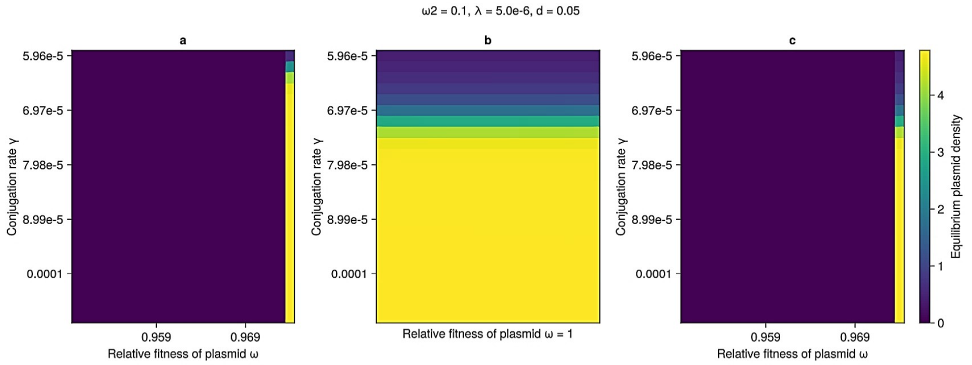
ω2 = 0.1, λ = 5.0e-6, d = 0.05
a
b
c
5.96e-5
6.97e-5
7.98e-5
8.99e-5
0.0001
Conjugation rate γ
0.959
0.969
Relative fitness of plasmid ω
Relative fitness of plasmid ω = 1
0
1
2
3
4
Equilibrium plasmid density

ω2 = 0.1, λ = 5.0e-6, d = 0.0125

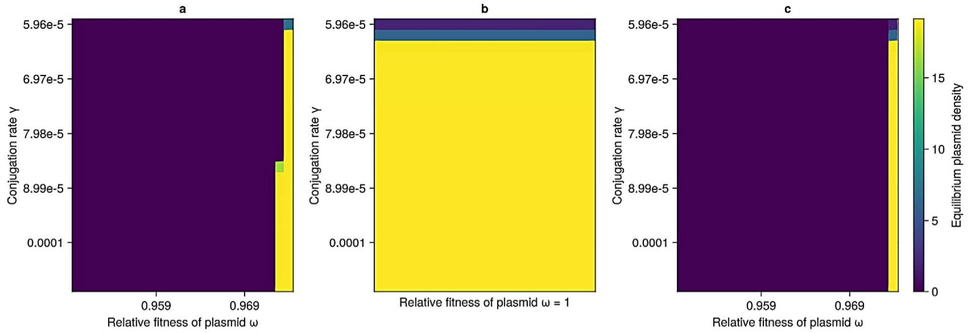


ω2 = 0.1, λ = 2.5e-6, d = 0.05

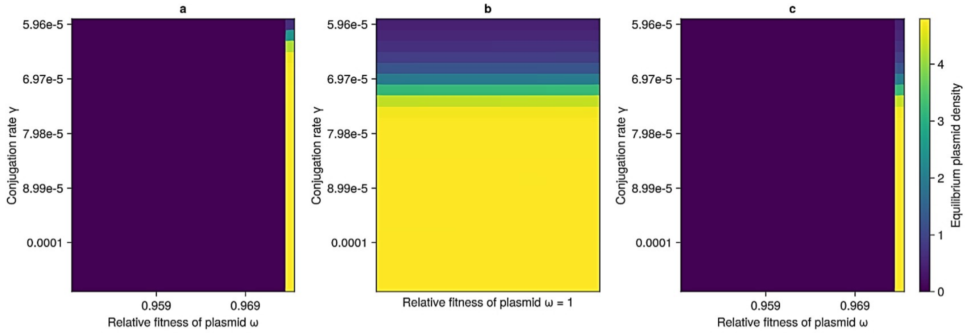


ω2 = 0.1, λ = 5.0e-6, d = 0.025

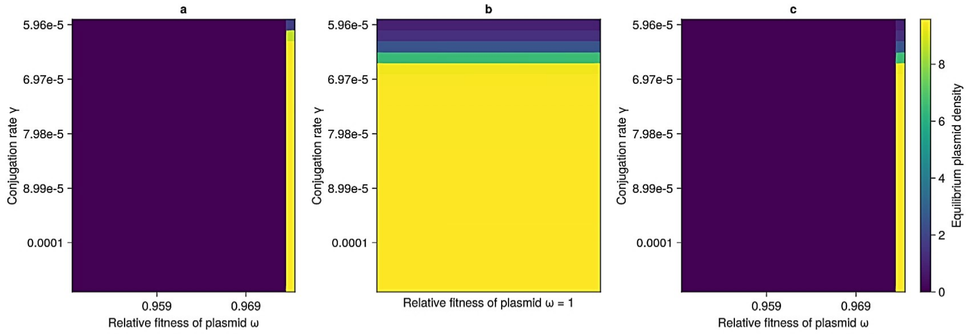


ω2 = 0.1, λ = 5.0e-6, d = 5.0e-5

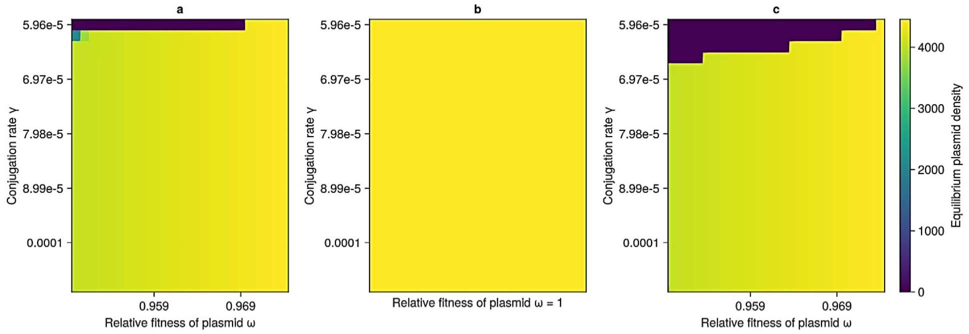

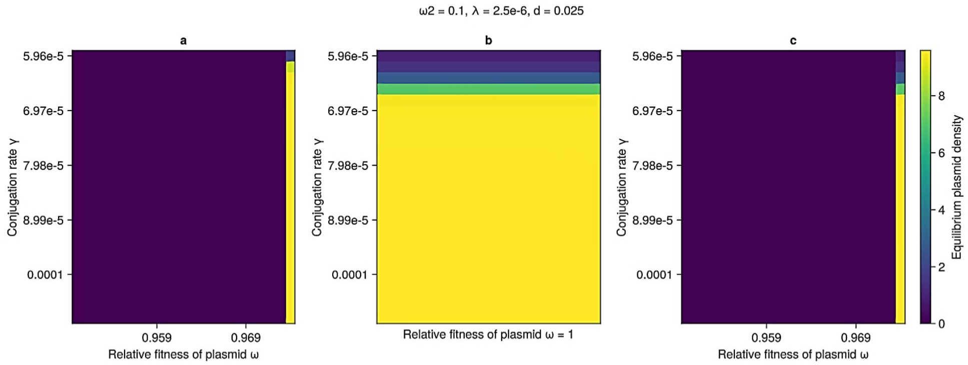
ω2 = 0.1, λ = 2.5e-6, d = 0.025
a
b
c
Conjugation rate γ
5.96e-5
6.97e-5
7.98e-5
8.99e-5
0.0001
0.959
0.969
Relative fitness of plasmid ω
Relative fitness of plasmid ω = 1
Equilibrium plasmid density
0
2
4
6
8

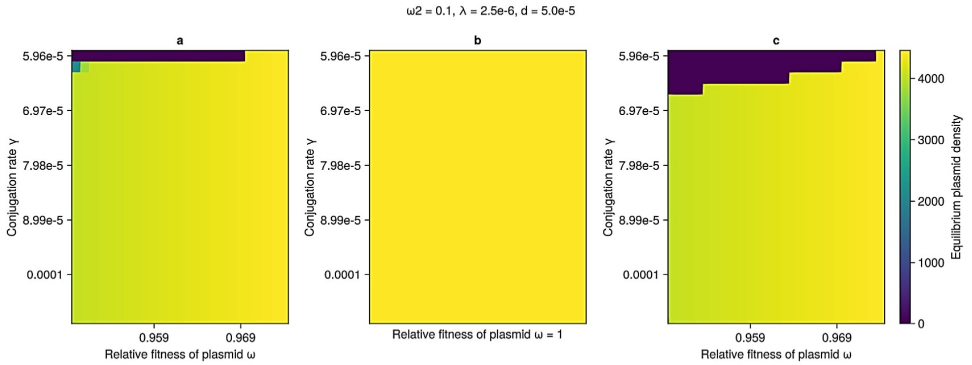
ω2 = 0.1, λ = 2.5e-6, d = 5.0e-5
a
b
c
Conjugation rate γ
5.96e-5
6.97e-5
7.98e-5
8.99e-5
0.0001
0.959
0.969
Relative fitness of plasmid ω
Relative fitness of plasmid ω = 1
Equilibrium plasmid density
0
1000
2000
3000
4000

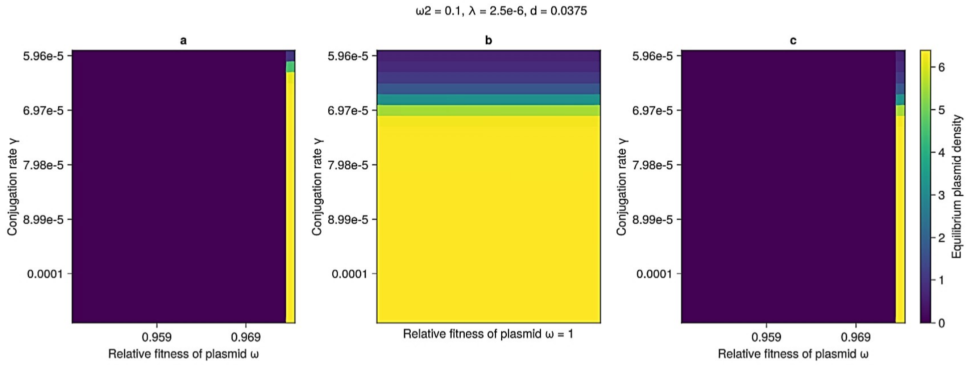
ω2 = 0.1, λ = 2.5e-6, d = 0.0375
a
b
c
Conjugation rate γ
5.96e-5
6.97e-5
7.98e-5
8.99e-5
0.0001
0.959
0.969
Relative fitness of plasmid ω
Relative fitness of plasmid ω = 1
Equilibrium plasmid density
0
1
2
3
4
5
6

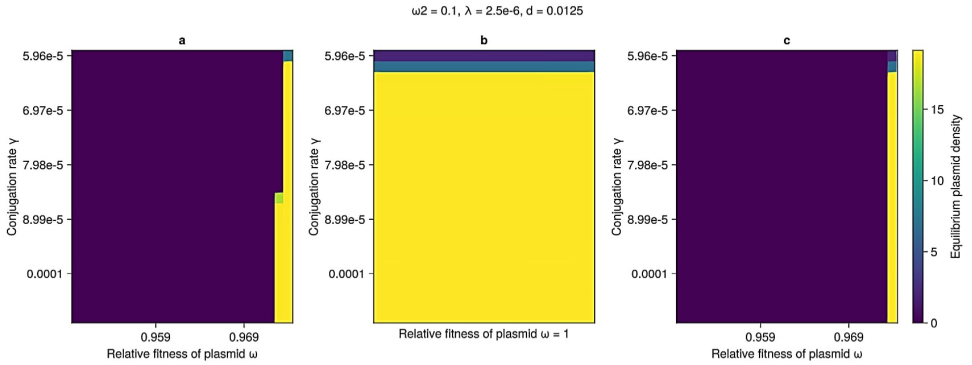
ω2 = 0.1, λ = 2.5e-6, d = 0.0125
a
b
c
Conjugation rate γ
5.96e-5
6.97e-5
7.98e-5
8.99e-5
0.0001
0.959
0.969
Relative fitness of plasmid ω
Relative fitness of plasmid ω = 1
Equilibrium plasmid density
0
5
10
15

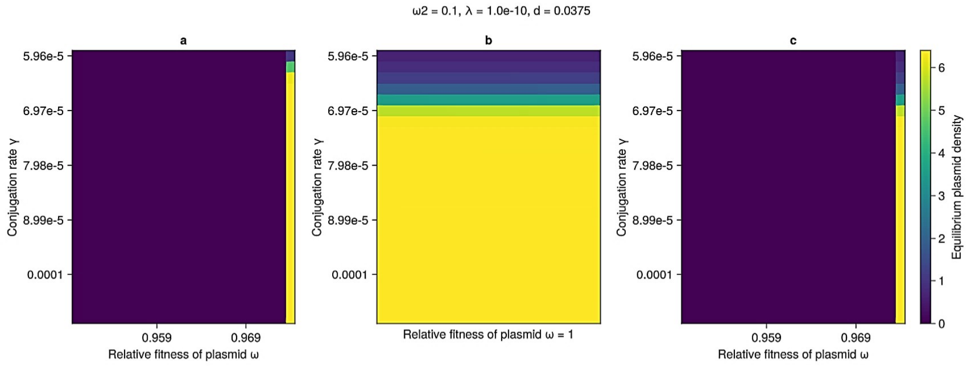
ω2 = 0.1, λ = 1.0e-10, d = 0.0375
a
b
c
Conjugation rate γ
5.96e-5
6.97e-5
7.98e-5
8.99e-5
0.0001
0.959
0.969
Relative fitness of plasmid ω
Relative fitness of plasmid ω = 1
Equilibrium plasmid density
0
1
2
3
4
5
6

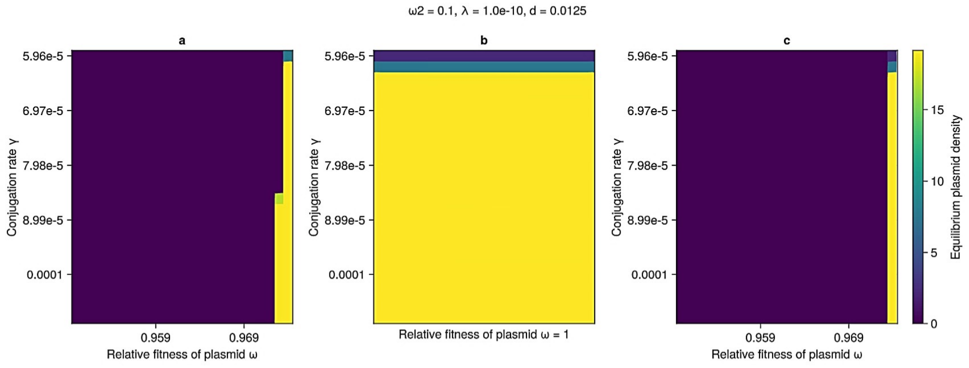
ω2 = 0.1, λ = 1.0e-10, d = 0.0125
a
b
c
Conjugation rate γ
5.96e-5
6.97e-5
7.98e-5
8.99e-5
0.0001
0.959
0.969
Relative fitness of plasmid ω
Relative fitness of plasmid ω = 1
Equilibrium plasmid density
0
5
10
15

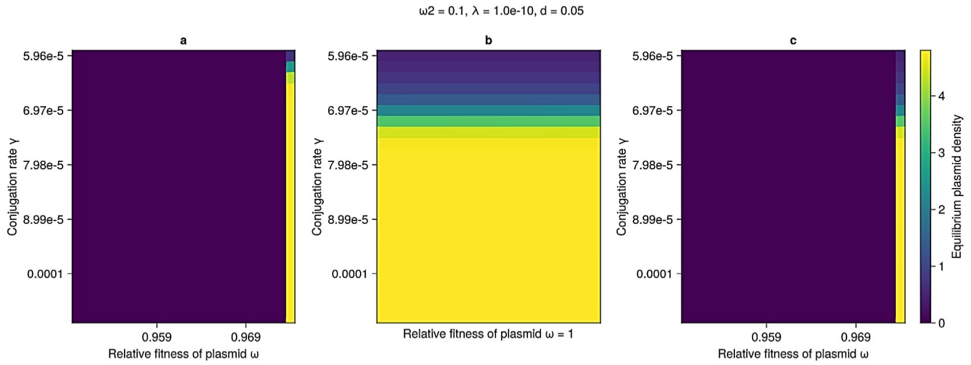
ω2 = 0.1, λ = 1.0e-10, d = 0.05
a
b
c
Conjugation rate γ
5.96e-5
6.97e-5
7.98e-5
8.99e-5
0.0001
0.959
0.969
Relative fitness of plasmid ω
Relative fitness of plasmid ω = 1
Equilibrium plasmid density
0
1
2
3
4

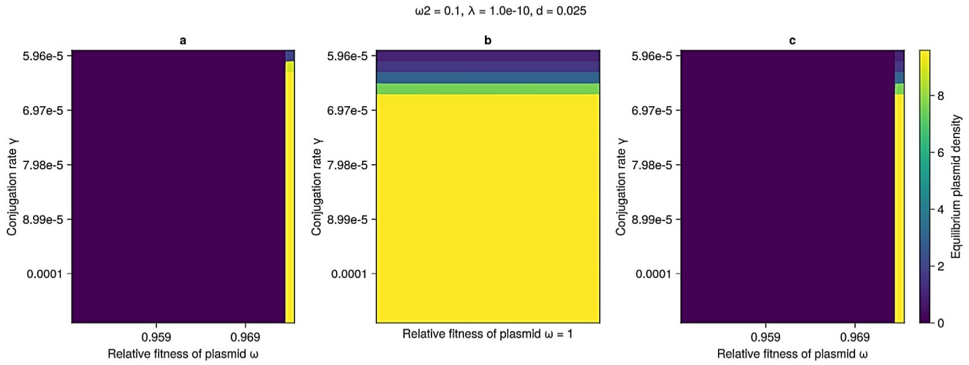
ω2 = 0.1, λ = 1.0e-10, d = 0.025
a
b
c
Conjugation rate γ
5.96e-5
6.97e-5
7.98e-5
8.99e-5
0.0001
0.959
0.969
Relative fitness of plasmid ω
Relative fitness of plasmid ω = 1
Equilibrium plasmid density
0
2
4
6
8

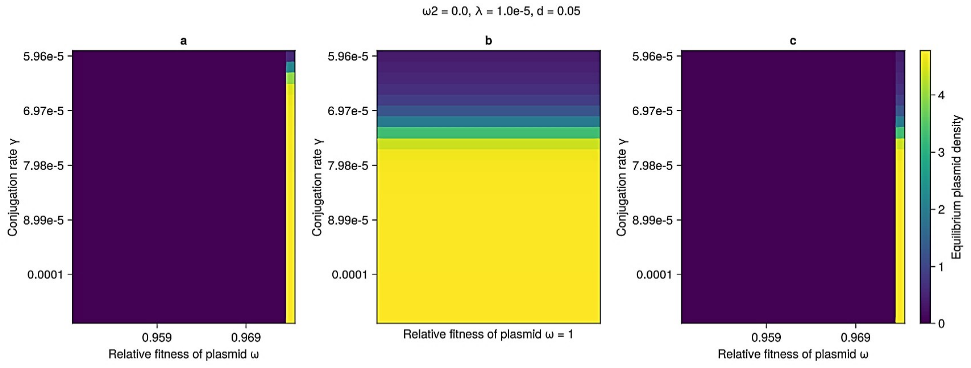
ω2 = 0.0, λ = 1.0e-5, d = 0.05
a
b
c
5.96e-5
6.97e-5
7.98e-5
8.99e-5
0.0001
Conjugation rate γ
0.959
0.969
Relative fitness of plasmid ω
Relative fitness of plasmid ω = 1
Equilibrium plasmid density
0
1
2
3
4

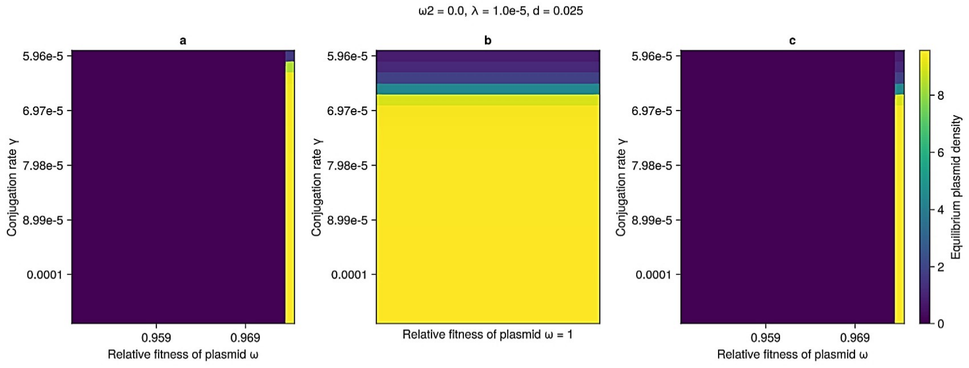
ω2 = 0.0, λ = 1.0e-5, d = 0.025
a
b
c
5.96e-5
6.97e-5
7.98e-5
8.99e-5
0.0001
Conjugation rate γ
0.959
0.969
Relative fitness of plasmid ω
Relative fitness of plasmid ω = 1
Equilibrium plasmid density
0
2
4
6
8

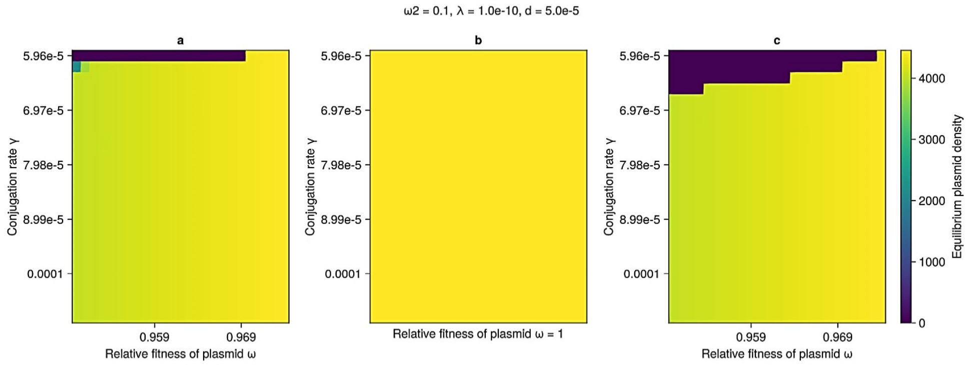
ω2 = 0.1, λ = 1.0e-10, d = 5.0e-5
a
b
c
5.96e-5
6.97e-5
7.98e-5
8.99e-5
0.0001
Conjugation rate γ
0.959
0.969
Relative fitness of plasmid ω
Relative fitness of plasmid ω = 1
Equilibrium plasmid density
0
1000
2000
3000
4000

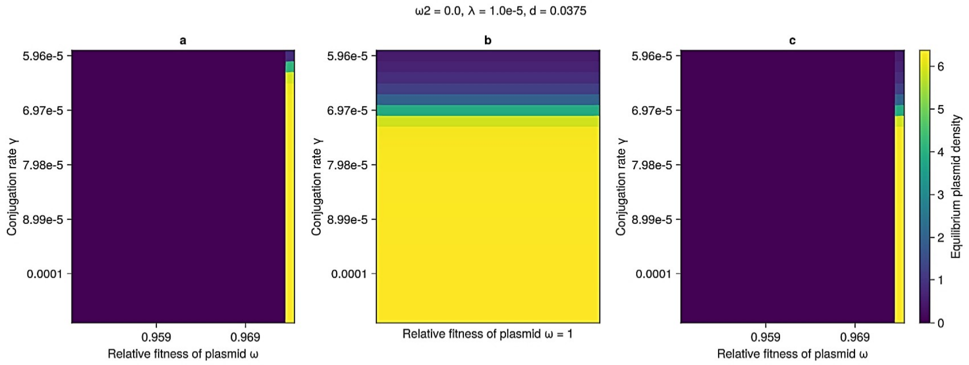
ω2 = 0.0, λ = 1.0e-5, d = 0.0375
a
b
c
5.96e-5
6.97e-5
7.98e-5
8.99e-5
0.0001
Conjugation rate γ
0.959
0.969
Relative fitness of plasmid ω
Relative fitness of plasmid ω = 1
Equilibrium plasmid density
0
1
2
3
4
5
6

ω2 = 0.0, λ = 1.0e-5, d = 5.0e-5

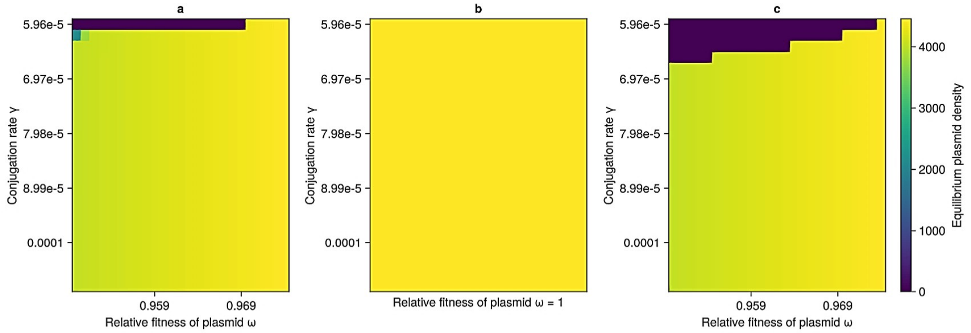


ω2 = 0.0, λ = 7.5e-6, d = 0.0375

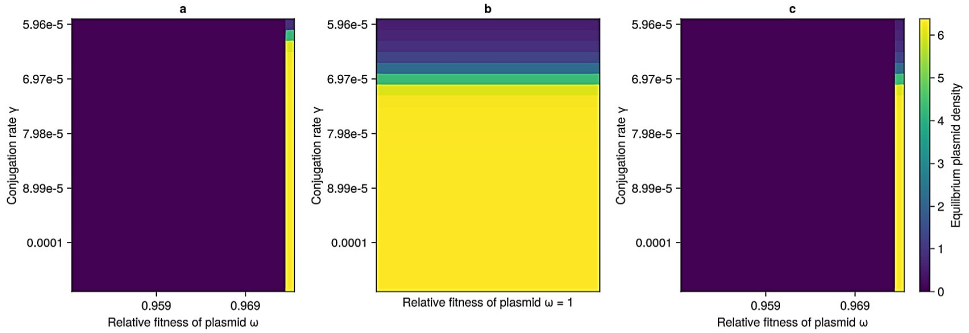


ω2 = 0.0, λ = 1.0e-5, d = 0.0125

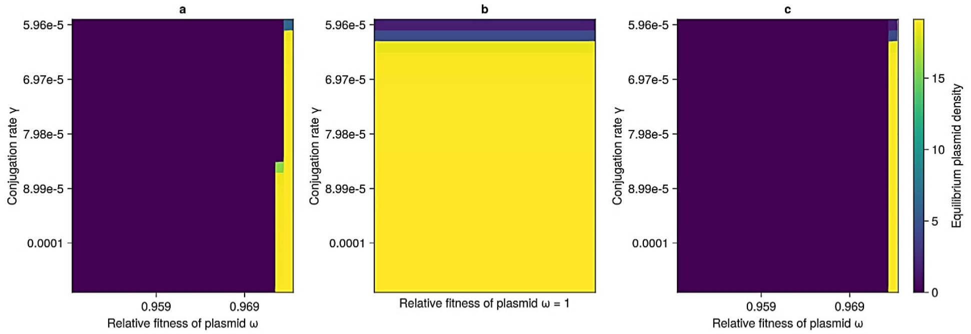


ω2 = 0.0, λ = 7.5e-6, d = 0.05

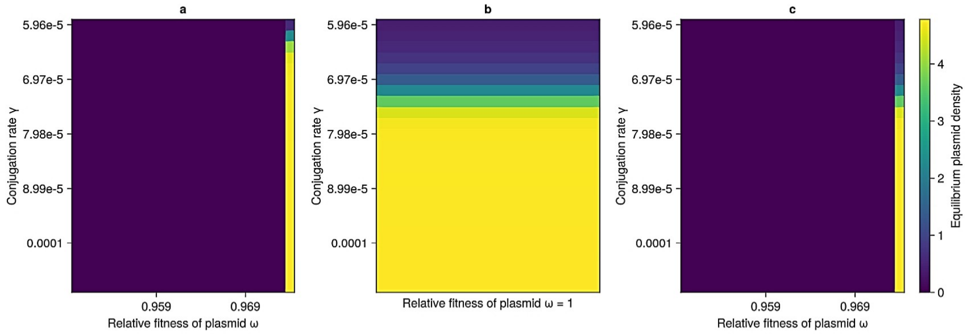

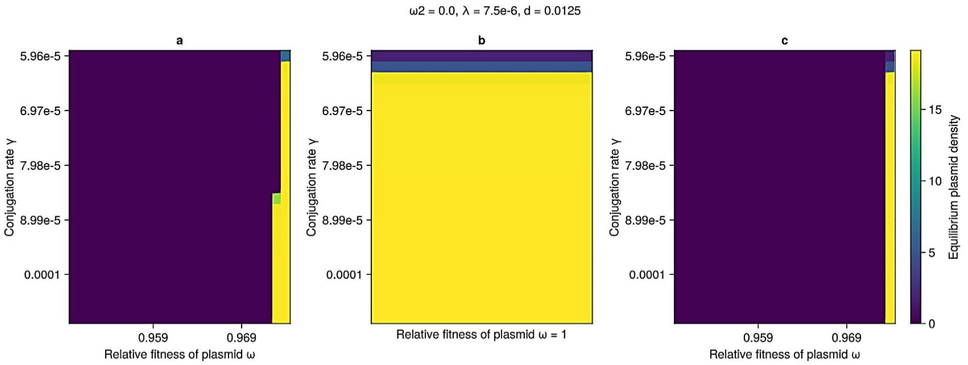
ω2 = 0.0, λ = 7.5e-6, d = 0.0125
a
b
c
Conjugation rate γ
5.96e-5
6.97e-5
7.98e-5
8.99e-5
0.0001
0.959
0.969
Relative fitness of plasmid ω
Relative fitness of plasmid ω = 1
Equilibrium plasmid density
0
5
10
15

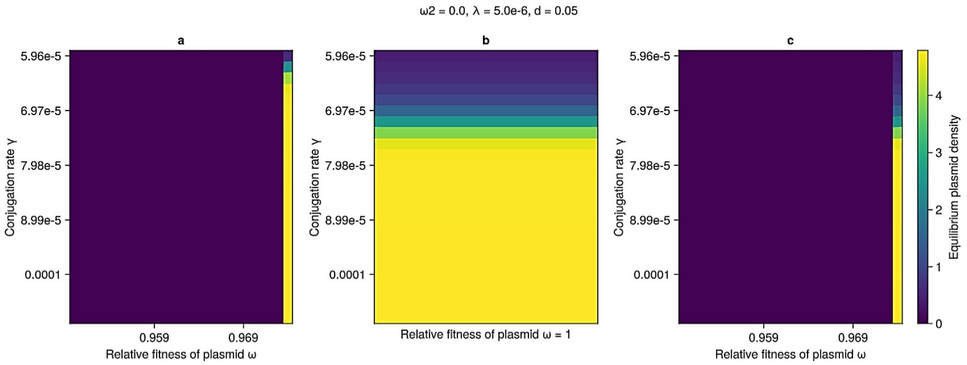
ω2 = 0.0, λ = 5.0e-6, d = 0.05
a
b
c
Conjugation rate γ
5.96e-5
6.97e-5
7.98e-5
8.99e-5
0.0001
0.959
0.969
Relative fitness of plasmid ω
Relative fitness of plasmid ω = 1
Equilibrium plasmid density
0
1
2
3
4

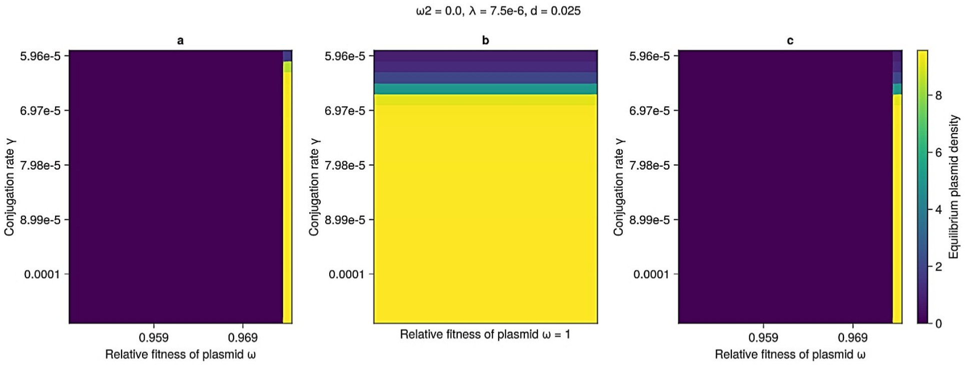
ω2 = 0.0, λ = 7.5e-6, d = 0.025
a
b
c
Conjugation rate γ
5.96e-5
6.97e-5
7.98e-5
8.99e-5
0.0001
0.959
0.969
Relative fitness of plasmid ω
Relative fitness of plasmid ω = 1
Equilibrium plasmid density
0
2
4
6
8

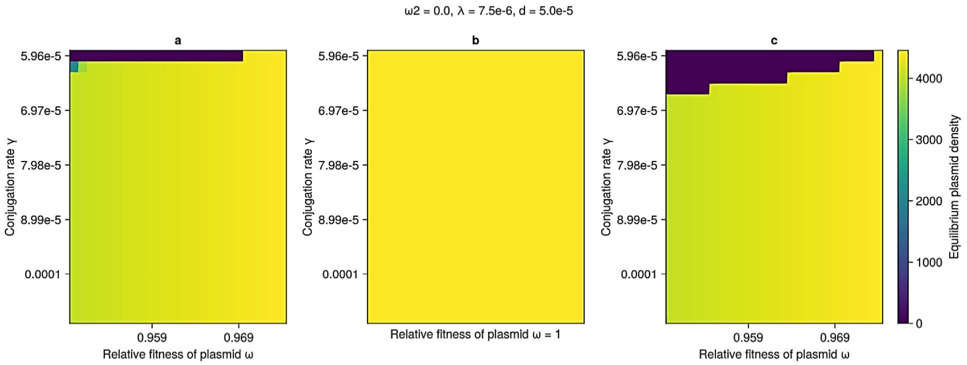
ω2 = 0.0, λ = 7.5e-6, d = 5.0e-5
a
b
c
Conjugation rate γ
5.96e-5
6.97e-5
7.98e-5
8.99e-5
0.0001
0.959
0.969
Relative fitness of plasmid ω
Relative fitness of plasmid ω = 1
Equilibrium plasmid density
0
1000
2000
3000
4000

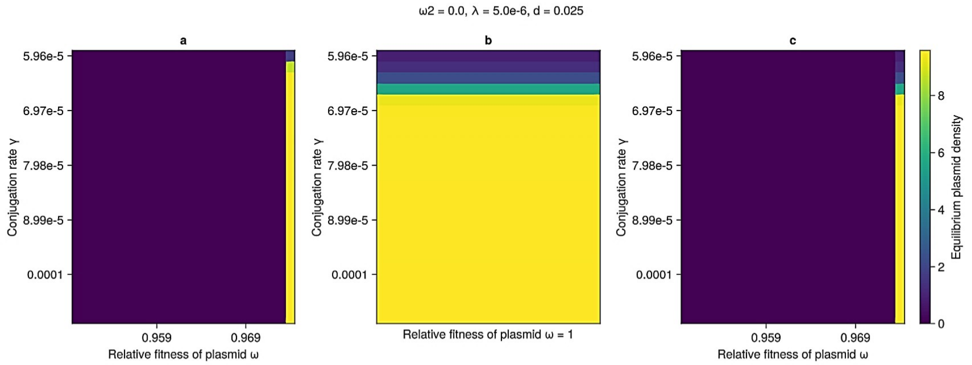
ω2 = 0.0, λ = 5.0e-6, d = 0.025
a
b
c
5.96e-5
6.97e-5
7.98e-5
8.99e-5
0.0001
Conjugation rate γ
0.959
0.969
Relative fitness of plasmid ω
Relative fitness of plasmid ω = 1
0
2
4
6
8
Equilibrium plasmid density

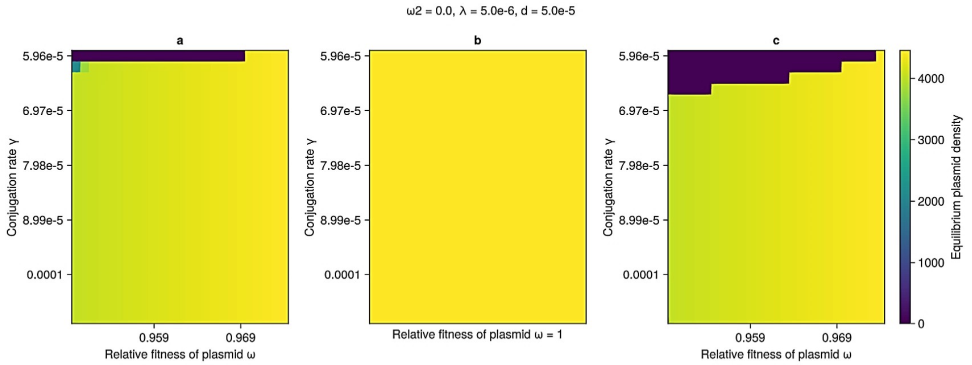
ω2 = 0.0, λ = 5.0e-6, d = 5.0e-5
a
b
c
5.96e-5
6.97e-5
7.98e-5
8.99e-5
0.0001
Conjugation rate γ
0.959
0.969
Relative fitness of plasmid ω
Relative fitness of plasmid ω = 1
0
1000
2000
3000
4000
Equilibrium plasmid density

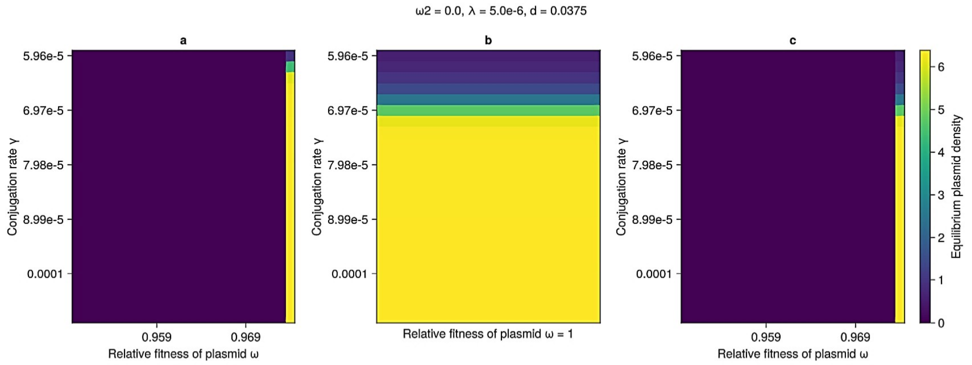
ω2 = 0.0, λ = 5.0e-6, d = 0.0375
a
b
c
5.96e-5
6.97e-5
7.98e-5
8.99e-5
0.0001
Conjugation rate γ
0.959
0.969
Relative fitness of plasmid ω
Relative fitness of plasmid ω = 1
0
1
2
3
4
5
6
Equilibrium plasmid density

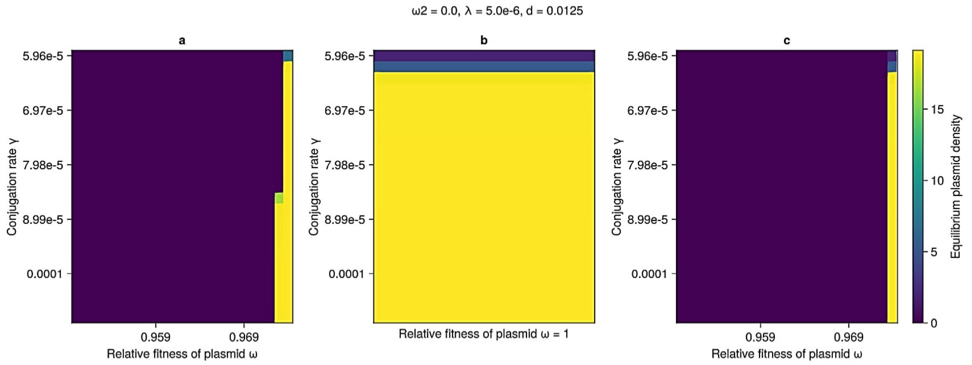
ω2 = 0.0, λ = 5.0e-6, d = 0.0125
a
b
c
5.96e-5
6.97e-5
7.98e-5
8.99e-5
0.0001
Conjugation rate γ
0.959
0.969
Relative fitness of plasmid ω
Relative fitness of plasmid ω = 1
0
5
10
15
Equilibrium plasmid density

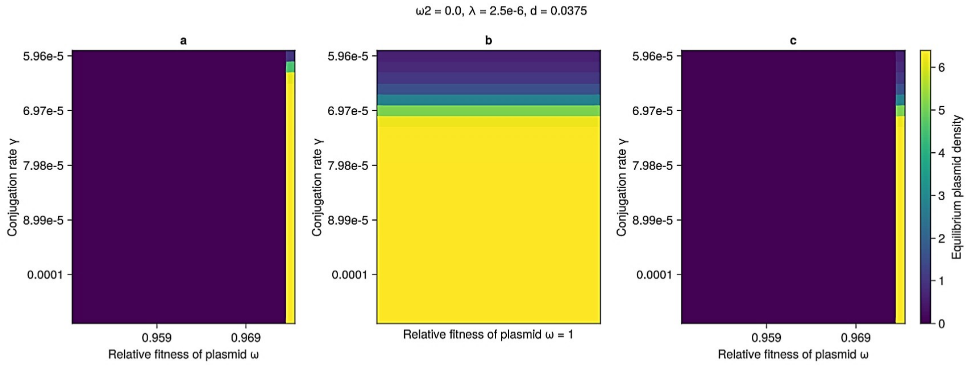
ω2 = 0.0, λ = 2.5e-6, d = 0.0375
a
b
c
Conjugation rate γ
5.96e-5
6.97e-5
7.98e-5
8.99e-5
0.0001
0.959
0.969
Relative fitness of plasmid ω
Relative fitness of plasmid ω = 1
Equilibrium plasmid density
0
1
2
3
4
5
6

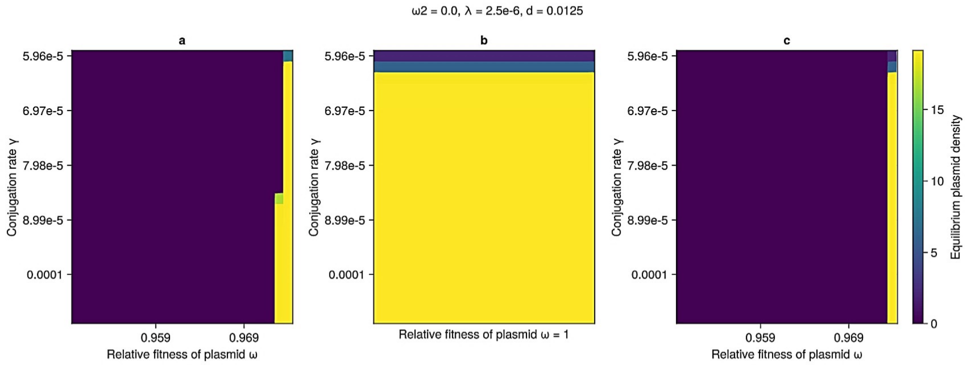
ω2 = 0.0, λ = 2.5e-6, d = 0.0125
a
b
c
Conjugation rate γ
5.96e-5
6.97e-5
7.98e-5
8.99e-5
0.0001
0.959
0.969
Relative fitness of plasmid ω
Relative fitness of plasmid ω = 1
Equilibrium plasmid density
0
5
10
15

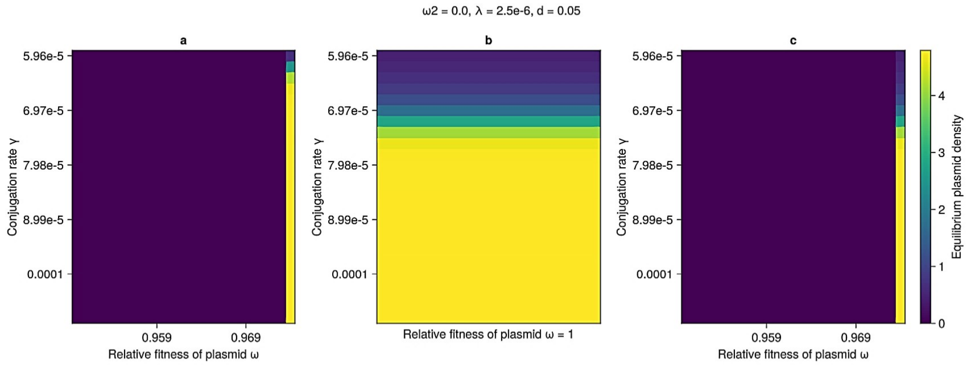
ω2 = 0.0, λ = 2.5e-6, d = 0.05
a
b
c
Conjugation rate γ
5.96e-5
6.97e-5
7.98e-5
8.99e-5
0.0001
0.959
0.969
Relative fitness of plasmid ω
Relative fitness of plasmid ω = 1
Equilibrium plasmid density
0
1
2
3
4

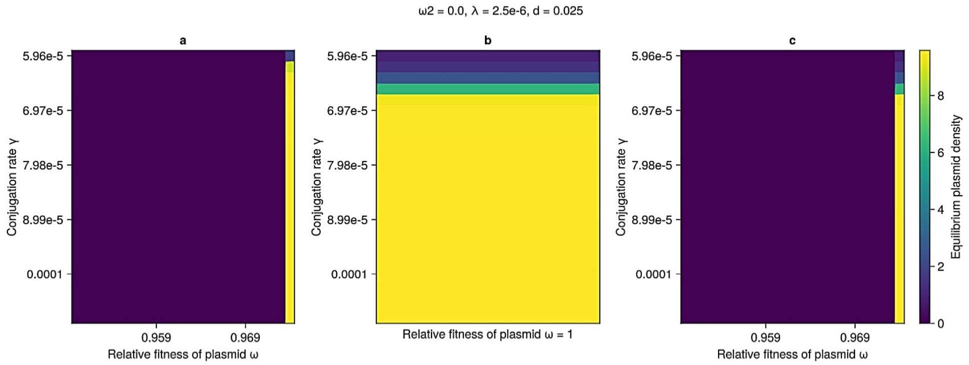
ω2 = 0.0, λ = 2.5e-6, d = 0.025
a
b
c
Conjugation rate γ
5.96e-5
6.97e-5
7.98e-5
8.99e-5
0.0001
0.959
0.969
Relative fitness of plasmid ω
Relative fitness of plasmid ω = 1
Equilibrium plasmid density
0
2
4
6
8

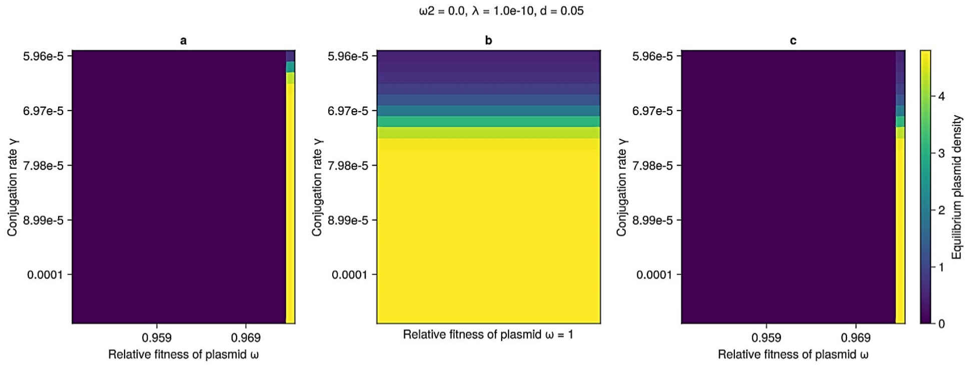
ω2 = 0.0, λ = 1.0e-10, d = 0.05
a
b
c
Conjugation rate γ
5.96e-5
6.97e-5
7.98e-5
8.99e-5
0.0001
0.959
0.969
Relative fitness of plasmid ω
Relative fitness of plasmid ω = 1
Equilibrium plasmid density
0
1
2
3
4

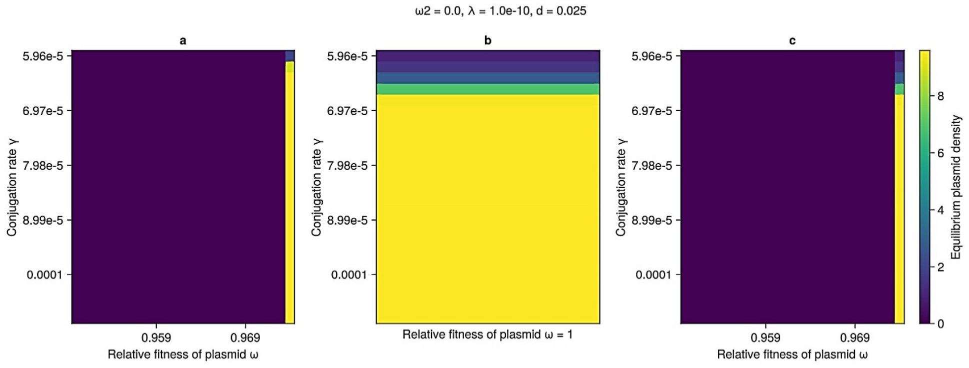
ω2 = 0.0, λ = 1.0e-10, d = 0.025
a
b
c
Conjugation rate γ
5.96e-5
6.97e-5
7.98e-5
8.99e-5
0.0001
0.959
0.969
Relative fitness of plasmid ω
Relative fitness of plasmid ω = 1
Equilibrium plasmid density
0
2
4
6
8

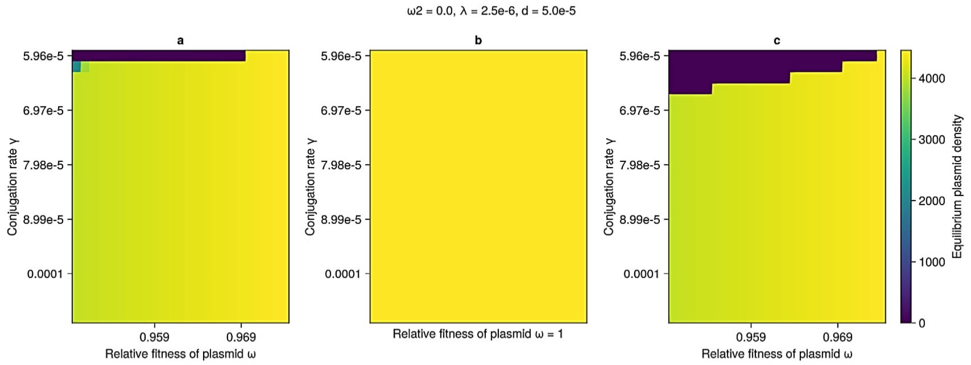
ω2 = 0.0, λ = 2.5e-6, d = 5.0e-5
a
b
c
Conjugation rate γ
5.96e-5
6.97e-5
7.98e-5
8.99e-5
0.0001
0.959
0.969
Relative fitness of plasmid ω
Relative fitness of plasmid ω = 1
Equilibrium plasmid density
0
1000
2000
3000
4000

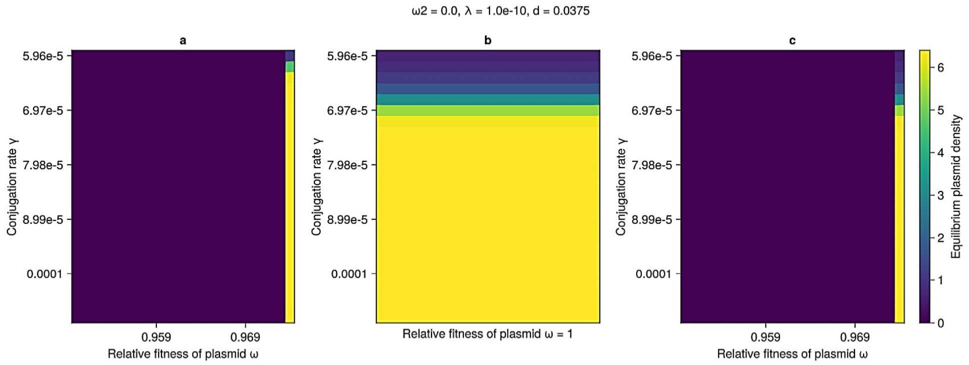
ω2 = 0.0, λ = 1.0e-10, d = 0.0375
a
b
c
Conjugation rate γ
5.96e-5
6.97e-5
7.98e-5
8.99e-5
0.0001
0.959
0.969
Relative fitness of plasmid ω
Relative fitness of plasmid ω = 1
Equilibrium plasmid density
0
1
2
3
4
5
6

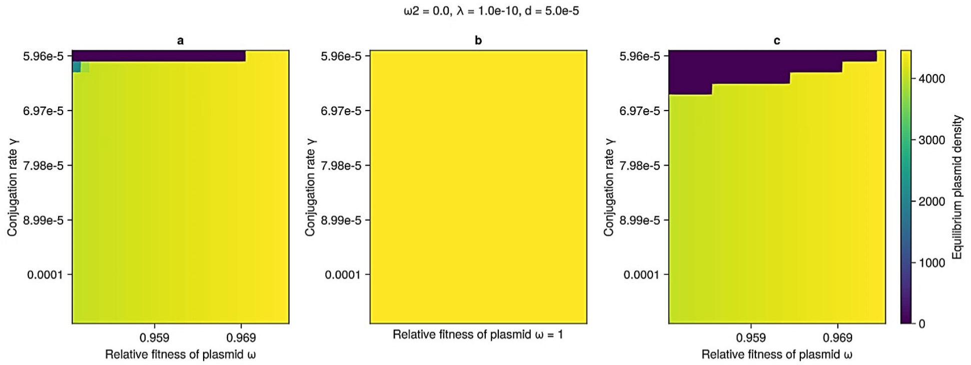
ω2 = 0.0, λ = 1.0e-10, d = 5.0e-5
a
b
c
Conjugation rate γ
5.96e-5
6.97e-5
7.98e-5
8.99e-5
0.0001
0.959
0.969
Relative fitness of plasmid ω
Relative fitness of plasmid ω = 1
Equilibrium plasmid density
0
1000
2000
3000
4000

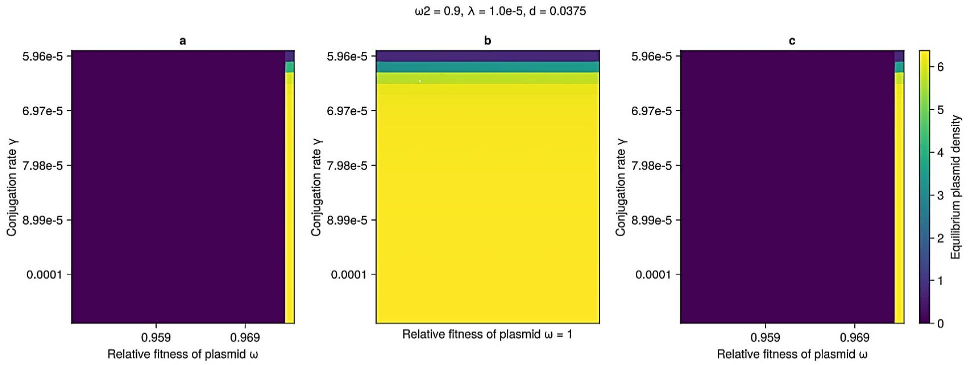
ω2 = 0.9, λ = 1.0e-5, d = 0.0375
a
b
c
Conjugation rate γ
5.96e-5
6.97e-5
7.98e-5
8.99e-5
0.0001
0.959
0.969
Relative fitness of plasmid ω
Relative fitness of plasmid ω = 1
Equilibrium plasmid density
0
1
2
3
4
5
6

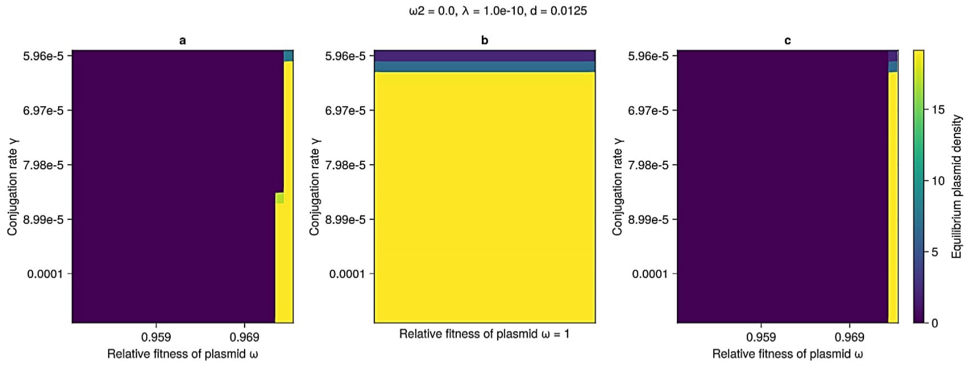
ω2 = 0.0, λ = 1.0e-10, d = 0.0125
a
b
c
Conjugation rate γ
5.96e-5
6.97e-5
7.98e-5
8.99e-5
0.0001
0.959
0.969
Relative fitness of plasmid ω
Relative fitness of plasmid ω = 1
Equilibrium plasmid density
0
5
10
15

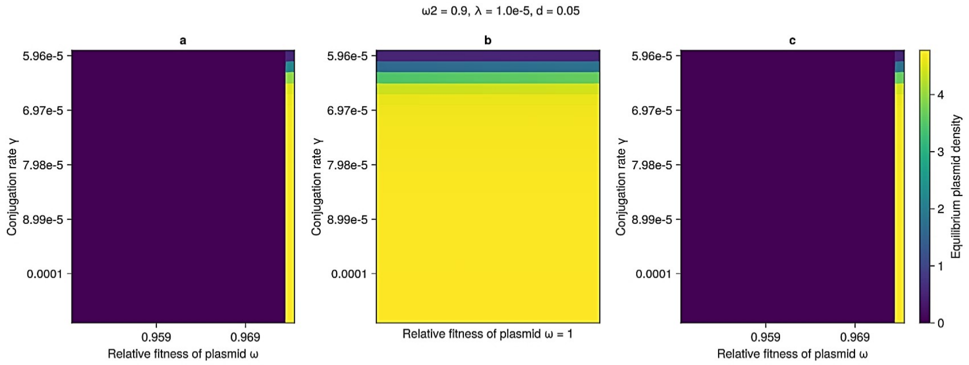
ω2 = 0.9, λ = 1.0e-5, d = 0.05
a
b
c
Conjugation rate γ
5.96e-5
6.97e-5
7.98e-5
8.99e-5
0.0001
0.959
0.969
Relative fitness of plasmid ω
Relative fitness of plasmid ω = 1
Equilibrium plasmid density
0
1
2
3
4

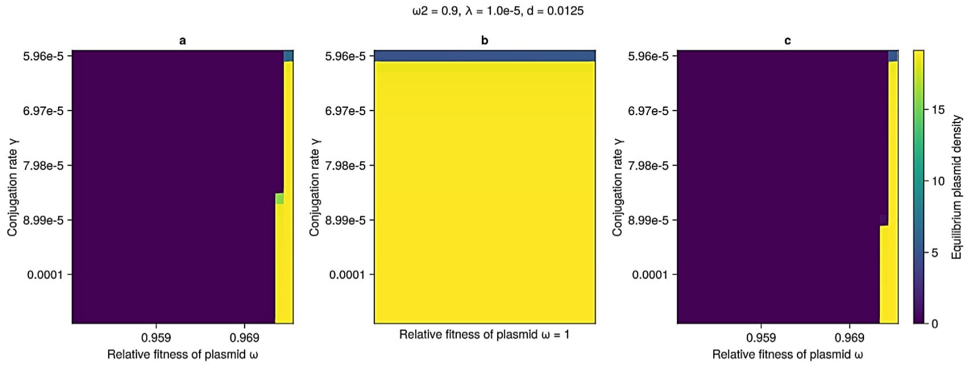
ω2 = 0.9, λ = 1.0e-5, d = 0.0125
a
b
c
Conjugation rate γ
5.96e-5
6.97e-5
7.98e-5
8.99e-5
0.0001
0.959
0.969
Relative fitness of plasmid ω
Relative fitness of plasmid ω = 1
Equilibrium plasmid density
0
5
10
15

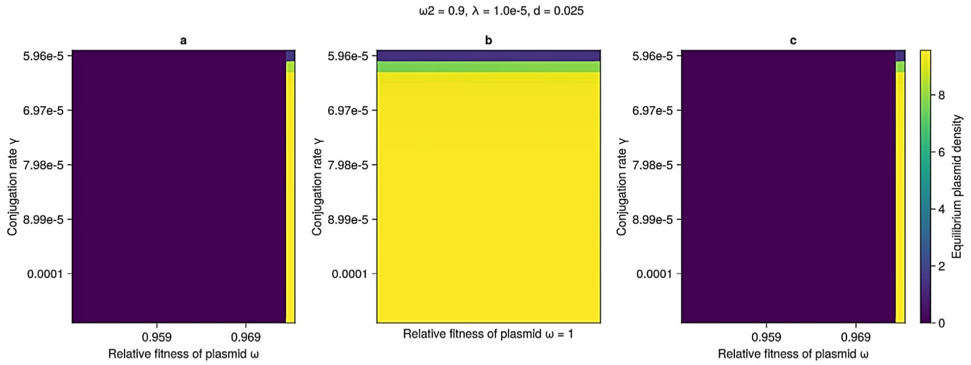
ω2 = 0.9, λ = 1.0e-5, d = 0.025
a
b
c
Conjugation rate γ
5.96e-5
6.97e-5
7.98e-5
8.99e-5
0.0001
0.959
0.969
Relative fitness of plasmid ω
Relative fitness of plasmid ω = 1
Equilibrium plasmid density
0
2
4
6
8

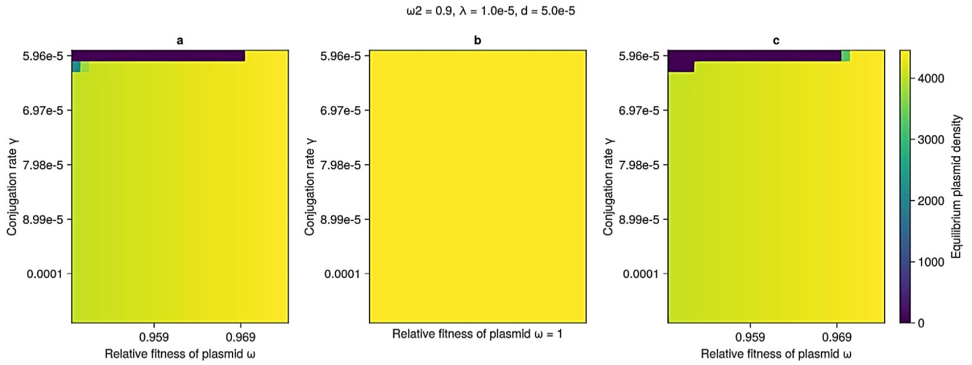
ω2 = 0.9, λ = 1.0e-5, d = 5.0e-5
a
b
c
Conjugation rate γ
5.96e-5
6.97e-5
7.98e-5
8.99e-5
0.0001
0.959
0.969
Relative fitness of plasmid ω
Relative fitness of plasmid ω = 1
Equilibrium plasmid density
0
1000
2000
3000
4000
